# Ribosome-tethered *in situ* sequencing for single-cell translatome analysis

**DOI:** 10.64898/2026.07.27.740970

**Authors:** Meng Jiang, Xiaojie He, Yanxiu Liu, Benchao Liu, Zhaoxiang Xie, Weiyan Ma, Lingyu Zhu, Chen Lin, Yongyou Zhang, Rongqin Ke

## Abstract

Translational regulation plays a critical role in shaping cellular states and functions, yet approaches for spatially resolved translatome profiling at single-cell resolution remain limited. Here, we develop Ribosome-tethered In Situ Sequencing (Ribo-ISS), an imaging-based spatial translatomics technology that enables high-throughput mapping of ribosome-associated mRNAs in intact tissues. Ribo-ISS integrates ribosome-dependent molecular anchoring with multiplexed in situ sequencing, allowing specific detection of translation-associated transcripts without genetic manipulation or exogenous ribosome labeling. We demonstrate that Ribo-ISS achieves high specificity and enables single-cell spatial translatome profiling in mouse brain, accurately recapitulating major cell types and their anatomical organization. Applied to a sleep deprivation model, Ribo-ISS simultaneously resolved transcriptional and translational changes, revealing extensive transcription-translation uncoupling and distinct cell-type-specific translational responses. Ribo-ISS provides a scalable framework for investigating spatially organized translational programs and expands the capability of spatial omics toward understanding gene regulation beyond transcription.

**Graphical Abstract:** 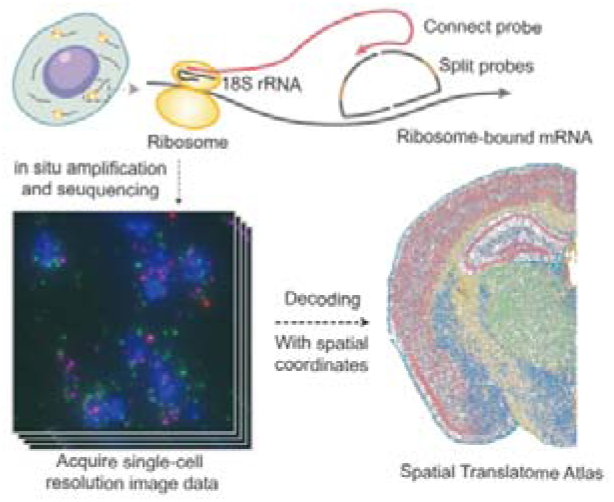

Ribo-ISS integrates ribosome-dependent molecular anchoring with multiplexed in situ sequencing, enabling specific detection of ribosome-bound transcripts in intact tissues without genetic manipulation or exogenous labeling. As an imaging-based spatial translatomics technology, it resolves translatome profiling at single-cell resolution. This scalable approach reveals spatial translational programs, extending spatial omics beyond transcription.

## Introduction

Within the framework of the central dogma, the flow of genetic information is often simplified as a linear process from DNA to RNA to protein. However, gene expression is governed by multilayered and dynamic regulatory mechanisms that extend far beyond transcription. These mechanisms operate at four major stages: transcriptional regulation, post-transcriptional regulation, translational regulation, and post-translational regulation, collectively enabling cells to precisely control protein production and rapidly adapt to changing environments^[1–5]^. Quantitative proteomic studies in mouse fibroblasts have demonstrated that protein abundance is strongly influenced by translational regulation, with translational control making a substantial contribution to proteome variability beyond transcriptional regulation alone^[6]^. Therefore, translation, during which ribosomes decode messenger RNAs (mRNAs) to synthesize proteins, represents a critical regulatory layer in the transmission of genetic information.

Despite its importance, most studies of gene expression have traditionally relied on transcriptome sequencing technologies, which measure mRNA abundance but often provide an incomplete representation of protein production^[7–8]^. This discrepancy arises because individual transcripts exhibit extensive heterogeneity in translational efficiency: some mRNAs are actively engaged with ribosomes and efficiently translated, whereas others undergo translational repression or storage^[9]^. Through dynamic control of translation, cells can rapidly and efficiently remodel their proteomes without altering transcript abundance, thereby adapting to environmental changes and responding promptly to endogenous and exogenous signals^[10–12]^. Thus, capturing translational regulation is essential for a comprehensive understanding of gene expression programs, yet remains challenging with conventional transcriptomic approaches.

The central role of translational regulation in gene expression has driven the continuous development of translatomics technologies, progressing from biochemical fractionation-based approaches to high-throughput sequencing and, more recently, to single-cell and spatially resolved measurements. Early approaches such as polysome profiling employed sucrose density gradient centrifugation to separate mRNAs associated with different numbers of ribosomes, enabling the assessment of ribosome occupancy and relative translational activity of transcripts^[13]^. However, this approach is labor-intensive, requires extensive sample processing, and suffers from substantial material loss during fraction collection, limiting its application for comprehensive transcriptome-wide analysis^[14]^.

Subsequent strategies aimed to directly characterize ribosome-mRNA complexes. Ribosome nascent-chain complex sequencing (RNC-seq) was developed to isolate ribosome-associated mRNAs and determine their transcript identities through high-throughput sequencing^[15]^. However, because ribosome-mRNA associations are mediated through noncovalent interactions, complex disruption during purification and subsequent mRNA degradation can introduce quantitative biases. To overcome the limitation of bulk-level measurements and enable cell type-specific translatome profiling in heterogeneous tissues, approaches such as translating ribosome affinity purification (TRAP)^[16–17]^ and RiboTag^[18]^ introduced affinity-tagged ribosomal proteins into defined cell populations, allowing selective isolation of actively translated mRNAs. Nevertheless, these approaches require tissue homogenization, resulting in the loss of spatial context and preventing the direct visualization of translational states within intact tissues.

A major breakthrough came with the development of ribosome profiling (Ribo-seq)^[19]^, in which RNase treatment is used to isolate ribosome-protected mRNA fragments (∼30 nucleotides), followed by deep sequencing to provide genome-wide maps of ribosome occupancy and translation activity. Since its introduction, numerous Ribo-seq derivatives have been developed to investigate specific aspects of translation, including Disome-seq^[20–21]^ for detecting ribosome collisions and Ribo-taper^[22]^ for identifying translation regulation with enhanced precision. In parallel, metabolic labeling-based approaches, including BONCAT^[23]^, FUNCAT^[24]^, and SUnSET^[25]^, have enabled direct measurement of newly synthesized proteins and provided complementary insights into translational output.

Although these approaches have substantially advanced our understanding of translational regulation, most remain reliant on population-level measurements and therefore obscure cellular heterogeneity. This limitation has motivated the development of single-cell translatomic technologies. However, adapting ribosome profiling to the single-cell level remains technically challenging because of the extremely limited abundance of ribosome-protected fragments (RPFs), resulting in difficulties in sensitivity, quantitative accuracy, throughput, and broad sample applicability^[26]^. To overcome these limitations, Ribo-STAMP, developed by Brannan et al., introduced an alternative strategy that avoids direct sequencing of scarce RPFs by using RNA editing to mark ribosome-associated transcripts, thereby enabling integration with single-cell RNA sequencing workflows^[27]^. More recently, Sison et al. combined Ribo-STAMP with long-read single-cell sequencing to achieve cell-type-specific and isoform-resolved translatomic profiling^[28]^. Nevertheless, these approaches require genetic introduction of exogenous APOBEC fusion proteins, and editing efficiency-dependent variability may influence quantitative measurements. In addition, the dissociation-based sequencing workflows eliminate spatial information, preventing the investigation of translational regulation within intact tissue architectures. Conversely, imaging-based approaches for monitoring nascent protein synthesis, such as FUNCAT and FLARIM^[29]^, preserve spatial context but are limited in molecular throughput and are therefore not readily scalable for transcriptome-wide translatomic profiling. Thus, a technology capable of simultaneously achieving high-throughput, single-cell resolution, and spatially resolved translatomic profiling without genetic manipulation remains highly desirable.

To address this limitation, RIBOmap, the first spatially resolved ribosome-bound transcriptomic method, established the feasibility of mapping ribosome-associated mRNAs within intact tissues by combining ribosome labeling with in situ sequencing^[30]^. The success of RIBOmap highlights the potential of spatial translatomics, while complementary approaches based on alternative molecular anchoring strategies, simplified probe architectures, and alternative detection schemes may further broaden the applicability of this emerging field. Building upon our previously developed in situ sequencing framework based on split-probe ligation and rolling circle amplification^[31]^, we developed Ribosome-tethered In Situ Sequencing (Ribo-ISS), which employs ribosomal RNA-mediated molecular anchoring and a streamlined probe architecture to enable high-throughput profiling of ribosome-associated mRNAs while preserving native tissue architecture. Integration of molecular identities with cellular spatial coordinates allows Ribo-ISS to generate single-cell spatial translatomic maps, providing insights into cellular heterogeneity and spatial regulation of translation programs across complex biological systems.

## Results and Discussion

### Design and Principle of Ribo-ISS

Ribo-ISS is an imaging-based in situ sequencing technology developed for high-throughput spatial profiling of ribosome-associated mRNAs. The principle and workflow of Ribo-ISS are illustrated in Figure 1a. The key innovation of Ribo-ISS lies in a ribosome-dependent probe architecture that couples endogenous ribosomal RNA recognition with mRNA-targeted in situ sequencing. Specifically, Ribo-ISS employs a split-probe system consisting of two probes targeting the mRNA of interest and a connect probe that simultaneously recognizes 18S rRNA and the split probes. The 5’ region of the connect probe (25 nt) hybridizes to 18S rRNA, whereas its 3’ region (12 nt) interacts with the split probes bound to the target mRNA. When ribosomes are associated with the target transcript, the connect probe is stably anchored through rRNA recognition, enabling efficient ligation of the adjacent split probes. In contrast, in the absence of ribosome-mediated anchoring, the short interaction between the connect probe and split probes is insufficient to support efficient ligation. This design therefore selectively enriches signals from ribosome-associated transcripts while minimizing contributions from unbound RNA molecules.

**Figure 1.**
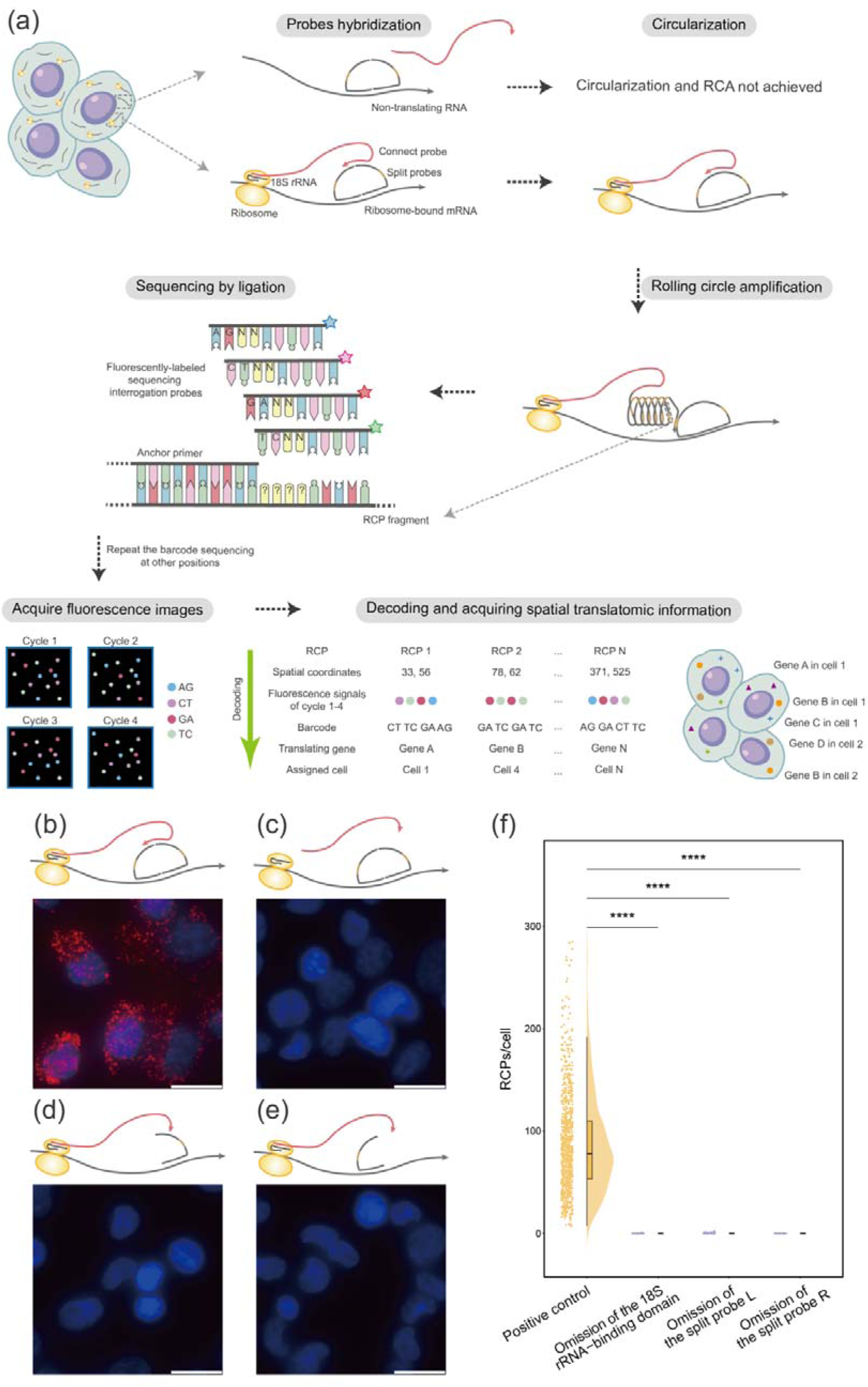
Design and implementation of Ribo-ISS. (a) Schematic diagram of Ribo-ISS. A ribosome-anchored connect probe bridges gene-specific split probes, enabling SplintR ligase-mediated circularization and rolling circle amplification; the resulting amplicons are decoded via improved combinatorial probe anchor ligation sequencing to map translating mRNAs at single-cell resolution. (b-e) Validation of probe functionality and specificity for detecting translating *ACTB* mRNA in H1299 cells. The complete probe set (b) generates a robust fluorescent signal. Control reactions omitting the 18S rRNA-binding domain (c) or either half of the split probe pair (d, e) all result in loss of signal. Scale bar, 20 μm. (f) Quantification of translating *ACTB* mRNA from the experiment shown in (b-e). Statistical significance was determined by Mann-Whitney U test, ****p < 0.0001.

Following ribosome-dependent probe assembly, the split probes carrying gene-specific barcodes are circularized by SplintR ligase, using the target mRNA and the connect probe as ligation templates. The circularized probes are subsequently amplified by rolling circle amplification (RCA), with the connect probe serving as the primer, generating localized rolling circle products (RCPs) containing gene-specific barcodes. These barcodes are decoded through our improved combinatorial probe anchor ligation-based in situ sequencing chemistry^[32]^, involving iterative cycles of fluorescent probe ligation, imaging, and signal removal. By integrating barcode decoding with image-based cell segmentation and spatial registration, Ribo-ISS enables the identification and spatial mapping of ribosome-associated mRNAs at single-cell resolution, thereby generating spatially resolved translatomic profiles within intact tissues.

### Validation of Ribo-ISS Performance

To validate the specificity and performance of Ribo-ISS, we first systematically evaluated the contribution of each probe component using actively translated *ACTB* mRNA in H1299 cells as a model target. Four experimental configurations were designed to interrogate the requirements for ribosome-dependent anchoring and target recognition (Figure 1b-e). The complete probe set containing both the connect probe and split probes generated robust *ACTB*-specific signals, with a median of 78 RCPs per cell (IQR, 53-110; n = 795), whereas removal of the 18S rRNA-binding region of the connect probe or omission of either split probe completely abolished signal generation (median = 0 RCPs per cell for all three negative controls; n = 975, 722 and 906, respectively). These results demonstrate that Ribo-ISS signal formation requires both ribosome-mediated anchoring through rRNA recognition and accurate dual recognition of the target transcript. Together, these controls establish the fundamental specificity of Ribo-ISS for detecting ribosome-associated mRNAs.

We next compared the detection performance of Ribo-ISS with RIBOmap using identical target regions in MCF-7 cells (Figure 2a). Actively translated *ACTB* mRNA and non-coding RNA *MALAT1* were selected as positive and negative controls, respectively, to evaluate detection efficiency and specificity. For *ACTB*, Ribo-ISS generated a comparable number of spatial signals to RIBOmap, with a median of 58 RCPs per cell (IQR, 44-75; n = 1,463), compared with 54 RCPs per cell (IQR, 38-77; n = 1,127) detected by RIBOmap (Figure 2b). Notably, Ribo-ISS exhibited substantially reduced background detection for *MALAT1*, with a median of 0 RCPs per cell (IQR, 0-0; n = 1,546), whereas RIBOmap generated a median of 4 RCPs per cell (IQR, 2-7; n = 1,091) (Figure 2c). These results demonstrate that Ribo-ISS achieves efficient detection of ribosome-associated transcripts while maintaining high specificity.

**Figure 2.**
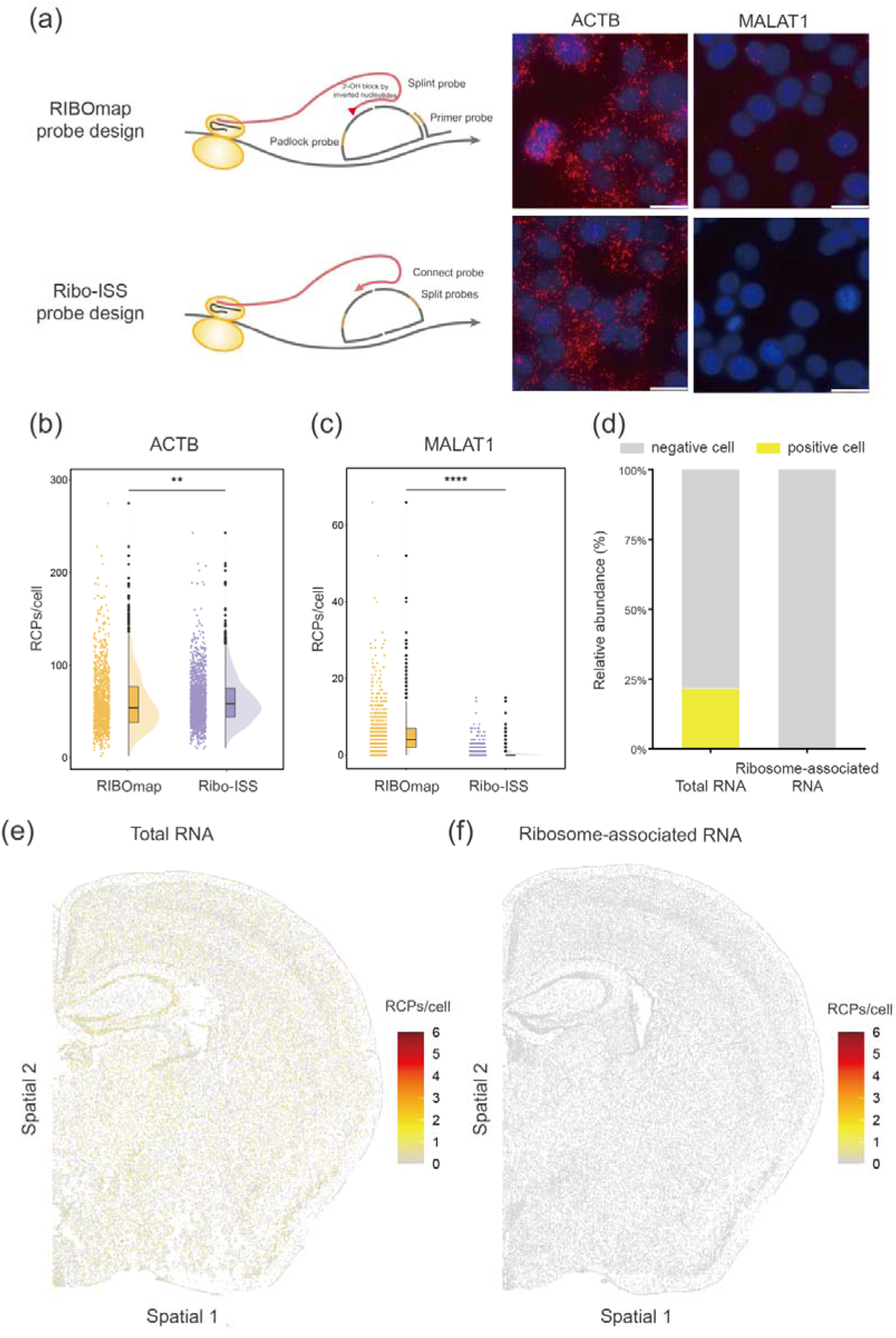
Validation of Ribo-ISS. (a) Probe design schematics and representative fluorescence images from comparative experiments between Ribo-ISS and RIBOmap, detecting *ACTB* and *MALAT1* in MCF-7 cells. Scale bar, 20 μm. (b and c) Quantification of *ACTB* (b) and *MALAT1* (c) from the experiment shown in (a), comparing the two technologies. Statistical significance was determined by Mann-Whitney U test, **p < 0.01, ****p < 0.0001. (d) Quantification of the proportion of *Malat1*-positive cells in mouse brain tissue at the ribosome-associated and total RNA levels.(e) Spatial expression of *Malat1* at the total RNA level in mouse brain tissue, detected by IISS. (f) Spatial expression of *Malat1* at the ribosome-associated RNA level in mouse brain tissue, detected by Ribo-ISS.

To further assess the ability of Ribo-ISS to distinguish transcriptional abundance from translation-associated signals, we analyzed *Malat1* expression at both RNA and ribosome-associated levels in mouse brain sections (Figure 2d-f). As expected, *Malat1* exhibited widespread transcriptional expression, with 10,504 of 48,456 cells showing detectable RNA signals (21.68%). In contrast, only 68 of 56,100 cells displayed ribosome-associated *Malat1* signals (0.12%) (Figure 2d). These results demonstrate that Ribo-ISS selectively captures ribosome-associated transcripts rather than total RNA abundance, enabling reliable discrimination between transcriptional and translational states.

In addition, we assessed whether Ribo-ISS can capture changes in ribosome-mRNA binding by treating cells with puromycin, a translation inhibitor that promotes premature termination of nascent polypeptide chains and ribosome dissociation from mRNA (Figure S1a-c). Ribosome-associated *ACTB* RNA was significantly reduced in puromycin-treated cells compared to untreated controls, with a median of 72 RCPs per cell (IQR, 50-102; n = 241) versus 208 RCPs per cell (IQR, 141-295; n = 199) (****p < 0.0001, Mann-Whitney U test). In contrast, total *ACTB* RNA showed no significant difference between the two groups, with a median of 259 RCPs per cell (IQR, 170-340; n = 235) in treated cells and 261 RCPs per cell (IQR, 161-346; n = 193) in untreated cells (ns, p ≥ 0.05, Mann-Whitney U test). These results demonstrate that Ribo-ISS successfully reports on ribosome-mRNA binding dynamics without confounding transcriptional signals.

### Single-Cell Translatome Profiling of Mouse Brain by Ribo-ISS

We next applied Ribo-ISS to mouse brain tissue for targeted spatial profiling of 211 ribosome-associated transcripts. Two adjacent brain sections were analyzed in parallel to generate translatome and transcriptome datasets, respectively. Unsupervised clustering and cell-type annotation identified nine major cell populations in both datasets (Figure 3a,b), including excitatory neurons (EXC), inhibitory neurons (INH), di- and mesencephalon neurons (DE/MEN), astrocytes (AC), oligodendrocytes (OLG), microglia (MLG), oligodendrocyte precursor cells (OPC), vascular cells (VAS), and choroid epithelial/ependymal cells (CHOR/EPEN).

**Figure 3.**
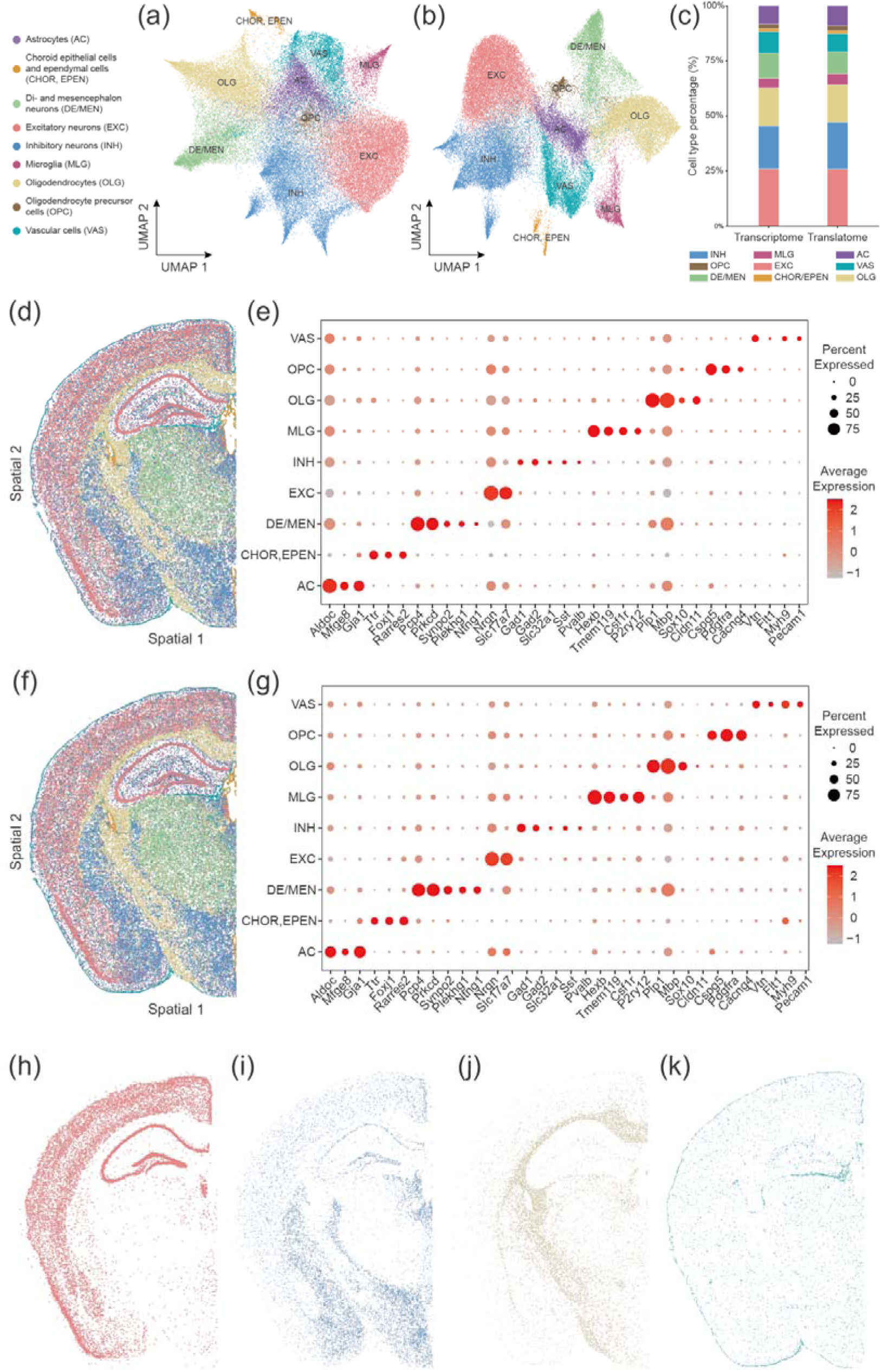
Application of Ribo-ISS to mouse brain cell typing. (a and b) UMAP visualizations of major cell type identified by unsupervised clustering of translatomic (a) and transcriptomic (b) datasets. (c) Stacked bar plot comparing the proportions of major cell types between translatomic and transcriptomic datasets. (d and f) Spatial mapping of major cell types in the translatomic (d) and transcriptomic (f) datasets. (e and g) Dot plots showing the expression of cell-type-specific marker genes in the translatomic (e) and transcriptomic (g) datasets. (h-k) Spatial distributions of representative cell types in the translatomic dataset, including EXC (h), INH (i), OLG (j), VAS (k).

The translatome and transcriptome datasets showed highly consistent cellular compositions and cell-type annotations, indicating that ribosome-associated transcript profiles retain sufficient molecular information for accurate cell identity assignment (Figure 3c). In addition, strong concordance was observed between the two datasets at both gene expression and spatial organization levels (Figure 3d-g). Major brain cell types displayed expected anatomical distributions in the Ribo-ISS translatome map, with excitatory neurons enriched in cortical layers and hippocampal regions (Figure 3h), inhibitory neurons distributed across deep cortical layers and limbic structures (Figure 3i), oligodendrocytes concentrated in white matter tracts and deep cortical regions (Figure 3j), and vascular cells forming continuous vascular networks consistent with the organization of the blood-brain barrier (Figure 3k). Spatial maps of additional cell populations are shown in Figure S2.

Together, these results demonstrate that Ribo-ISS enables single-cell spatial translatome profiling in complex tissues while preserving accurate cell-type identification and anatomical organization. By directly measuring ribosome-associated transcripts, Ribo-ISS provides an additional layer of information beyond conventional transcriptome profiling, enabling investigation of cell-state heterogeneity and spatial regulation of translational programs within intact tissues.

### In situ translatome profiling reveals cell-type-specific translational responses to sleep deprivation

Following the systematic validation of Ribo-ISS in mouse brain tissue, we applied this technology to investigate spatially resolved translational responses in a mouse model of sleep deprivation. Mice were subjected to continuous 48-hour sleep deprivation, while control animals were maintained under identical housing conditions with normal sleep-wake cycles. Brain tissues were collected, and adjacent sections were analyzed in parallel using Ribo-ISS and transcriptome profiling to simultaneously capture ribosome-associated and total RNA expression patterns. Ribo-ISS showed high reproducibility between replicates (Pearson *r* = 0.995; Figure S3), validating its reliability for subsequent spatial analysis. A panel of 193 quality-filtered genes was targeted for spatial profiling. Single-cell datasets from control and sleep-deprived brains were integrated, followed by unsupervised clustering and cell-type annotation. Consistent with previous analyses, both translatome and transcriptome datasets resolved nine major brain cell populations (Figure 4a,b).

**Figure 4.**
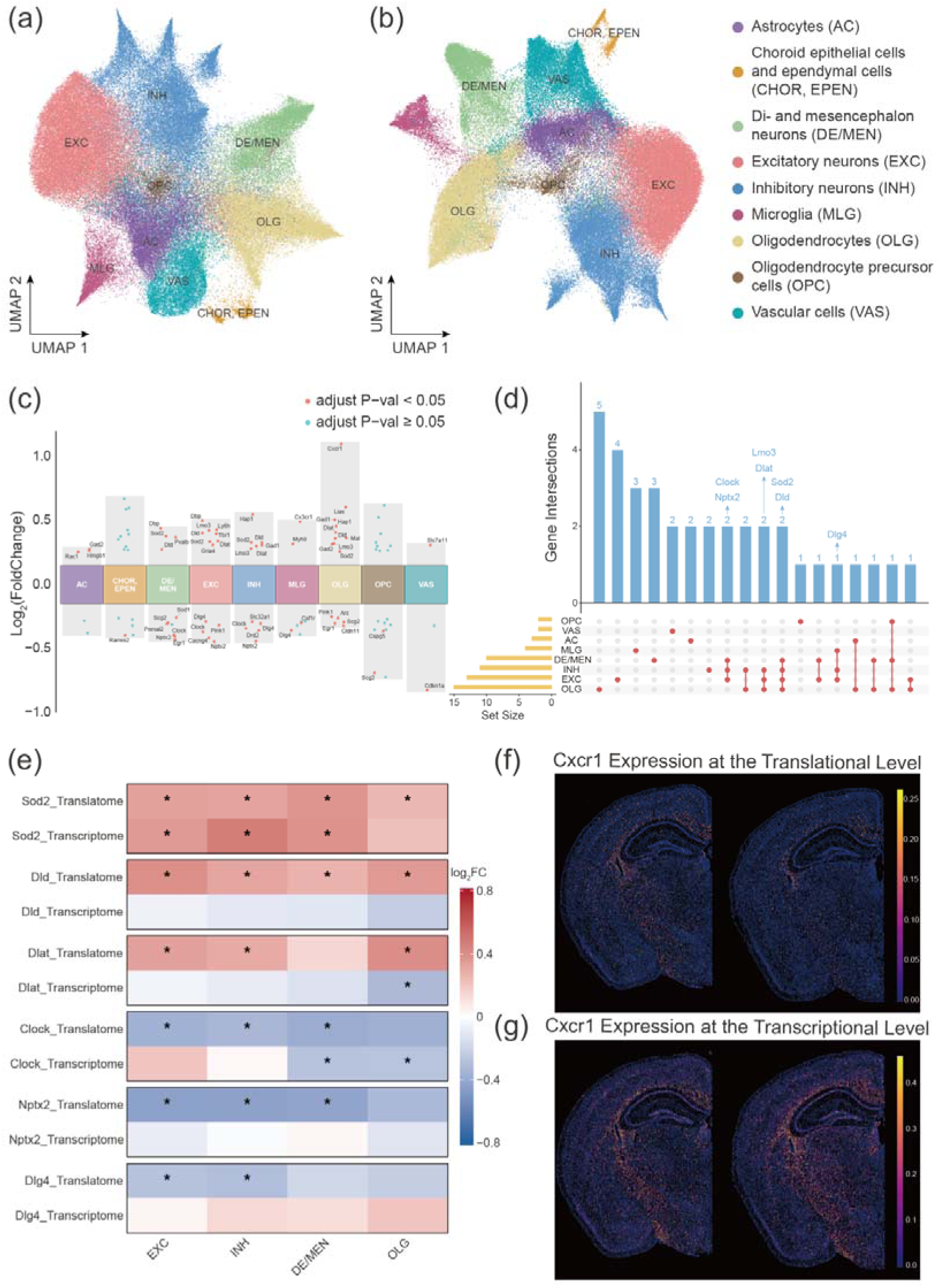
Differential profiling of sleep deprivation by Ribo-ISS. (a and b) UMAP visualization of major cell types after integrating sleep deprivation and control samples via unsupervised clustering using translatomic (a) and transcriptomic (b) data. (c) Grouped volcano plots showing cell-type-specific DEGs identified in the translatome between sleep deprivation and control groups across major cell types. (d) UpSet plot depicting intersections of DEG sets across the major cell types defined in (c). (e) Heatmap comparing log_2_ fold changes of shared DEGs from (d) in neurons and OLG between translatomic and transcriptomic profiles. * indicates statistically significant differences. (f and g) Spatial distribution of *Cxcr1* expression in the tranlatmoe (f) and transcriptome (g), with sleep deprivation (left) and control (right) groups.

We next identified differentially expressed genes (DEGs) across individual cell types in both datasets (Figure 4c, Figure S4-S6) and focused on genes specifically altered at the translatome level. Notably, *Sod2* was the only DEG exhibiting concordant upregulation at both RNA and ribosome-associated levels across four cell types (Figure 4d,e), suggesting a conserved antioxidant response induced by sleep deprivation. In contrast, several genes displayed prominent changes at the ribosome-associated level without corresponding transcriptional alterations, highlighting the contribution of post-transcriptional regulation to cellular adaptation.

For example, mitochondrial metabolic enzymes *Dld* and *Dlat* showed increased ribosome-associated signals without significant changes in transcript abundance in major neuronal populations and oligodendrocytes (Figure 4e). *Dld* exhibited the strongest increase in excitatory neurons (32.2%), whereas *Dlat* showed the largest elevation in oligodendrocytes (32.7%). In addition, *Lias* displayed a 52.2% increase in oligodendrocytes, further suggesting enhanced mitochondrial metabolic activity at the translational level in response to sleep deprivation-associated stress (Figure 4c). Conversely, neuronal genes associated with circadian regulation and synaptic function, including *Clock*, *Nptx2*, and *Dlg4*, showed reduced ribosome-associated signals without significant transcriptional changes (Figure 4e). *Clock* displayed the largest decrease in DE/MEN neurons (24% reduction), *Nptx2* in excitatory neurons (26.6% reduction), and *Dlg4* in inhibitory neurons (19% reduction), indicating cell-type-specific translational suppression of neuronal functional programs.

Among all cell populations, oligodendrocytes exhibited the greatest number of differentially regulated genes, and shared differential genes were predominantly observed between neuronal and oligodendrocyte populations, suggesting coordinated neuron-glia responses during sleep deprivation (Figure 4c). Notably, *Cxcr1* showed significant elevation at the translatome level in oligodendrocytes (log_2_FC = 1.10, adjusted p-value = 6.49 × 10^-80^), whereas no corresponding change was detected at the transcriptome level (log_2_FC = −0.18, adjusted p-value = 9.55 × 10^-51^, Figure 4f-g). This finding highlights the ability of Ribo-ISS to identify translationally regulated genes that are not apparent from conventional transcriptomic measurements and suggests a potential role for translational control in shaping glial responses under sleep deprivation-associated stress.

### Functional Consequences of Translatome Changes in Sleep-Deprived Mouse Brain

To investigate the functional implications of sleep deprivation-induced translatome remodeling, we performed gene set variation analysis (GSVA) on matched transcriptomic and translatomic datasets. By comparing pathway activity changes between transcriptional and translational layers, we identified three major regulatory patterns (Figure 5a-d). The first pattern was characterized by translation-prioritized activation, in which unchanged or reduced transcriptional levels were accompanied by increased ribosome-associated signals. This mode was predominantly observed in pathways related to energy metabolism and innate immune responses. For example, cytosolic DNA-sensing, Toll-like receptor, and chemokine signaling pathways showed preferential activation at the translational level. Consistently, enhanced ribosome association of TCA cycle-related genes was accompanied by increased translational signals of mitochondrial metabolic enzymes, including *Dld*, *Dlat*, and *Lias*, suggesting that cells rapidly adjust metabolic programs through translational regulation during sleep deprivation.

**Figure 5.**
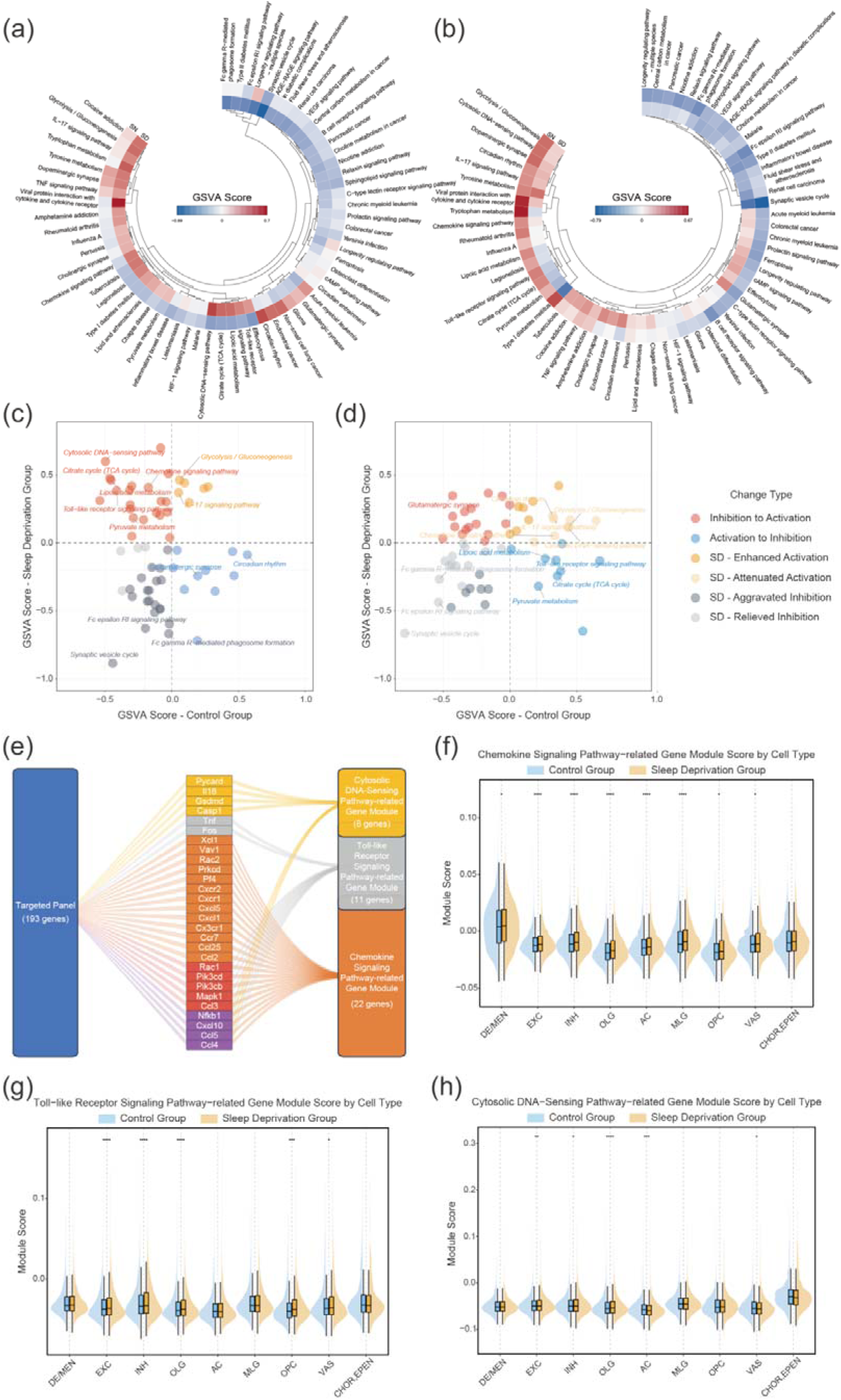
Translationally regulated activity states during sleep deprivation identified by Ribo-ISS. (a and b) Heatmaps of GSVA scores for pathway activities in the translatome (a) and transcriptome (b). (c and d) Scatter plots of GSVA scores for representative pathways in the translatome (c) and transcriptome (d). The X- and Y-axes represent GSVA scores in the control and sleep deprivation groups, respectively. Deviation frome the identity line (y=x) indicates the direction and extent of pathway activity dysregulation following sleep deprivation. (e) Gene composition of the innate immune pathway-related modules used for AddModuleScore analysis. (f-h) Module activity scores for innate immune pathways across cell types, including the chemokine signaling pathway (f), the toll-like receptor signaling pathway (g), and the cytosolic DNA-sensing pathway (h), calculated using AddModuleScore. Statistical significance was determined by the Wilcoxon rank-sum test, *p < 0.05, **p < 0.01, ***p < 0.001, ****p < 0.0001.

The second pattern represented translation-prioritized suppression, characterized by upregulated transcriptional levels but actively suppressed translation. This pattern was observed in pathways associated with synaptic function and adaptive immune regulation. For example, glutamatergic synapse and synaptic vesicle cycle pathways exhibited reduced translational activity despite upregulated transcriptional GSVA scores, indicating selective translational suppression of neuronal functional programs under sleep deprivation. The third pattern involved coordinated transcriptional and translational regulation, exemplified by the circadian rhythm pathway, which showed concordant downregulation at both levels. Together, these three regulatory modes demonstrate that sleep deprivation induces complex translatome remodeling, involving selective translational activation, repression, and transcription-translation coordination across distinct biological processes.

To further characterize cell type-specific translational regulation of stress-responsive pathways, we examined three representative innate immune-related pathways, including cytosolic DNA sensing, Toll-like receptor signaling, and chemokine signaling pathways. Gene sets corresponding to these pathways were extracted from the targeted panel (Figure 5e) and scored at single-cell resolution using AddModuleScore. The resulting translational pathway maps revealed substantial cell-type-specific heterogeneity in response to sleep deprivation. Chemokine signaling-related genes exhibited increased ribosome-associated activity across multiple cell populations, including excitatory neurons, inhibitory neurons, oligodendrocytes, microglia, and astrocytes (Figure 5f), indicating broad engagement of neuroimmune-related translational programs beyond canonical immune cell populations. Toll-like receptor signaling showed preferential translational activation in neuronal and oligodendrocyte populations, whereas relatively limited changes were observed in microglia and astrocytes (Figure 5g), suggesting distinct cellular responses to sleep deprivation-associated stress. In contrast, cytosolic DNA-sensing pathways displayed divergent regulation among glial populations, with increased translational activity in astrocytes but reduced activity in oligodendrocytes (Figure 5h). These distinct patterns highlight cell-type-specific strategies of translational regulation and demonstrate the ability of Ribo-ISS to resolve spatially heterogeneous translatome remodeling within complex tissues.

## Conclusion

To address the need for spatially resolved translatome profiling at single-cell resolution without genetic manipulation, we developed Ribosome-tethered In Situ Sequencing (Ribo-ISS), a high-throughput in situ sequencing technology for mapping ribosome-associated mRNAs within intact tissues. Ribo-ISS employs a ribosome-dependent tri-probe architecture, in which the connect probe simultaneously recognizes endogenous 18S rRNA and mRNA-associated split probes, enabling selective detection of ribosome-associated transcripts. Through systematic validation, we demonstrated that Ribo-ISS achieves high specificity by integrating ribosome-dependent molecular anchoring with precise target recognition. This streamlined probe design provides an alternative strategy for spatial translatome profiling with enhanced accessibility and scalability.

Application of Ribo-ISS in mouse brain tissue demonstrated its ability to generate single-cell spatial translatomic maps while preserving tissue architecture. Translatome profiles successfully recapitulated major brain cell populations and their anatomical organization, showing strong concordance with transcriptomic measurements. In a sleep deprivation model, simultaneous profiling of transcriptional and translational layers revealed extensive transcription-translation uncoupling and uncovered distinct modes of translational regulation, including translation-prioritized activation, translation-prioritized suppression, and coordinated transcriptional-translational regulation. These findings highlight the importance of translational control in shaping cell-type-specific responses to physiological stress and demonstrate the unique information provided by spatial translatome profiling beyond conventional transcriptomic analysis.

Compared with existing approaches, Ribo-ISS provides several advantages. First, it enables spatial translatome analysis without requiring transgenic expression or exogenous ribosome labeling, allowing direct investigation of intact tissue architecture. Second, by integrating molecular detection with spatial coordinates, Ribo-ISS preserves cellular identity and microenvironmental context, enabling cell-type-resolved analysis of translational regulation. Third, its modular probe architecture is compatible with highly multiplexed in situ sequencing strategies, providing a flexible framework for future expansion.

Several limitations remain to be addressed. Similar to other targeted spatial profiling technologies, the current implementation of Ribo-ISS is constrained by the size of the predefined gene panel, limiting comprehensive characterization of genome-wide translational programs. However, the underlying probe chemistry and sequencing-by-ligation framework are inherently compatible with increased molecular coverage through expanded probe libraries and improved encoding strategies. Future integration with advanced cyclic sequencing chemistry, high-density probe synthesis, and enhanced imaging platforms may enable progression toward spatially resolved whole-translatome profiling.

In addition, Ribo-ISS currently reports ribosome-associated transcript localization rather than directly measuring ribosome occupancy, translation kinetics, or nascent peptide production. Combining Ribo-ISS with complementary approaches, including spatial proteomics, spatial metabolomics, and other multimodal imaging technologies, may provide a more complete view of gene regulation from transcript availability to translational output and protein production. Furthermore, systematic validation across diverse physiological and pathological models, together with temporal profiling of dynamic translational responses, will be essential for uncovering the broader biological principles governed by spatial translation regulation.

In summary, Ribo-ISS establishes a spatial translatomics framework that integrates high-throughput molecular detection, single-cell resolution, and in situ spatial analysis. By enabling the construction of spatial translatome maps within intact tissues, Ribo-ISS provides a new dimension for investigating translational regulation and offers a versatile platform for exploring complex tissue functions and disease mechanisms through the lens of translational control.

## Experimental Section

### Cell culture and sample preparation

H1299 (Cell Bank of the Chinese Academy of Sciences, Shanghai Branch) and MCF-7 (ATCC) cells were maintained in a humidified incubator at 37 with 5% CO2. H1299 cells were cultured in RPMI-1640 medium (Shanghai Basalmedia Technologies, L210KJ), whereas MCF-7 cells were cultured in high-glucose DMEM (Shanghai Basalmedia Technologies, L110KJ). Both media were supplemented with 10% fetal bovine serum (FBS, Gibco, 10099-141C). Upon reaching sufficient confluence, cells were detached using 0.25% trypsin-EDTA (Shanghai Basalmedia Technologies, S330JV) and seeded onto adhesive microscope slides (Citotest, 188105) preplaced in EasYDish (Nunc, Thermo Scientific, 150468) containing complete growth medium. The cells were allowed to attach and grow under identical conditions until they reached approximately 60% confluence. For a subset of H1299 cells, the culture medium was replaced with fresh complete growth medium containing 200 μg/mL puromycin (Sangon Biotech, B607054), and the cells were incubated for 1 h at 37 with 5% CO2. The culture medium was then removed, and the slides were washed twice with diethyl pyrocarbonate (DEPC, Sigma-Aldrich, D5758)-treated 1× phosphate-bufferd saline (DEPC-PBS) for 3 min each. Fixation was performed by incubating the slides in 4% paraformaldehyde (PFA, Sigma-Aldrich, 16005) in 1× DEPC-PBS for 30 min at room temperature. After two additional washes with 1× DEPC-PBS for 3 min each, the slides were dehydrated through a graded ethanol series of 70%, 85%, and absolute ethanol for 5 min per step. Following air drying at room temperature, the slides were stored at −80 until further use.

### Animals, sleep deprivation, and tissue processing

In this study, 8-week-old male C57BL/6 mice (purchased from Jiangsu GemPharmatech Co., Ltd, Nanjing, China) were used. The source of the experimental mice and all animal procedures were approved by the Laboratory Animal Management Ethics Committee of the Medical College of Huaqiao University (Approval No. A2024077). The mice were housed in a specific pathogen-free animal facility with free access to food and water under a 12 h light/dark cycle. They were randomly divided into the normal control group and the sleep deprivation group. Mice in the sleep deprivation group were placed in individual cages attached to a sleep deprivation apparatus and subjected to continuous 48 h sleep deprivation. Mice in the control group were housed under standard conditions with an undisturbed sleep-week cycle and received no intervention. After 48 h of sleep deprivation, mice from both groups were subjected to sample collection. The mice were anesthetized, perfused with saline to clear the blood, and then euthanized. The whole brain was rapidly dissected out. The brain samples were then embedded in optimal cutting temperature compound (OCT, SAKURA, 4583). The frozen OCT blocks were trimmed and sectioned at a thickness of 10 μm onto adhesive glass slides using a Leica cryostat. The slides were stored at −80 until further use.

### Sample pretreatment

For cell samples on slides, the samples were treated with 1× DEPC-PBS for 5 min and subsequently permeabilized with ice-cold methanol at −80 for 1 h. For fresh frozen mouse brain tissue sections, slides were equilibrated at room temperature for 10 min after removal from −80. The sections were then fixed with 4% PFA in PBS at room temperature for 10 min, rinsed twice in 1× DEPC-PBS for 2 min each time, and permeabilized with ice-cold methanol at −80 for 1 h. Prior to subsequent steps, ImmEdge Hydrophobic Barrier Pen (Vector Laboratories, H-4000) was applied to circumscribe the sample area to retain the reaction solution on the cell and tissue.

### Ribo-ISS probe hybridization and enzymatic circularization

Prior to hybridization, samples were washed three times with 0.1% (v/v) Tween-20 (Sigma-Aldrich, P9416) in 1× DEPC-PBS (DEPC-PBST). Next, 0.1 μM connect probes and 0.04 μM split probes in a hybridization buffer containing 10% formamide (Sigma-Aldrich, 47671), 4× SSC (Sigma-Aldrich, S6639), 5% PEG-4000 was added to the samples, followed by overnight incubation at 37. After two washes with DEPC-PBST for 2 min each, the circularization reaction mixture containing 0.5 U/μL SplintR Ligase (NEB, M0375L), 1× SplintR Ligase reaction buffer, 25% (v/v) glycerol (Sigma-Aldrich, G9012), 2 mg/mL BSA (Sangon Biotech, B600036), and 5% PEG-4000 was applied and incubated at 37 for 30 min. The samples were subsequently rinsed twice with DEPC-PBST for 2 min each.

### IISS probe hybridization and enzymatic circularization

Samples were rinsed three times in DEPC-PBST before hybridization. Next, 0.04 μM split probes in a hybridization buffer containing 10% formamide, 4× SSC, 5% PEG-4000 was added to the samples, followed by incubation at 37 for 2 h. After two washes with DEPC-PBST for 2 min each, the circularization reaction mixture containing 0.5 U/μL SplintR Ligase, 1× SplintR Ligase reaction buffer, 25% (v/v) glycerol, 2 mg/mL BSA, 5% PEG-4000, and 0.5 μM splint oligonucleotide was applied and incubated at 37 for 30 min. Finally, the samples were washed twice with DEPC-PBST for 2 min per wash.

### Rolling circle amplification and in situ sequencing of RCPs

An RCA reaction mix consisted of 1 U/μL Phi29 DNA Polymerase (Thermo Scientific, EP0094), 1× Phi29 DNA Polymerase reaction buffer, 5% (v/v) glycerol, 2 mg/mL BSA, 1 mM dNTPs (Thermo Scientific, R0182) and 5% PEG-4000. The incubation was carried out at 37 overnight, followed by two washes for 2 min each with DEPC-PBST.

For single-target detection in cell samples, a mixture containing 0.1 μM fluorescently labeled detection probes, 2× SSC, 20% formamide, and 20 μg/mL Hoechst 33342 was applied to the samples and incubated at 30 for 30 min. The samples were then washed twice with DEPC-PBST for 2 min per was, air-dried, and mouted in Slowfade Gold Antifade Mountant medium (Thermo Scientific, S36936).

For high-throughput detection in tissue samples, an improved in situ sequencing approach was employed. Prior to the first sequencing cycle, the samples were treated twice 50% formamide at room temperature for 10 min each, followed by three washes with DEPC-PBST. A sequencing reaction mixture containing 0.1 U/μL T4 DNA ligase (Thermo Scientific, EL0012), 1× T4 DNA ligase reaction buffer, 1 mM ATP (Thermo Scientific, R0441), 2 mg/mL BSA, 20 μg/mL Hoechst 33342, 0.2 μM anchor primer, and 0.2 μM fluorescently labeled sequencing interrogation probes was applied and incubated at 30 for 45 min. The samples were then washed twice with DEPC-PBST for 2 min per wash, air-dried, and mouted in Slowfade Gold Antifade Mountant medium.

Images were acquired using a Leica DM6B fluorescence microscope equipped with DFC9000GT camera and a 20× objective.

To stripping off ligated sequencing probes and prepare the samples for the next sequencing cycle, the samples were treated three times with stripping buffer containing 50% formamide, 0.05× SSC, 0.5% SDS and 0.5 mM EDTA at 37 for 10 min each, followed by three washes with DEPC-PBST. The sequencing reaction, imaging, and stripping procedures were repeated for subsequent sequencing cycle.

### Transcript decoding and generation of single-cell spatial expression matrix

Image processing and decoding were performed as follows. To achieve precise spatial registration across imaging rounds, feature-point matching was performed on nuclear-stained images from each round as the reference channel. Inter-round transformation matrices were computed and applied to correct the remaining fluorescent channels, enabling accurate multi-round image alignment. Background signals were subsequently suppressed using top-hat filtering. RCPs were detected as fluorescent spots in each base channel and mapped onto a unified reference layer. Base scores were calculated based on the relative grayscale contrast between the spot center its surrounding neighborhood; the channel with the highest score was assigned as the base call at each locus, and the corresponding sequencing quality score was derived accordingly. Fluorescent signals across multiple rounds were spatially matched and aligned within a common coordinated framework. The base calls from individual rounds were concatenated to generate a barcode for each signal spot, which was then compared against the designed gene barcode library to decode gene identity. Quality scores from the four base-calling rounds were aggregated, and the total quality score was computed for each transcript. Transcripts ranking in the bottom 5% by total quality score were filtered out to retain high-confidence molecules for subsequent spatial expression analysis. To construct the single-cell expression matrix, nuclei were first segmented from the grayscale nuclear-stained images of the reference round using Cellpose3. To ensure that transcripts localized in the cytoplasm were correctly assigned to their corresponding cells, the nuclear masks were expended by 50 pixels. Finally, each transcripts was assigned to its nearest cell based on the expanded segmentation.

### Single-cell data processing and downstream analysis

The original gene panel comprised 211 genes. Gene-level filtering was performed using the negative control probe dapB as a reference threshold. For each gene, total expression across all cells and mean expression in positive cells were calculated, where positive cells were defined as those with detectable expression greater than zero. Genes satisfying both exclusion criteria, specifically those with total expression and mean positive-cell expression each lower than the corresponding dapB values, were classified as low-abundance noise targets and removed. This filter was independently applied to the transcriptomic and translatomic datasets. The union of excluded genes from both omics layers formed a unified exclusion list, ensuring identical gene content between the two panels. Cell-level quality control was applied to exclude cells with fewer than five detected transcripts, retaining high-quality single-cell profiles.

The filtered expression matrix was analyzed using the standard Seurat (v5.4.0) workflow in R (v4.5.3). Expression data were normalized by LogNormalize and scaled with ScaleData for downstream dimensionality reduction. Given the limited size of this targeted panel, all genes were retained as variable features for principal component analysis without additional feature selection to avoid information loss. Unsupervised clustering was performed using Louvain algorithm based on the top 16 principal components, and the results were visualized in two dimensions via UMAP. Cluster identities were manually refined by merging closely related subpopulations. Cell type labels were assigned through manual annotation based on cluster-specific marker expression and established reference signatures. Spatial coordinates derived from cell centroid files were matched to the expression matrix by unique cell identifiers and integrated into the Seurat metadata to enable spatially resolved visualization.

Differential expression analysis was performed using the Seurat package. For each cell type, differentially expressed genes (DEGs) between the sleep deprivation group and control group were identified using the FindMarkers function with the MAST (Model-based Analysis of Single-cell Transcriptomics) test. Only genes detected in at least 10% of cells in either group and with a minimum log2 fold change of 0.25 were considered. P-values were adjusted for multiple testing using the Bonferroni correction, and genes with adjusted P-value < 0.05 were defined as significant DEGs.

Pathway activity was assessed using the GSVA (v2.2.0) package. KEGG pathway gene sets for mouse (Mus musculus, mmu) were retrieved via the KEGGREST package. Gens in the expression matrix were mapped to KEGG gene sets, and only pathways with ≥ 2 matched genes were retained for GSVA analysis. Pseudo-bulk expression matrices were constructed using the AverageExpression function in Seurat, and enrichment scores for each pathway in the sleep deprivation group and control group were estimated based on the GSVA algorithm. To identify well-represented pathways in the dataset, overlap statistics between pathways and genes were integrated with GSVA scores. Specifically, panel-targeted genes in this study were matched against each pathway gene set. The top 70 pathways were selected by gene match percentage, and pathways with fewer than 5 matched genes were further excluded. Ultimately, 61 representative pathways were obtained for subsequent visualization and data analysis.

The AddModuleScore function in the Seurat package was used to compute gene module scores for individual cells. By integrating cell type annotations and sample group information, the activity of representative pathways across different cell types was evaluated. Statistical significance between groups was assessed using the Wilcoxon rank-sum test, with significance levels denoted as follows: ns, not significant; *, significant (p < 0.05); **, very significant (p < 0.01); ***, highly significant (p < 0.001); ****, extremely significant (p < 0.0001).

## Acknowledgements

This work was supported by the Natural Science Foundation of Fujian Province (2026J002030), the Fundamental Research Funds for the Central Universities of Huaqiao University (ZQN-1123), and Huaqiao University Research Startup Funds (23BS115, 24BS139). The authors would like to thank Dr. Taobo Hu and Dr. Mengping Long for valuable discussions on data analysis.

## Data Availability Statement

The data that support the findings of this study are available from the corresponding author upon reasonable request.

