## Supplementary Material for "Ribosome-tethered *in situ* sequencing for single-cell translatome analysis"

### Supplementary Figures

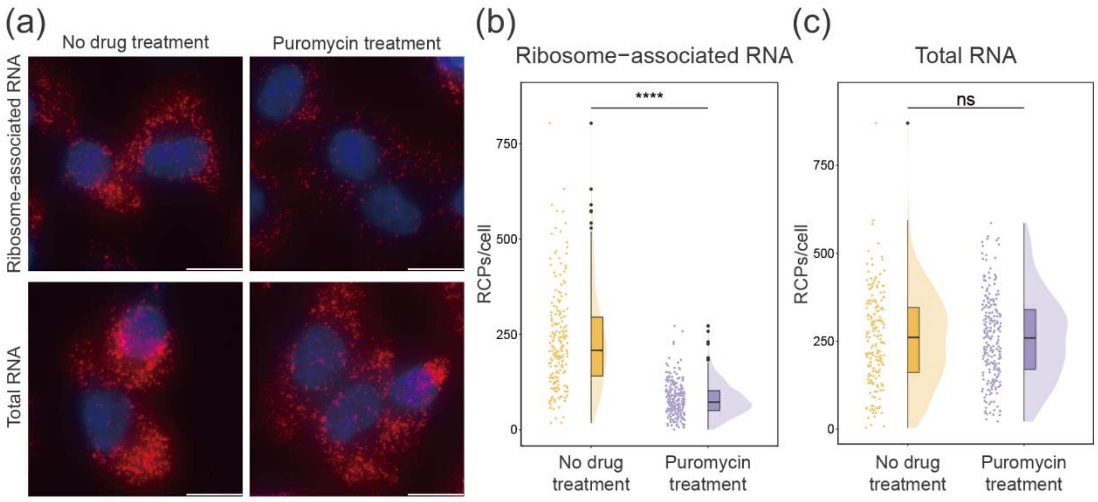

**Figure S1. Puromycin-induced changes in ACTB expression in H1299 cells.** (a) Representative fluorescence images of ribosome-associated ACTB RNA (top) and total ACTB RNA (bottom) without or with puromycin treatment. Scale bar, 20 μm. Quantification of ribosome-associated RNA (b) and total RNA (c) from the experiment shown in (a). Statistical significance was determined by Mann-Whitney U test, ****$p < 0.0001$; ns, $p \geq 0.05$ (not significant).

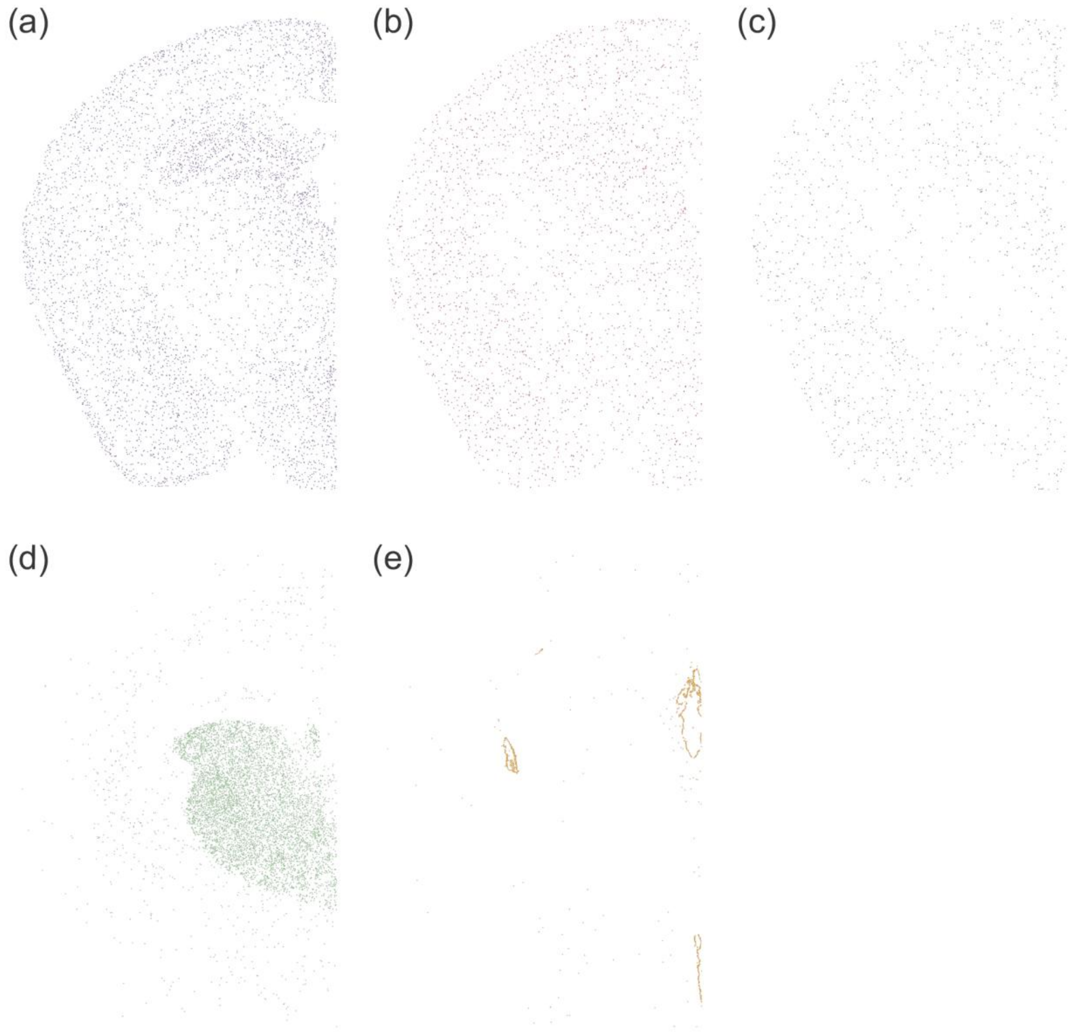

**Figure S2. Spatial mapping of major cell types at single-cell resolution based on Ribo-ISS.** (a) AC. (b) MLG. (c) OPC. (d) DE/MEN. (e) CHOR, EPEN.

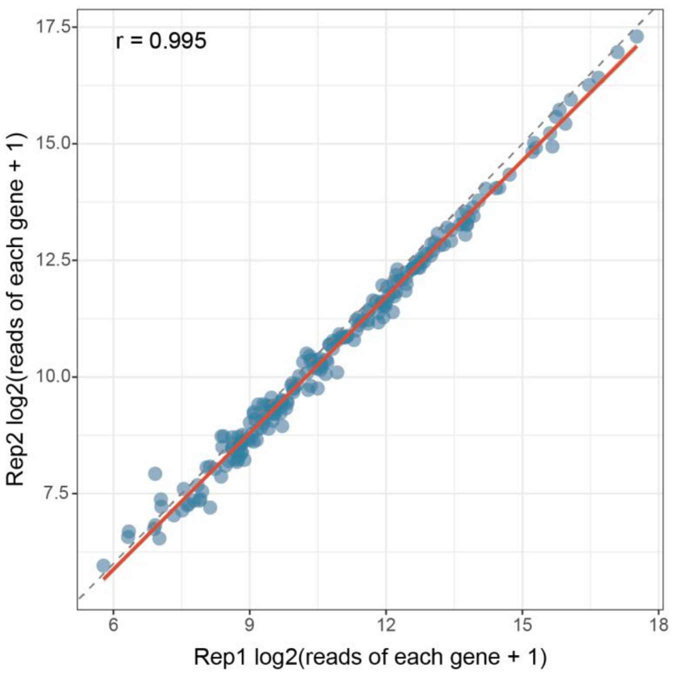

**Figure S3. Correlation of gene read counts for 211 genes between two replicate samples**.

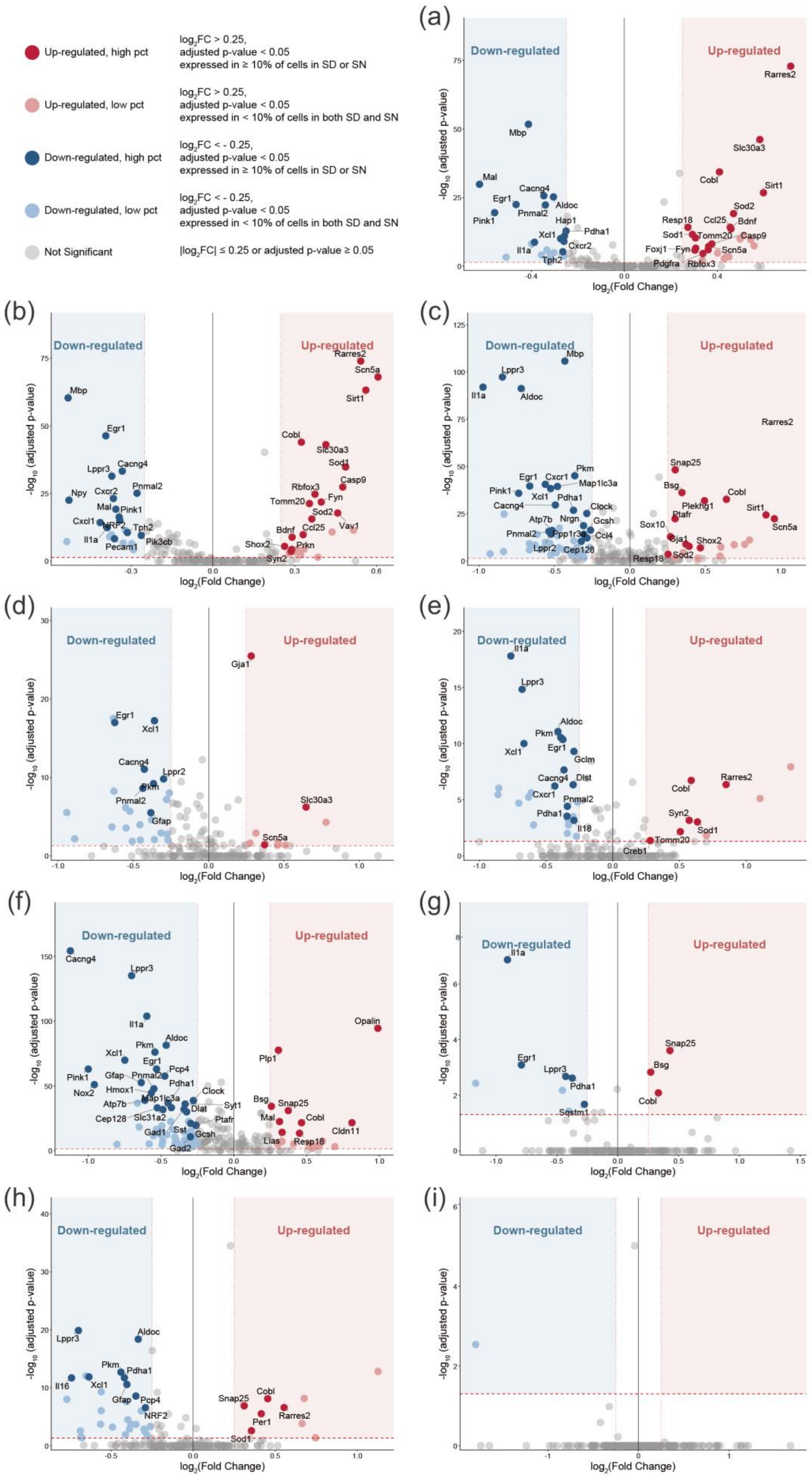

**Figure S4. Volcano plots of cell type-specific differentially expressed genes (DEGs) between sleep deprivation and control groups, identified by transcriptome analysis.** (a) INH. (b) EXC. (c) DE/MEN. (d) AC. (e) MLG.(f) OLG. (g) OPC. (h) VAS. (i) CHOR, EPEN.

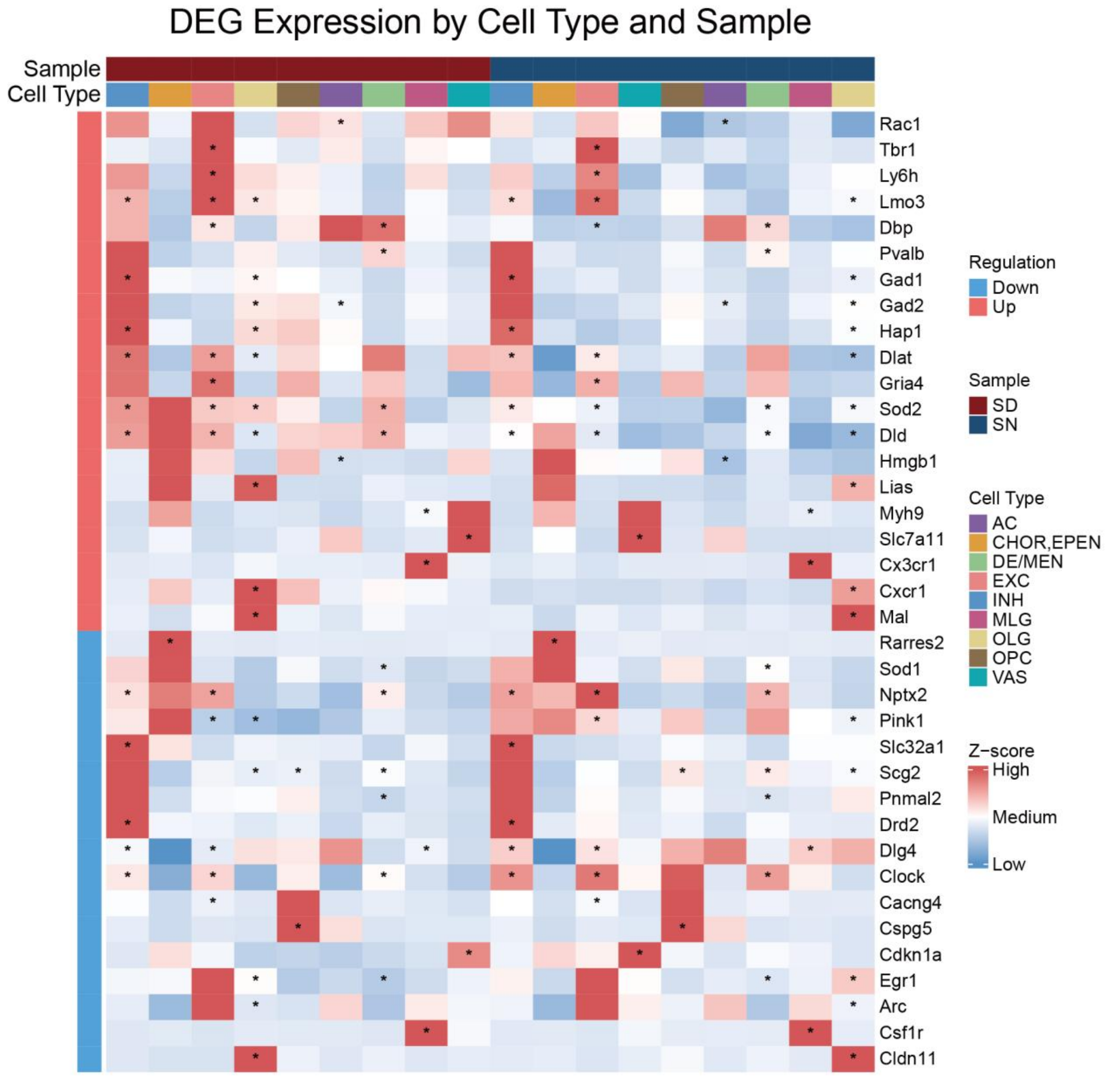

**Figure S5. Heatmap of significant DEGs identified from translatome data across celltype.** Each row represents a gene; columns are grouped by cell type and condition, sleep deprivation group (SD) versus sleep-normal control group (SN). ＊ indicates significant DEGs for that column-specific cell type.

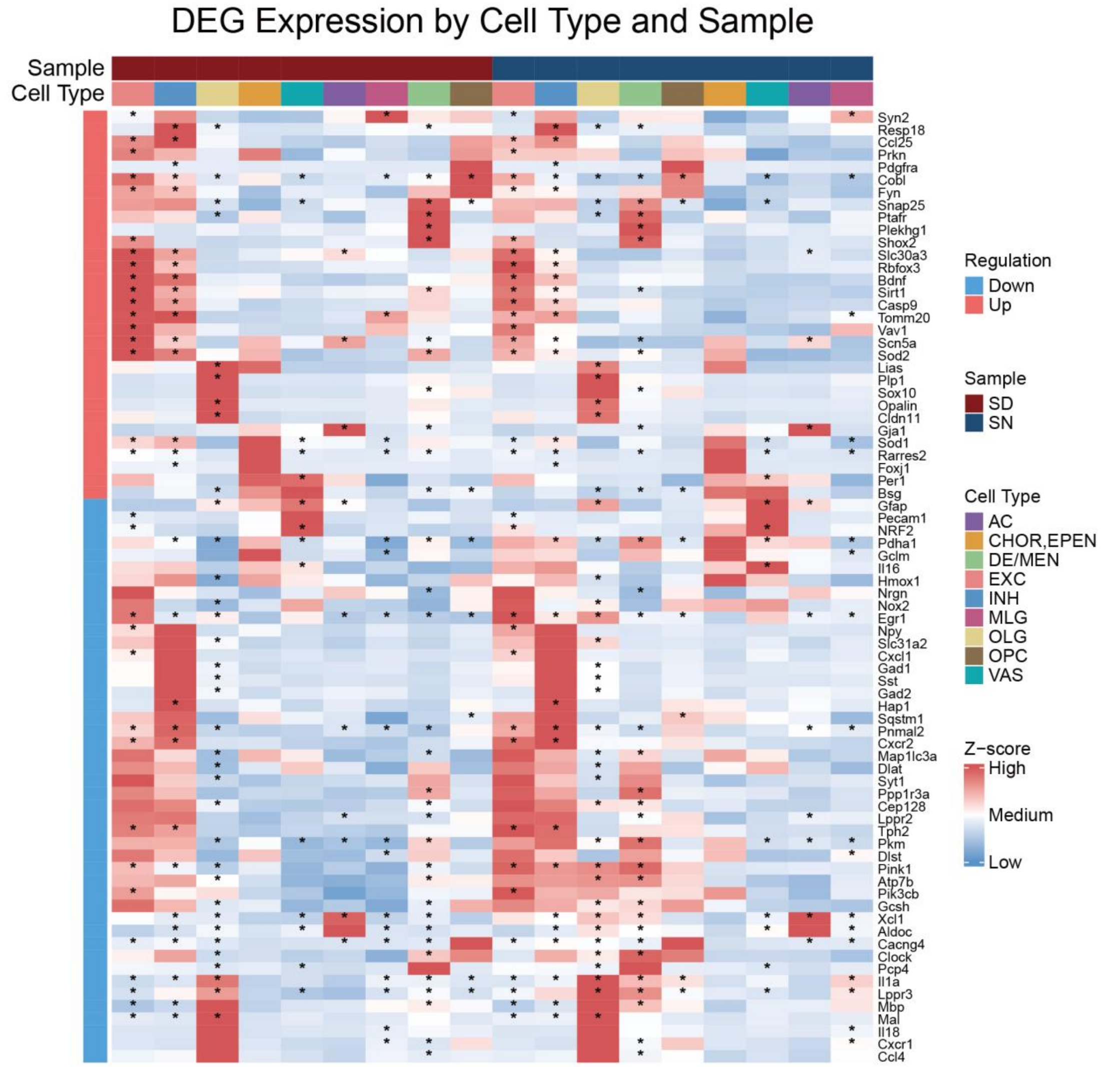

**Figure S6. Heatmap of significant DEGs identified from transcriptome data across celltype.** Each row represents a gene; columns are grouped by cell type and condition, sleep deprivation group (SD) versus sleep-normal control group (SN). ＊ indicates significant DEGs for that column-specific cell type.

### Supplementary Tables

**Table S1.** Oligonucleotide sequences used in Ribo-ISS. Split probes need to be either modified a phosphate group at the 5' end during oligonucleotide synthesis or subjected to phosphorylation using T4 polynucleotide kinase after synthesis, before being used.

| Oligonucleotide name | Sequence (5'-3') |
|---|---|
| ACTB-1-SPL | /5Phos/AGGCTCTGTGCTCGCGAAAAAACTCATCAGTCAGGTCATACACTA |
| ACTB-1-SPR | /5Phos/AAGATAAACATCGTAGACTACTGAACTCACGGATCGGCAAAGGCG |
| ACTB-2-SPL | /5Phos/TGGTGCCTGGGGCGCCAAAAAACTCATCAGTCAGGTCATACACTA |
| ACTB-2-SPR | /5Phos/AAGATAAACATCGTAGACTACTGAACTCAGCCCACCATCACGCCC |
| ACTB-3-SPL | /5Phos/TGGATAGCAACGTACAAAAAAACTCATCAGTCAGGTCATACACTA |
| ACTB-3-SPR | /5Phos/AAGATAAACATCGTAGACTACTGAACTCACAGGGATAGCACAGCC |
| ACTB-4-SPL | /5Phos/AGGATCTTCATGAGGTAAAAAACTCATCAGTCAGGTCATACACTA |
| ACTB-4-SPR | /5Phos/AAGATAAACATCGTAGACTACTGAACTCAGTAGCCGCGCTCGGTG |
| ACTB-5-SPL | /5Phos/TTTGCGGATGTCCACGAAAAAACTCATCAGTCAGGTCATACACTA |
| ACTB-5-SPR | /5Phos/AAGATAAACATCGTAGACTACTGAACTCATGTTGGCGTACAGGTC |
| ACTB-6-SPL | /5Phos/TGCGCAAGTTAGGTTTAAAAAACTCATCAGTCAGGTCATACACTA |
| ACTB-6-SPR | /5Phos/AAGATAAACATCGTAGACTACTGAACTCAATCTCATCTTGTTTTC |
| MALAT1-1-SPL | /5Phos/CTGAACCGGAGCAGGAAAAAAACTCACTGATCAGGTCATACACTA |
| MALAT1-1-SPR | /5Phos/AAGATAAACATCGTAGACTATCAGACTCAATGAGCTTCAGACCTT |
| MALAT1-2-SPL | /5Phos/TCATGCCCACAAGGATAAAAAACTCACTGATCAGGTCATACACTA |
| MALAT1-2-SPR | /5Phos/AAGATAAACATCGTAGACTATCAGACTCATGAAACCGATTATGGA |
| MALAT1-3-SPL | /5Phos/ACTGCCAACTAATTGCAAAAAACTCACTGATCAGGTCATACACTA |
| MALAT1-3-SPR | /5Phos/AAGATAAACATCGTAGACTATCAGACTCACCAACCGTAACAGGCC |
| MALAT1-4-SPL | /5Phos/TTCCGCCGCCTTTGTGAAAAAACTCACTGATCAGGTC |

| | |
|---|---|
| | ATACACTA |
| MALAT1-4-SPR | /5Phos/AAGATAAACATCGTAGACTATCAGACTCACCGGAATT<br>CGATCACC |
| MALAT1-5-SPL | /5Phos/TCTGCAGTTTCTATAGAAAAAACTCACTGATCAGGTC<br>ATACACTA |
| MALAT1-5-SPR | /5Phos/AAGATAAACATCGTAGACTATCAGACTCAAAGCCACT<br>TCCTTTGC |
| MALAT1-6-SPL | /5Phos/ACGTCCCCATATAAATAAAAAACTCACTGATCAGGTC<br>ATACACTA |
| MALAT1-6-SPR | /5Phos/AAGATAAACATCGTAGACTATCAGACTCAACCCGGAA<br>ATCGGCCT |
| Malat1-1-SPL | /5Phos/CCGACCTCAAGGAATGAAAAAACTCATCTCTCAGGTC<br>ATACACTA |
| Malat1-1-SPR | /5Phos/AAAGATAAACATCGTAGACTACTCTACTCAAACCACC<br>ACCATGTTG |
| Malat1-2-SPL | /5Phos/CGGCAACTGGGAAAACAAAAAACTCATCTCTCAGGTC<br>ATACACTA |
| Malat1-2-SPR | /5Phos/AAAGATAAACATCGTAGACTACTCTACTCAACTGTGC<br>TGACTTCAG |
| Malat1-3-SPL | /5Phos/GATTAAAGGCGTGCGCAAAAAACTCATCTCTCAGGTC<br>ATACACTA |
| Malat1-3-SPR | /5Phos/AAAGATAAACATCGTAGACTACTCTACTCAGCCTCCC<br>AAGTGCTAG |
| connected probe-1 | ACAAAATAGAACCGCGGTCCTATTCAAAAAAAAAAAAAAAAA<br>AAAAAAAAAAAAAAAAAAAAAAAAAAAAAAAATATCTTTAGT<br>GT |
| connected probe-2 | CATCGTTTATGGTCGGAACTACGACAAAAAAAAAAAAAAAAA<br>AAAAAAAAAAAAAAAAAAAAAAAAAAAAAAAATATCTTTAGT<br>GT |
| connected probe-3 | TGTTATTGCTCAATCTCGGGTGGCTAAAAAAAAAAAAAAAAAA<br>AAAAAAAAAAAAAAAAAAAAAAAAAAAAAAAATATCTTTAGTG<br>T |
| connected probe-4 | AGATAGTCAAGTTCGACCGTCTTCTAAAAAAAAAAAAAAAAAA<br>AAAAAAAAAAAAAAAAAAAAAAAAAAAAAAAATATCTTTAGTG<br>T |
| connected probe-5 | AGGTTTCCCGTGTTGAGTCAAATTAAAAAAAAAAAAAAAAAAA<br>AAAAAAAAAAAAAAAAAAAAAAAAAAAAAAATATCTTTAGTGT |
| connected probe without the 18S rRNA-binding domain | AAAAAAAAAAAAAAAAAAAAAAAAAAAAAAAAAAAAAAAAAAA<br>AAAAAAATATCTTTAGTGT |
| detection probe | /5Cy3/TCATACACTAAAGATAAACA |
| splint oligonucleotide-1 | TGTTTATCTTTAGTGTATGA |
| splint oligonucleotide-2 | ACCAAGTATCTTTAGTGTTTACGT |
| dapB-1-SPL | /5Phos/CGGAACTGCGCTTAAAAAAAAACTCAAGGATCAGGTC |

| | |
|---|---|
| | ATACACTA |
| dapB-1-SPR | /5Phos/AGATAAACATCGTAGACTAGAAGACTCAAGCTTGACG CACCAAG |
| dapB-2-SPL | /5Phos/GGAACAACCGGTTTCTAAAAAACTCAAGGATCAGGTC ATACACTA |
| dapB-2-SPR | /5Phos/AAGATAAACATCGTAGACTAGAAGACTCAAGTCCGTC CAGTTGTC |
| dapB-3-SPL | /5Phos/GCGTGGAAGAATGGGGAAAAAACTCAAGGATCAGGT CATACACTA |
| dapB-3-SPR | /5Phos/AAGATAAACATCGTAGACTAGAAGACTCATAGTCATT GCGGGACC |
| Ttr-1-SPL | /5Phos/GTTGCTGACGACAGCCAAAAAACTCATCAGTCAGGTC ATACACTA |
| Ttr-1-SPR | /5Phos/AAGATAAACATCGTAGACTAGACTACTCAGTCTCTCA ATTCTGGGG |
| Plp1-1-SPL | /5Phos/TGACAGCGCCGGTGGTAAAAAACTCATCCTTCAGGTC ATACACTA |
| Plp1-1-SPR | /5Phos/AAGATAAACATCGTAGACTAGAAGACTCATCGCCAAA GATCTGCC |
| Plp1-2-SPL | /5Phos/TGACAGGTGGTCCAGGAAAAAACTCATCCTTCAGGTC ATACACTA |
| Plp1-2-SPR | /5Phos/AAGATAAACATCGTAGACTAGAAGACTCAAGGGAAGG CAATAGAC |
| Plp1-3-SPL | /5Phos/ATTCTGGCATCAGCGCAAAAAACTCATCCTTCAGGTC ATACACTA |
| Plp1-3-SPR | /5Phos/AAGATAAACATCGTAGACTAGAAGACTCATGGGAGAA CACCATAC |
| Plp1-4-SPL | /5Phos/AGCAATCATGAAGGTGAAAAAACTCATCCTTCAGGTC ATACACTA |
| Plp1-4-SPR | /5Phos/AAGATAAACATCGTAGACTAGAAGACTCACGAAGTTG TAAGTGGC |
| Sox10-1-SPL | /5Phos/CTGCTGTTCCTTCTTGAAAAAACTCATCCTTCAGGTCA TACACTA |
| Sox10-1-SPR | /5Phos/AAGATAAACATCGTAGACTATCAGACTCACGTCCGCC TCGCCGTC |
| Sox10-2-SPL | /5Phos/CGGACTGCAGCTCTGTAAAAAACTCATCCTTCAGGTC ATACACTA |
| Sox10-2-SPR | /5Phos/AAGATAAACATCGTAGACTATCAGACTCATTGGGGTC TGCCTTGC |
| Sox10-3-SPL | /5Phos/CTCGTGGCTGATCTCCAAAAAACTCATCCTTCAGGTC ATACACTA |
| Sox10-3-SPR | /5Phos/AAGATAAACATCGTAGACTATCAGACTCACCATGTTG GACATTAC |
| Sox10-4-SPL | /5Phos/TGAGGGAAGTGTAGGCAAAAAACTCATCCTTCAGGTC |

| | |
|---|---|
| | ATACACTA |
| Sox10-4-SPR | /5Phos/AAGATAAACATCGTAGACTATCAGACTCACCGTAGTG GGGCAGAC |
| Gad2-1-SPL | /5Phos/CATCCCCTTTGGGGCAAAAAAACTCATCGATCAGGTC ATACACTA |
| Gad2-1-SPR | /5Phos/AAGATAAACATCGTAGACTACTAGACTCAAGAAACGC GTAGTTGA |
| Gad2-2-SPL | /5Phos/TTGCGGTTGGTCTGCCAAAAAACTCATCGATCAGGTC ATACACTA |
| Gad2-2-SPR | /5Phos/AAGATAAACATCGTAGACTACTAGACTCAAAATTTCCT CCAGATT |
| Gad2-3-SPL | /5Phos/TCCACCCCAAGCAGCAAAAAAACTCATCGATCAGGTC ATACACTA |
| Gad2-3-SPR | /5Phos/AAGATAAACATCGTAGACTACTAGACTCAGGGACATC AGTAACCC |
| Gad2-4-SPL | /5Phos/CCCATACTCCATCATTAAAAAACTCATCGATCAGGTC ATACACTA |
| Gad2-4-SPR | /5Phos/AAGATAAACATCGTAGACTACTAGACTCAAGCTGACC ATTGTGGT |
| Slc32a1-1-SPL | /5Phos/CGGGAGCTTCTGCGCCAAAAAACTCATCGATCAGGT CATACACTA |
| Slc32a1-1-SPR | /5Phos/AAGATAAACATCGTAGACTAAGCTACTCATGAATGTC TCCCTCGA |
| Slc32a1-2-SPL | /5Phos/CGTTCCAGCCCGCTTCAAAAAACTCATCGATCAGGTC ATACACTA |
| Slc32a1-2-SPR | /5Phos/AAGATAAACATCGTAGACTAAGCTACTCATGAATGGC ATTTGTCA |
| Slc32a1-3-SPL | /5Phos/CACGGGCAGCCCCGGGAAAAAACTCATCGATCAGGT CATACACTA |
| Slc32a1-3-SPR | /5Phos/AAGATAAACATCGTAGACTAAGCTACTCAACCAGGAC TTCTGCGA |
| Slc32a1-4-SPL | /5Phos/CGATGATGCCAATGGAAAAAAACTCATCGATCAGGTC ATACACTA |
| Slc32a1-4-SPR | /5Phos/AAGATAAACATCGTAGACTAAGCTACTCAGACGTGTA GCTGAACA |
| Tbr1-1-SPL | /5Phos/TGGACTCGGCCTGGCTAAAAAACTCATCAGTCAGGTC ATACACTA |
| Tbr1-1-SPR | /5Phos/AAGATAAACATCGTAGACTACTGAACTCACGGAAAAG TGAACGTC |
| Tbr1-2-SPL | /5Phos/CGGGTGGTGGGCCATGAAAAAACTCATCAGTCAGGT CATACACTA |
| Tbr1-2-SPR | /5Phos/AAGATAAACATCGTAGACTACTGAACTCACTCCGTTG GTAATGAC |
| Tbr1-3-SPL | /5Phos/ACATTGGTGTCCGCTTAAAAAACTCATCAGTCAGGTC |

| | |
|---|---|
| | ATACACTA |
| Tbr1-3-SPR | /5Phos/AAGATAAACATCGTAGACTACTGAACTCAGACCCGGT TTCCTTGC |
| Tbr1-4-SPL | /5Phos/TCAGATCCGCCCGAATAAAAAACTCATCAGTCAGGTC ATACACTA |
| Tbr1-4-SPR | /5Phos/AAGATAAACATCGTAGACTACTGAACTCAATCATGCA AGACAAGC |
| Mfge8-1-SPL | /5Phos/TTCCGCAGAAGGTTCAAAAAAACTCAGATCTCAGGTC ATACACTA |
| Mfge8-1-SPR | /5Phos/AAGATAAACATCGTAGACTAGAGAACTCAACCTGATA CCCGCATC |
| Mfge8-2-SPL | /5Phos/TGTGCCTCCAGAGTCGAAAAAACTCAGATCTCAGGTC ATACACTA |
| Mfge8-2-SPR | /5Phos/AAGATAAACATCGTAGACTAGAGAACTCAGTACAGCT TTATGTAC |
| Mfge8-3-SPL | /5Phos/CTTCCCAAGTGGGGGTAAAAAACTCAGATCTCAGGTC ATACACTA |
| Mfge8-3-SPR | /5Phos/AAGATAAACATCGTAGACTAGAGAACTCAGCCCTGAT TATCCAGC |
| Mfge8-4-SPL | /5Phos/TTGTAGGACGCCACATAAAAAACTCAGATCTCAGGTC ATACACTA |
| Mfge8-4-SPR | /5Phos/AAGATAAACATCGTAGACTAGAGAACTCAATCACTGT GGGCTACC |
| Aldoc-1-SPL | /5Phos/CTTATCCTGGATGGTGAAAAAACTCAGAAGTCAGGTC ATACACTA |
| Aldoc-1-SPR | /5Phos/AAGATAAACATCGTAGACTACTTCACTCATGCCTACG AGAATGCC |
| Aldoc-2-SPL | /5Phos/TAGAGGCACTACACCCAAAAAACTCAGAAGTCAGGTC ATACACTA |
| Aldoc-2-SPR | /5Phos/AAGATAAACATCGTAGACTACTTCACTCACCCCGTCG GTCCCAGC |
| Aldoc-3-SPL | /5Phos/ATGATGGTCACTCAGGAAAAAACTCAGAAGTCAGGTC ATACACTA |
| Aldoc-3-SPR | /5Phos/AAGATAAACATCGTAGACTACTTCACTCATCCCTTCG AGGTATAC |
| Flt1-1-SPL | /5Phos/AGTGTGCATCTCTATGAAAAAACTCAGAAGTCAGGTC ATACACTA |
| Flt1-1-SPR | /5Phos/AAGATAAACATCGTAGACTATCCTACTCACAAGTTTG GGTATGTC |
| Flt1-2-SPL | /5Phos/TGCCGATGGGTCAGATAAAAAACTCAGAAGTCAGGTC ATACACTA |
| Flt1-2-SPR | /5Phos/AAGATAAACATCGTAGACTATCCTACTCATAGGATTGT ATTGGTC |
| Flt1-3-SPL | /5Phos/CCCTGCATCCTCGGTTAAAAAACTCAGAAGTCAGGTC |

| | |
|---|---|
| | ATACACTA |
| Flt1-3-SPR | /5Phos/AAGATAAACATCGTAGACTATCCTACTCAGCAAGATC GTATAGTC |
| Vtn-1-SPL | /5Phos/TTCTTGCTGGCCATGAAAAAAACTCAGACTTCAGGTC ATACACTA |
| Vtn-1-SPR | /5Phos/AAGATAAACATCGTAGACTAAGTCACTCACTCGTCAC ACTGACAC |
| Vtn-2-SPL | /5Phos/TTTCCACTGCACAGTTAAAAAACTCAGACTTCAGGTC ATACACTA |
| Vtn-2-SPR | /5Phos/AAGATAAACATCGTAGACTAAGTCACTCAGAAGGCGT CAAAGGGC |
| Vtn-3-SPL | /5Phos/AGCATCGATGGGGCCCAAAAAACTCAGACTTCAGGT CATACACTA |
| Vtn-3-SPR | /5Phos/AAGATAAACATCGTAGACTAAGTCACTCATGATGCGA GTGAAGGC |
| Vtn-4-SPL | /5Phos/TTAGACTTCTGTTTTTAAAAAACTCAGACTTCAGGTCA TACACTA |
| Vtn-4-SPR | /5Phos/AAGATAAACATCGTAGACTAAGTCACTCACTTTCGGC TTCTACGC |
| Mrc1-1-SPL | /5Phos/CAAACAGCTTGTCTTTAAAAAACTCAGAGATCAGGTC ATACACTA |
| Mrc1-1-SPR | /5Phos/AAGATAAACATCGTAGACTAAGAGACTCATGCAATGG ACAAAATC |
| Mrc1-2-SPL | /5Phos/TCGTGGATCTCCGTGAAAAAAACTCAGAGATCAGGTC ATACACTA |
| Mrc1-2-SPR | /5Phos/AAGATAAACATCGTAGACTAAGAGACTCATGTGAGGT ACATTTGC |
| Mrc1-3-SPL | /5Phos/CAATCCAGAGTCCCGAAAAAAACTCAGAGATCAGGTC ATACACTA |
| Mrc1-3-SPR | /5Phos/AAGATAAACATCGTAGACTAAGAGACTCACTCAGACT GTTGAGTC |
| Mrc1-4-SPL | /5Phos/TGAGAGTCCTGTCCAAAAAAAACTCAGAGATCAGGTC ATACACTA |
| Mrc1-4-SPR | /5Phos/AAGATAAACATCGTAGACTAAGAGACTCACTTTGTTTT GAACATC |
| S100a8-1-SPL | /5Phos/AAGGCCTTCTCCAGTTAAAAAACTCAGACTTCAGGTC ATACACTA |
| S100a8-1-SPR | /5Phos/AAGATAAACATCGTAGACTAGAGAACTCAATCAATGA GGTTGCTC |
| S100a8-2-SPL | /5Phos/TGAGGACACTCAGTAGAAAAAACTCAGACTTCAGGTC ATACACTA |
| S100a8-2-SPR | /5Phos/AAGATAAACATCGTAGACTAGAGAACTCATATATTCTG CACAAAC |
| S100a8-3-SPL | /5Phos/CTCGAAGTTAATTGCAAAAAAACTCAGACTTCAGGTC |

| | |
|---|---|
| | ATACACTA |
| S100a8-3-SPR | /5Phos/AAGATAAACATCGTAGACTAGAGAACTCACCATCGCA AGGAACTC |
| S100a9-1-SPL | /5Phos/TGCGCTCCATCTGAGAAAAAAACTCAGACTTCAGGTC ATACACTA |
| S100a9-1-SPR | /5Phos/AAGATAAACATCGTAGACTAGATCACTCAATGATGGT GGTTATGC |
| S100a9-2-SPL | /5Phos/TGTGCTTCCACCATTTAAAAAACTCAGACTTCAGGTC ATACACTA |
| S100a9-2-SPR | /5Phos/AAGATAAACATCGTAGACTAGATCACTCACATAAAGG TTGCCAAC |
| S100a9-3-SPL | /5Phos/AGGCAAAGATCAACTTAAAAAACTCAGACTTCAGGTC ATACACTA |
| S100a9-3-SPR | /5Phos/AAGATAAACATCGTAGACTAGATCACTCATGCAGCTT CTCATGAC |
| Fcgr2b-1-SPL | /5Phos/TCCATGGGGATCAGTAAAAAAACTCAGACTTCAGGTC ATACACTA |
| Fcgr2b-1-SPR | /5Phos/AAGATAAACATCGTAGACTATCCTACTCAGACAGTCC AGTTGCTC |
| Fcgr2b-2-SPL | /5Phos/TGGCTTGGACCTGGCTAAAAAACTCAGACTTCAGGTC ATACACTA |
| Fcgr2b-2-SPR | /5Phos/AAGATAAACATCGTAGACTATCCTACTCAGCCTTAAA CGTGTAGC |
| Fcgr2b-3-SPL | /5Phos/GTGATGGTTTCCCCTTAAAAAACTCAGACTTCAGGTC ATACACTA |
| Fcgr2b-3-SPR | /5Phos/AAGATAAACATCGTAGACTATCCTACTCAGCTATGGC ACCTTAGC |
| Fcgr2b-4-SPL | /5Phos/CTGGGAGAGCTGGAACAAAAAACTCAGACTTCAGGT CATACACTA |
| Fcgr2b-4-SPR | /5Phos/AAGATAAACATCGTAGACTATCCTACTCACTGTGATC AGGGTTTC |
| Nod2-1-SPL | /5Phos/AGCCAGTCTAAGATGCAAAAAACTCAGACTTCAGGTC ATACACTA |
| Nod2-1-SPR | /5Phos/AAGATAAACATCGTAGACTATCAGACTCACACATCCC AAGACAGC |
| Nod2-2-SPL | /5Phos/TTCAGTCTGCAGCTCCAAAAAACTCAGACTTCAGGTC ATACACTA |
| Nod2-2-SPR | /5Phos/AAGATAAACATCGTAGACTATCAGACTCACCCCGGCT GTGCCCAC |
| Nod2-3-SPL | /5Phos/AGTTGCAACTCTGTGCAAAAAACTCAGACTTCAGGTC ATACACTA |
| Nod2-3-SPR | /5Phos/AAGATAAACATCGTAGACTATCAGACTCATTCAGAGA AGCCCTTC |
| Nod2-4-SPL | /5Phos/AAATGAAGCCCGGCATAAAAAACTCAGACTTCAGGTC |

| | |
|---|---|
| | ATACACTA |
| Nod2-4-SPR | /5Phos/AAGATAAACATCGTAGACTATCAGACTCAAGGCTACGGATGAGCC |
| Ptger4-1-SPL | /5Phos/AGCCAGCCCACATACTAAAAAACTCAGACTTCAGGTCATACACTA |
| Ptger4-1-SPR | /5Phos/AAGATAAACATCGTAGACTAAGTCACTCACCAGAAGGTCAGTGAC |
| Ptger4-2-SPL | /5Phos/AGAGTGTGAGGCCGGCAAAAAACTCAGACTTCAGGTCATACACTA |
| Ptger4-2-SPR | /5Phos/AAGATAAACATCGTAGACTAAGTCACTCAGATGCATAGATGGCGA |
| Ptger4-3-SPL | /5Phos/TCGCACCACGAGCGGAAAAAAACTCAGACTTCAGGTCATACACTA |
| Ptger4-3-SPR | /5Phos/AAGATAAACATCGTAGACTAAGTCACTCAACTGGTTAATGAACAC |
| Ptger4-4-SPL | /5Phos/TGATCTCCTTTAACTCAAAAAACTCAGACTTCAGGTCATACACTA |
| Ptger4-4-SPR | /5Phos/AAGATAAACATCGTAGACTAAGTCACTCAGTCTGGGACGTGCTGC |
| Hmgb1-1-SPL | /5Phos/ATCCGGGTGCTTCTTCAAAAAACTCAGATCTCAGGTCATACACTA |
| Hmgb1-1-SPR | /5Phos/AAGATAAACATCGTAGACTAAGCTACTCAAGAAGTTGACAGAAGC |
| Hmgb1-2-SPL | /5Phos/CTTGTCAGCCTTTGCCAAAAAACTCAGATCTCAGGTCATACACTA |
| Hmgb1-2-SPR | /5Phos/AAGATAAACATCGTAGACTAAGCTACTCACTCTTTCATAACGAGC |
| Hmgb1-3-SPL | /5Phos/CTTTGATTTTGGGGCGAAAAAACTCAGATCTCAGGTCATACACTA |
| Hmgb1-3-SPR | /5Phos/AAGATAAACATCGTAGACTAAGCTACTCAAAGCCAGGATGCTCGC |
| Hmgb1-4-SPL | /5Phos/CTTGACCACCCCCTTTAAAAAACTCAGATCTCAGGTCATACACTA |
| Hmgb1-4-SPR | /5Phos/AAGATAAACATCGTAGACTAAGCTACTCATCTTGCTCTTTTCAGC |
| Pf4-1-SPL | /5Phos/TCGAAACACCGCAGCGAAAAAACTCAGATCTCAGGTCATACACTA |
| Pf4-1-SPR | /5Phos/AAGATAAACATCGTAGACTACTTCACTCAGACTGGGCCGGAGGCC |
| Pf4-2-SPL | /5Phos/TTCGGGACCAGCGCTGAAAAAACTCAGATCTCAGGTCATACACTA |
| Pf4-2-SPR | /5Phos/AAGATAAACATCGTAGACTACTTCACTCAGATCTCCATCGCTTTC |
| Pf4-3-SPL | /5Phos/TCCCATTCTTCAGGGTAAAAAACTCAGATCTCAGGTC |

| | |
|---|---|
| | ATACACTA |
| Pf4-3-SPR | /5Phos/AAGATAAACATCGTAGACTACTTCACTCATCCAGGCA AATTTTCC |
| Ccl5-1-SPL | /5Phos/AGTGAGGATGATGGTGAAAAAACTCAAGGATCAGGT CATACACTA |
| Ccl5-1-SPR | /5Phos/AAGATAAACATCGTAGACTAAGCTACTCATGCAGAGG GCGGCTGC |
| Ccl5-2-SPL | /5Phos/TGCTGGTGTAGAAATAAAAAAACTCAAGGATCAGGTC ATACACTA |
| Ccl5-2-SPR | /5Phos/AAGATAAACATCGTAGACTAAGCTACTCAAGATTGGA GCACTTGC |
| Ccl5-3-SPL | /5Phos/TTGAACCCACTTCTTCAAAAAACTCAAGGATCAGGTC ATACACTA |
| Ccl5-3-SPR | /5Phos/AAGATAAACATCGTAGACTAAGCTACTCAAATAGTTG ATGTATTC |
| Ccl3-1-SPL | /5Phos/AGAGTGTCATGGTACAAAAAAACTCAAGGATCAGGTC ATACACTA |
| Ccl3-1-SPR | /5Phos/AAGATAAACATCGTAGACTAAGAGACTCAGAGAAGAC TTGGTTGC |
| Ccl3-2-SPL | /5Phos/CTGGCTGGGAGCAAAGAAAAAACTCAAGGATCAGGT CATACACTA |
| Ccl3-2-SPR | /5Phos/AAGATAAACATCGTAGACTAAGAGACTCAGTCAGGAA AATGACAC |
| Ccl3-3-SPL | /5Phos/AGTGATGTATTCTTGGAAAAAACTCAAGGATCAGGTC ATACACTA |
| Ccl3-3-SPR | /5Phos/AAGATAAACATCGTAGACTAAGAGACTCACATTCAGT TCCAGGTC |
| Ccl4-1-SPL | /5Phos/AAGAGGAGAGAGAGGGAAAAAACTCAAGGATCAGGT CATACACTA |
| Ccl4-1-SPR | /5Phos/AAGATAAACATCGTAGACTATCGAACTCAGAAGGCAG CCACGAGC |
| Ccl4-2-SPL | /5Phos/TGGCTTGGAGCAAAGAAAAAAACTCAAGGATCAGGTC ATACACTA |
| Ccl4-2-SPR | /5Phos/AAGATAAACATCGTAGACTATCGAACTCATCAGGAAT ACCACAGC |
| Slc6a3-1-SPL | /5Phos/AGCTGCACTCCATTCTAAAAAACTCATCAGTCAGGTC ATACACTA |
| Slc6a3-1-SPR | /5Phos/AAGATAAACATCGTAGACTACTAGACTCAGAGGGTGG AGTTGGTC |
| Slc6a3-2-SPL | /5Phos/ACCAGCAGCTCCTTCTAAAAAACTCATCAGTCAGGTC ATACACTA |
| Slc6a3-2-SPR | /5Phos/AAGATAAACATCGTAGACTACTAGACTCAGGCAGATC TTCCAGAC |
| Slc6a3-3-SPL | /5Phos/CCAGGAAGGAGAAGACAAAAAACTCATCAGTCAGGT |

| | |
|---|---|
| | CATACACTA |
| Slc6a3-3-SPR | /5Phos/AAGATAAACATCGTAGACTACTAGACTCATTCTGTGC CATGTACC |
| Slc6a3-4-SPL | /5Phos/CAGTACAGGTTGGGTCAAAAAACTCATCAGTCAGGTC ATACACTA |
| Slc6a3-4-SPR | /5Phos/AAGATAAACATCGTAGACTACTAGACTCACTTCCAGC ATAGCCGC |
| Ccl25-1-SPL | /5Phos/AGGCAACCAGGCAGGCAAAAAACTCAAGGATCAGGT CATACACTA |
| Ccl25-1-SPR | /5Phos/AAGATAAACATCGTAGACTATCTCACTCACAGGCCCC AACAAAAC |
| Ccl25-2-SPL | /5Phos/TCTTCACATTCATGTCAAAAAACTCAAGGATCAGGTC ATACACTA |
| Ccl25-2-SPR | /5Phos/AAGATAAACATCGTAGACTATCTCACTCAAAGATTCTC ATCGCCC |
| Ccl25-3-SPL | /5Phos/ACCATCCTGGGATGACAAAAAACTCAAGGATCAGGTC ATACACTA |
| Ccl25-3-SPR | /5Phos/AAGATAAACATCGTAGACTATCTCACTCACTTTCTGG GCATCATC |
| Cxcr1-1-SPL | /5Phos/CAGCAAGCTCAGAAGGAAAAAACTCAAGAGTCAGGT CATACACTA |
| Cxcr1-1-SPR | /5Phos/AAGATAAACATCGTAGACTATCGAACTCATCACCAGC GAGTTTCC |
| Cxcr1-2-SPL | /5Phos/CCAGATGCCCACGCACAAAAAACTCAAGAGTCAGGT CATACACTA |
| Cxcr1-2-SPR | /5Phos/AAGATAAACATCGTAGACTATCGAACTCAGAATCAAA GATAGACC |
| Cxcr1-3-SPL | /5Phos/AGCGGCAGGAGGAAGCAAAAAACTCAAGAGTCAGGT CATACACTA |
| Cxcr1-3-SPR | /5Phos/AAGATAAACATCGTAGACTATCGAACTCAGACCAGCA TAGTGAGC |
| Cxcr1-4-SPL | /5Phos/AAGTGGGCTCCTAAGAAAAAAACTCAAGAGTCAGGTC ATACACTA |
| Cxcr1-4-SPR | /5Phos/AAGATAAACATCGTAGACTATCGAACTCAGCAAGTAT CTTCAATC |
| Cxcr2-1-SPL | /5Phos/ATGGGACAGCATCTGGAAAAAACTCAAGAGTCAGGT CATACACTA |
| Cxcr2-1-SPR | /5Phos/AAGATAAACATCGTAGACTACTAGACTCAAGGTTCTC TGAGTGGC |
| Cxcr2-2-SPL | /5Phos/AGGGCAAAGAACAGGTAAAAAACTCAAGAGTCAGGT CATACACTA |
| Cxcr2-2-SPR | /5Phos/AAGATAAACATCGTAGACTACTAGACTCACCAGACAG GCAAGGTC |
| Cxcr2-3-SPL | /5Phos/ATGATGAGCAGCGGCAAAAAAACTCAAGAGTCAGGT |

| | |
|---|---|
| | CATACACTA |
| Cxcr2-3-SPR | /5Phos/AAGATAAACATCGTAGACTACTAGACTCACCCGTAGCAGAACAGC |
| Cxcr2-4-SPL | /5Phos/ATGGCGAAATTTCTGGAAAAAACTCAAGAGTCAGGTCATACACTA |
| Cxcr2-4-SPR | /5Phos/AAGATAAACATCGTAGACTACTAGACTCATGATCTTGAGAAGTCC |
| Il2-1-SPL | /5Phos/TTGAAGTGGGTGCGCTAAAAAACTCAAGTCTCAGGTCATACACTA |
| Il2-1-SPR | /5Phos/AAGATAAACATCGTAGACTAGATCACTCAGAGCTTGAAGTGGAGC |
| Il2-2-SPL | /5Phos/AACAGCTGCTCCAGGTAAAAAACTCAAGTCTCAGGTCATACACTA |
| Il2-2-SPR | /5Phos/AAGATAAACATCGTAGACTAGATCACTCACTCCTGTAGGTCCATC |
| Il2-3-SPL | /5Phos/CTGCTTGGGCAAGTAAAAAAAACTCAAGTCTCAGGTCATACACTA |
| Il2-3-SPR | /5Phos/AAGATAAACATCGTAGACTAGATCACTCACTTTCAATTCTGTGGC |
| Il2-4-SPL | /5Phos/TTTAGTTTTACAACAGAAAAAACTCAAGTCTCAGGTCATACACTA |
| Il2-4-SPR | /5Phos/AAGATAAACATCGTAGACTAGATCACTCATGTGTTGTCAGAGCCC |
| Cxcl5-1-SPL | /5Phos/CGAGATGGAACCGCTGAAAAAACTCAAGTCTCAGGTCATACACTA |
| Cxcl5-1-SPR | /5Phos/AAGATAAACATCGTAGACTAAGAGACTCACCATCCGCATGAATGG |
| Cxcl5-2-SPL | /5Phos/TATGACTGAGGAAGGGAAAAAACTCAAGTCTCAGGTCATACACTA |
| Cxcl5-2-SPR | /5Phos/AAGATAAACATCGTAGACTAAGAGACTCAGCAGCTCCGTTGCGGC |
| Cxcl5-3-SPL | /5Phos/ACCGTAGGGCACTGTGAAAAAACTCAAGTCTCAGGTCATACACTA |
| Cxcl5-3-SPR | /5Phos/AAGATAAACATCGTAGACTAAGAGACTCATTTAGCTATGACTTCC |
| Cxcl5-4-SPL | /5Phos/TGCCCAATATTTTCTGAAAAAACTCAAGTCTCAGGTCATACACTA |
| Cxcl5-4-SPR | /5Phos/AAGATAAACATCGTAGACTAAGAGACTCAGCTTTCTTTTTGTCAC |
| Cxcl10-1-SPL | /5Phos/AGCACTTGGGTTCATGAAAAAACTCAAGTCTCAGGTCATACACTA |
| Cxcl10-1-SPR | /5Phos/AAGATAAACATCGTAGACTATCGAACTCAGGCAGAAAATGACGGC |
| Cxcl10-2-SPL | /5Phos/CTTGAGTCCCACTCAGAAAAAACTCAAGTCTCAGGTC |

| | |
|---|---|
| | ATACACTA |
| Cxcl10-2-SPR | /5Phos/AAGATAAACATCGTAGACTATCGAACTCACTTGCGAG AGGGATCC |
| Cxcl10-3-SPL | /5Phos/TCACTGGCCCGTCATCAAAAAACTCAAGTCTCAGGTC ATACACTA |
| Cxcl10-3-SPR | /5Phos/AAGATAAACATCGTAGACTATCGAACTCACCTATGGC CCTCATTC |
| Cxcl10-4-SPL | /5Phos/CGGATTCAGACATCTCAAAAAACTCAAGTCTCAGGTC ATACACTA |
| Cxcl10-4-SPR | /5Phos/AAGATAAACATCGTAGACTATCGAACTCATGATGGTC TTAGATTC |
| Il1a-1-SPL | /5Phos/AGAACTGTAGTCTTCGAAAAAACTCAAGCTTCAGGTC ATACACTA |
| Il1a-1-SPR | /5Phos/AAGATAAACATCGTAGACTAAGGAACTCAAGAGATGG TCAATGGC |
| Il1a-2-SPL | /5Phos/CGTTGCTTGACGTTGCAAAAAACTCAAGCTTCAGGTC ATACACTA |
| Il1a-2-SPR | /5Phos/AAGATAAACATCGTAGACTAAGGAACTCATTCTTCAG AATCTTCC |
| Il1a-3-SPL | /5Phos/TGACGAGCTTCATCAGAAAAAACTCAAGCTTCAGGTC ATACACTA |
| Il1a-3-SPR | /5Phos/AAGATAAACATCGTAGACTAAGGAACTCAATGACAAA CTTCTGCC |
| Il1a-4-SPL | /5Phos/AGCAACACGGGCTGGTAAAAAACTCAAGCTTCAGGT CATACACTA |
| Il1a-4-SPR | /5Phos/AAGATAAACATCGTAGACTAAGGAACTCATTCTGGCA ACTCCTTC |
| Il18-1-SPL | /5Phos/CAGGTCTCCATTTTCTAAAAAACTCAAGCTTCAGGTC ATACACTA |
| Il18-1-SPR | /5Phos/AAGATAAACATCGTAGACTAGAAGACTCACAAAGTTG TCTGATTC |
| Il18-2-SPL | /5Phos/TGGGGTTCACTGGCACAAAAAACTCAAGCTTCAGGTC ATACACTA |
| Il18-2-SPR | /5Phos/AAGATAAACATCGTAGACTAGAAGACTCATATTATCA GTCTGGTC |
| Il18-3-SPL | /5Phos/GAGGGTAGACATTTTAAAAAAACTCAAGCTTCAGGTC ATACACTA |
| Il18-3-SPR | /5Phos/AAGATAAACATCGTAGACTAGAAGACTCATCTTGTTCT TACAGGA |
| Il18-4-SPL | /5Phos/ACGTTTCTGAAAGAATAAAAAACTCAAGCTTCAGGTC ATACACTA |
| Il18-4-SPR | /5Phos/AAGATAAACATCGTAGACTAGAAGACTCATGTTGTGT CCTGGAAC |
| Il16-1-SPL | /5Phos/TTGGCCCTTCATCAGCAAAAAACTCAAGCTTCAGGTC |

| | |
|---|---|
| | ATACACTA |
| Il16-1-SPR | /5Phos/AAGATAAACATCGTAGACTACTTCACTCAAGCCCAGG CCCTTTGC |
| Il16-2-SPL | /5Phos/TCAGGATTGTGTAGACAAAAAACTCAAGCTTCAGGTC ATACACTA |
| Il16-2-SPR | /5Phos/AAGATAAACATCGTAGACTACTTCACTCACCAGGGTC ACAGTGGC |
| Il16-3-SPL | /5Phos/AAGAGACTGAAGATGGAAAAAACTCAAGCTTCAGGTC ATACACTA |
| Il16-3-SPR | /5Phos/AAGATAAACATCGTAGACTACTTCACTCACAGGTGAT CTGGCCAC |
| Il16-4-SPL | /5Phos/CTTGGGTCCCGAGCTTAAAAAACTCAAGCTTCAGGTC ATACACTA |
| Il16-4-SPR | /5Phos/AAGATAAACATCGTAGACTACTTCACTCAGACAATCA CAGCTTGC |
| Il15-1-SPL | /5Phos/ATATATGGTTTCAAAAAAAAAACTCAAGCTTCAGGTCA TACACTA |
| Il15-1-SPR | /5Phos/AAGATAAACATCGTAGACTACTCTACTCAGATGGATG TATTCCTC |
| Il15-2-SPL | /5Phos/CAGCCTCAGTTAAAAAAAAAAACTCAAGCTTCAGGTC ATACACTA |
| Il15-2-SPR | /5Phos//5Phos/AAGATAAACATCGTAGACTACTCTACTCAATGA AGACATGAATGC |
| Il15-3-SPL | /5Phos/AGTTCATTGCAGTAACAAAAAACTCAAGCTTCAGGTC ATACACTA |
| Il15-3-SPR | /5Phos/AAGATAAACATCGTAGACTACTCTACTCAAATTCCAG GAGAAAGC |
| Il15-4-SPL | /5Phos/TCCTCACATTCCTTGCAAAAAACTCAAGCTTCAGGTC ATACACTA |
| Il15-4-SPR | /5Phos/AAGATAAACATCGTAGACTACTCTACTCAGGTTTTCTC CTCCAGC |
| Fdx1-1-SPL | /5Phos/TCGCCATCTCGGTTCTAAAAAACTCATCGATCAGGTC ATACACTA |
| Fdx1-1-SPR | /5Phos/AAGATAAACATCGTAGACTAGACTACTCACTTGGTCG TTAGCGTC |
| Fdx1-2-SPL | /5Phos/CAAACCCATCGATATCAAAAAACTCATCGATCAGGTC ATACACTA |
| Fdx1-2-SPR | /5Phos/AAGATAAACATCGTAGACTAGACTACTCAGTTCCCTC ACACGCAC |
| Fdx1-3-SPL | /5Phos/CAGGTCAAGCATGTCAAAAAAACTCATCGATCAGGTC ATACACTA |
| Fdx1-3-SPR | /5Phos/AAGATAAACATCGTAGACTAGACTACTCACTGTTAGT CCAAAAGC |
| Fdx1-4-SPL | /5Phos/CGGACATCCGCCACTGAAAAAACTCATCGATCAGGTC |

| | |
|---|---|
| | ATACACTA |
| Fdx1-4-SPR | /5Phos/AAGATAAACATCGTAGACTAGACTACTCACATGTCAA CAGACTGT |
| Lipt1-1-SPL | /5Phos/AGTTTGATTCCTTCCTAAAAAACTCATCCTTCAGGTCA TACACTA |
| Lipt1-1-SPR | /5Phos/AAGATAAACATCGTAGACTAGACTACTCAACTCTTCCT CCGAGCC |
| Lipt1-2-SPL | /5Phos/ACATCCAGCTGGGGTTAAAAAACTCATCCTTCAGGTC ATACACTA |
| Lipt1-2-SPR | /5Phos/AAGATAAACATCGTAGACTAGACTACTCACTTTTTGGT AGGCTGC |
| Lipt1-3-SPL | /5Phos/TGCAGCCACCGCACTCAAAAAACTCATCCTTCAGGTC ATACACTA |
| Lipt1-3-SPR | /5Phos/AAGATAAACATCGTAGACTAGACTACTCAGATGCGCC GCGTACTC |
| Lipt1-4-SPL | /5Phos/CGTATACCCACTCCCAAAAAAACTCATCCTTCAGGTC ATACACTA |
| Lipt1-4-SPR | /5Phos/AAGATAAACATCGTAGACTAGACTACTCAAACTTGGG AGTCCTGC |
| Lias-1-SPL | /5Phos/TCACACACTGTGTGGAAAAAAACTCATCAGTCAGGTC ATACACTA |
| Lias-1-SPR | /5Phos/AAGATAAACATCGTAGACTAGAAGACTCAGGGGCAC CGGGCTTCC |
| Lias-2-SPL | /5Phos/AATGGAGGGGGATTTCAAAAAACTCATCAGTCAGGTC ATACACTA |
| Lias-2-SPR | /5Phos/AAGATAAACATCGTAGACTAGAAGACTCAGGGCTCAT TGGCATCC |
| Lias-3-SPL | /5Phos/TCCACTGCTCTCAGATAAAAAACTCATCAGTCAGGTC ATACACTA |
| Lias-3-SPR | /5Phos/AAGATAAACATCGTAGACTAGAAGACTCAAGACAGAG CCACCTTC |
| Lias-4-SPL | /5Phos/ACGTCGGCCGCACGGAAAAAAACTCATCAGTCAGGT CATACACTA |
| Lias-4-SPR | /5Phos/AAGATAAACATCGTAGACTAGAAGACTCATAGAGTTA AACAGTCC |
| Dld-1-SPL | /5Phos/AGCCTCAATTGGTTGAAAAAAACTCAGATCTCAGGTC ATACACTA |
| Dld-1-SPR | /5Phos/AAGATAAACATCGTAGACTATCGAACTCACTATCACT GTCACGTC |
| Dld-2-SPL | /5Phos/CGTGGGCCATATGGTAAAAAAACTCAGATCTCAGGTC ATACACTA |
| Dld-2-SPR | /5Phos/AAGATAAACATCGTAGACTATCGAACTCAGATGCAAA ATCTTTTC |
| Dld-3-SPL | /5Phos/TGAACCCAGTTCCACAAAAAAACTCAGATCTCAGGTC |

| | |
|---|---|
| | ATACACTA |
| Dld-3-SPR | /5Phos/AAGATAAACATCGTAGACTATCGAACTCACAAGTCTTT GCCAAAC |
| Dld-4-SPL | /5Phos/AAAGGGGAATTTTCCAAAAAAACTCAGATCTCAGGTC ATACACTA |
| Dld-4-SPR | /5Phos/AAGATAAACATCGTAGACTATCGAACTCACTCTGCTG TTTGCAGC |
| Dlat-1-SPL | /5Phos/TACCGCAACGGACCGCAAAAAACTCAGAAGTCAGGT CATACACTA |
| Dlat-1-SPR | /5Phos/AAGATAAACATCGTAGACTAAGTCACTCATAGCGGGG GATCCCAC |
| Dlat-2-SPL | /5Phos/AGGAAGAACAATCTGCAAAAAACTCAGAAGTCAGGTC ATACACTA |
| Dlat-2-SPR | /5Phos/AAGATAAACATCGTAGACTAAGTCACTCATGGTTGGG GAGAGGGC |
| Dlat-3-SPL | /5Phos/TGACGTCAACCACATGAAAAAACTCAGAAGTCAGGTC ATACACTA |
| Dlat-3-SPR | /5Phos/AAGATAAACATCGTAGACTAAGTCACTCAGTACTGAC AGCAACAC |
| Dlat-4-SPL | /5Phos/TCGCTTTGGAGGCTAAAAAAAACTCAGAAGTCAGGTC ATACACTA |
| Dlat-4-SPR | /5Phos/AAGATAAACATCGTAGACTAAGTCACTCATGAAGTTTA CCCTCTC |
| Pdha1-1-SPL | /5Phos/TCTCTGGTGAGCACTGAAAAAACTCAGACTTCAGGTC ATACACTA |
| Pdha1-1-SPR | /5Phos/AAGATAAACATCGTAGACTACTTCACTCAGTACTTGA GCCCATCC |
| Pdha1-2-SPL | /5Phos/TCCAACGATGCCGTTGAAAAAACTCAGACTTCAGGTC ATACACTA |
| Pdha1-2-SPR | /5Phos/AAGATAAACATCGTAGACTACTTCACTCACCAGGGGC ACCTGAGC |
| Pdha1-3-SPL | /5Phos/TCACAGATAAAAATGCAAAAAACTCAGACTTCAGGTC ATACACTA |
| Pdha1-3-SPR | /5Phos/AAGATAAACATCGTAGACTACTTCACTCAGCCATAGC GGTTGTTC |
| Pdha1-4-SPL | /5Phos/TCATGCTGTGTCCATGAAAAAACTCAGACTTCAGGTC ATACACTA |
| Pdha1-4-SPR | /5Phos/AAGATAAACATCGTAGACTACTTCACTCACTTACTCCA GGGTCAC |
| Pdhb-1-SPL | /5Phos/CTCTTCAGCAGCCCGGAAAAAACTCACTCTTCAGGTC ATACACTA |
| Pdhb-1-SPR | /5Phos/AAGATAAACATCGTAGACTATCCTACTCACGAGCGGT GAAAACGC |
| Pdhb-2-SPL | /5Phos/CACAGGCCTCTGCTAAAAAAAACTCACTCTTCAGGTC |

| | |
|---|---|
| | ATACACTA |
| Pdhb-2-SPR | /5Phos/AAGATAAACATCGTAGACTATCCTACTCAGTCACCGT ATTTCTTC |
| Pdhb-3-SPL | /5Phos/AGCTGAGTTTATAACCAAAAAACTCACTCTTCAGGTC ATACACTA |
| Pdhb-3-SPR | /5Phos/AAGATAAACATCGTAGACTATCCTACTCATATAGTAAG TCTTGGC |
| Pdhb-4-SPL | /5Phos/TTCTAGGCAGTGGCCCAAAAAACTCACTCTTCAGGTC ATACACTA |
| Pdhb-4-SPR | /5Phos/AAGATAAACATCGTAGACTATCCTACTCAACAATACA GCTGCAGC |
| Dbt-1-SPL | /5Phos/AGTATGGCTCCTGTCTAAAAAACTCAGAGATCAGGTC ATACACTA |
| Dbt-1-SPR | /5Phos/AAGATAAACATCGTAGACTAGAAGACTCATTTTGGTG AGGGAGGC |
| Dbt-2-SPL | /5Phos/CTGTCACTGGCTCTGTAAAAAACTCAGAGATCAGGTC ATACACTA |
| Dbt-2-SPR | /5Phos/AAGATAAACATCGTAGACTAGAAGACTCAATTGCCTT CTGGAAGC |
| Dbt-3-SPL | /5Phos/CATGGCGATCTCAAATAAAAAACTCAGAGATCAGGTC ATACACTA |
| Dbt-3-SPR | /5Phos/AAGATAAACATCGTAGACTAGAAGACTCAGGAGGCG GTTCAGTTC |
| Dbt-4-SPL | /5Phos/CAAATCGGGGAAGAGCAAAAAACTCAGAGATCAGGT CATACACTA |
| Dbt-4-SPR | /5Phos/AAGATAAACATCGTAGACTAGAAGACTCAACGTCTCC TTTCTGGT |
| Gcsh-1-SPL | /5Phos/AGAAGGTCAGGGCGGTAAAAAACTCACTAGTCAGGT CATACACTA |
| Gcsh-1-SPR | /5Phos/AAGATAAACATCGTAGACTATCAGACTCACAGGGCCG CGGCGAGC |
| Gcsh-2-SPL | /5Phos/CGTTCCAATACCTTCCAAAAAACTCACTAGTCAGGTC ATACACTA |
| Gcsh-2-SPR | /5Phos/AAGATAAACATCGTAGACTATCAGACTCAAATTGCTG ATTCCCAC |
| Gcsh-3-SPL | /5Phos/ATCTTCGTAACAGGATAAAAAACTCACTAGTCAGGTC ATACACTA |
| Gcsh-3-SPR | /5Phos/AAGATAAACATCGTAGACTATCAGACTCATCTTGATCA GCCAACC |
| Dlst-1-SPL | /5Phos/CTTCTGGAAGGCAGAAAAAAAACTCACTGATCAGGTC ATACACTA |
| Dlst-1-SPR | /5Phos/AAGATAAACATCGTAGACTAAGGAACTCACTAGAGGG CAGTTCCC |
| Dlst-2-SPL | /5Phos/TCCAACAGCTTTCTCCAAAAAACTCACTGATCAGGTC |

| | |
|---|---|
| | ATACACTA |
| Dlst-2-SPR | /5Phos/AAGATAAACATCGTAGACTAAGGAACTCACTTCTGCAACCGCATC |
| Dlst-3-SPL | /5Phos/TGAGTGAGCACAGGTGAAAAAACTCACTGATCAGGTCATACACTA |
| Dlst-3-SPR | /5Phos/AAGATAAACATCGTAGACTAAGGAACTCAGGGCACTGGTGGCATC |
| Dlst-4-SPL | /5Phos/TGATGGTGAAGGTACCAAAAAACTCACTGATCAGGTCATACACTA |
| Dlst-4-SPR | /5Phos/AAGATAAACATCGTAGACTAAGGAACTCAAAAACTCCTCCGTTGC |
| Atp7a-1-SPL | /5Phos/ACGGTATTGGTTAAGAAAAAAACTCACTTCTCAGGTCATACACTA |
| Atp7a-1-SPR | /5Phos/AAGATAAACATCGTAGACTAGATCACTCAAGTAACAGTCAGGAAC |
| Atp7a-2-SPL | /5Phos/TGGAGGGAGAGCTGGCAAAAAACTCACTTCTCAGGTCATACACTA |
| Atp7a-2-SPR | /5Phos/AAGATAAACATCGTAGACTAGATCACTCATTCTGAAGAGATGAGC |
| Atp7a-3-SPL | /5Phos/AATCAGCACATCCATGAAAAAACTCACTTCTCAGGTCATACACTA |
| Atp7a-3-SPR | /5Phos/AAGATAAACATCGTAGACTAGATCACTCATGGTGGTTGCCAGCAC |
| Atp7a-4-SPL | /5Phos/AACTCGGCCTCAGGTTAAAAAACTCACTTCTCAGGTCATACACTA |
| Atp7a-4-SPR | /5Phos/AAGATAAACATCGTAGACTAGATCACTCACAGAATGTGTACAGCC |
| Atp7b-1-SPL | /5Phos/AGACTGGAGATCCTGTAAAAAACTCAAGAGTCAGGTCATACACTA |
| Atp7b-1-SPR | /5Phos/AAGATAAACATCGTAGACTAAGGAACTCAGTTCACAATGCCTTTC |
| Atp7b-2-SPL | /5Phos/CTGCCTCTTGGTTGCTAAAAAACTCAAGAGTCAGGTCATACACTA |
| Atp7b-2-SPR | /5Phos/AAGATAAACATCGTAGACTAAGGAACTCAGGCTGATAGGTAATGA |
| Atp7b-3-SPL | /5Phos/ATGGGCTTTGCTGGTGAAAAAACTCAAGAGTCAGGTCATACACTA |
| Atp7b-3-SPR | /5Phos/AAGATAAACATCGTAGACTAAGGAACTCACAGGATCGAACTTCAC |
| Atp7b-4-SPL | /5Phos/ACTTGAGCTGAAGAGAAAAAAACTCAAGAGTCAGGTCATACACTA |
| Atp7b-4-SPR | /5Phos/AAGATAAACATCGTAGACTAAGGAACTCATCGGGCTTTCTATAGC |
| Slc31a1-1-SPL | /5Phos/CGTGTGGTTCATACCCAAAAAACTCAAGTCTCAGGTC |

| | |
|---|---|
| | ATACACTA |
| Slc31a1-1-SPR | /5Phos/AAGATAAACATCGTAGACTACTTCACTCATGGTAATGTTGTCGTC |
| Slc31a1-2-SPL | /5Phos/CACGAAGGCTCCAGCCAAAAAACTCAAGTCTCAGGTCATACACTA |
| Slc31a1-2-SPR | /5Phos/AAGATAAACATCGTAGACTACTTCACTCACTAGTAAAAACACTGC |
| Slc31a1-3-SPL | /5Phos/TCAGCATCTGCTGCCCAAAAAACTCAAGTCTCAGGTCATACACTA |
| Slc31a1-3-SPR | /5Phos/AAGATAAACATCGTAGACTACTTCACTCAAGGAGGTGGGGGAAGC |
| Slc31a1-4-SPL | /5Phos/CACCACTGCCTTCTTCAAAAAACTCAAGTCTCAGGTCATACACTA |
| Slc31a1-4-SPR | /5Phos/AAGATAAACATCGTAGACTACTTCACTCAGCTCTGTGATGTCCAC |
| Slc31a2-1-SPL | /5Phos/ACGGCCTCATCTGAGAAAAAAACTCAAGGATCAGGTCATACACTA |
| Slc31a2-1-SPR | /5Phos/AAGATAAACATCGTAGACTACTGAACTCAGAAATCAAAGAGAAGC |
| Slc31a2-2-SPL | /5Phos/CAACCTTGATGCCCTCAAAAAACTCAAGGATCAGGTCATACACTA |
| Slc31a2-2-SPR | /5Phos/AAGATAAACATCGTAGACTACTGAACTCAAGCAATTTGGCTTTGC |
| Slc31a2-3-SPL | /5Phos/TATTGTCTGAAGTTGAAAAAAACTCAAGGATCAGGTCATACACTA |
| Slc31a2-3-SPR | /5Phos/AAGATAAACATCGTAGACTACTGAACTCACACCTGAGGCGGGTCC |
| Slc31a2-4-SPL | /5Phos/AGCCAGCATCACAAAGAAAAAACTCAAGGATCAGGTCATACACTA |
| Slc31a2-4-SPR | /5Phos/AAGATAAACATCGTAGACTACTGAACTCATGTTGTAGGACATGAC |
| Ddc-1-SPL | /5Phos/CGAAGATAGCCAGGCTAAAAAACTCATCAGTCAGGTCATACACTA |
| Ddc-1-SPR | /5Phos/AAGATAAACATCGTAGACTATCGAACTCAGGCAGGGATCAGGGGC |
| Ddc-2-SPL | /5Phos/TGATCAGATGTGTAAGAAAAAACTCATCAGTCAGGTCATACACTA |
| Ddc-2-SPR | /5Phos/AAGATAAACATCGTAGACTATCGAACTCATACAGAGGAATGCGCC |
| Ddc-3-SPL | /5Phos/CACACCATTCAGAAGAAAAAAACTCATCAGTCAGGTCATACACTA |
| Ddc-3-SPR | /5Phos/AAGATAAACATCGTAGACTATCGAACTCAAGGAATCTGCAAACTC |
| Ddc-4-SPL | /5Phos/GAGGGTCCTGGCGTACAAAAAACTCATCAGTCAGGT |

| | |
|---|---|
| | CATACACTA |
| Ddc-4-SPR | /5Phos/AAGATAAACATCGTAGACTATCGAACTCAGTGCAAAT TTCAAAGC |
| Th-1-SPL | /5Phos/TGAAGCCCTTGGGCTGAAAAAACTCATCAGTCAGGTC ATACACTA |
| Th-1-SPR | /5Phos/AAGATAAACATCGTAGACTAAGTCACTCATCTGAGAC GGCTCTTC |
| Th-2-SPL | /5Phos/AGTGAGGAGGGTTTTGAAAAAACTCATCAGTCAGGTC ATACACTA |
| Th-2-SPR | /5Phos/AAGATAAACATCGTAGACTAAGTCACTCATTTCAAAG CCCGAGAC |
| Th-3-SPL | /5Phos/CAGCTGTGGAATGCTGAAAAAACTCATCAGTCAGGTC ATACACTA |
| Th-3-SPR | /5Phos/AAGATAAACATCGTAGACTAAGTCACTCAAGTGAGAC ACATCCTC |
| Th-4-SPL | /5Phos/AAAGGCCCGGACCTCGAAAAAACTCATCAGTCAGGT CATACACTA |
| Th-4-SPR | /5Phos/AAGATAAACATCGTAGACTAAGTCACTCACTGCTGTG TCTGGGTC |
| Chat-1-SPL | /5Phos/TCGTTGGACGCCATTTAAAAAACTCAAGCTTCAGGTC ATACACTA |
| Chat-1-SPR | /5Phos/AAGATAAACATCGTAGACTAAGTCACTCAAGGCAGGC GTTCATCC |
| Chat-2-SPL | /5Phos/TCAGCCTTCTGGGAGCAAAAAACTCAAGCTTCAGGTC ATACACTA |
| Chat-2-SPR | /5Phos/AAGATAAACATCGTAGACTAAGTCACTCAGGGGAACA TTTCCACC |
| Chat-3-SPL | /5Phos/AATGGCCATGCCGGTTAAAAAACTCAAGCTTCAGGTC ATACACTA |
| Chat-3-SPR | /5Phos/AAGATAAACATCGTAGACTAAGTCACTCACCAGAAGA TGGTTGTC |
| Gria1-1-SPL | /5Phos/TCCACTGCTGCATGATAAAAAACTCAAGTCTCAGGTC ATACACTA |
| Gria1-1-SPR | /5Phos/AAGATAAACATCGTAGACTATCAGACTCACGAGCGTC ACTTGTCC |
| Gria1-2-SPL | /5Phos/AAAGCGCACCTGCTGCAAAAAACTCAAGTCTCAGGTC ATACACTA |
| Gria1-2-SPR | /5Phos/AAGATAAACATCGTAGACTATCAGACTCATTCCTGTC AAACCTTC |
| Gria1-3-SPL | /5Phos/AGGTCCTCTGCACTCTAAAAAACTCAAGTCTCAGGTC ATACACTA |
| Gria1-3-SPR | /5Phos/AAGATAAACATCGTAGACTATCAGACTCATTCCGTCT GCTTTGCC |
| Rbfox3-1-SPL | /5Phos/AGAGGGTCATGCCGTGAAAAAACTCAAGGATCAGGT |

| | |
|---|---|
| | CATACACTA |
| Rbfox3-1-SPR | /5Phos/AAGATAAACATCGTAGACTAGATCACTCAGTCTGTGC GGGTGTGT |
| Rbfox3-2-SPL | /5Phos/CGCAGGTCGGGGTCCCAAAAAACTCAAGGATCAGGT CATACACTA |
| Rbfox3-2-SPR | /5Phos/AAGATAAACATCGTAGACTAGATCACTCATTGCCCGA ACATTTGC |
| Rbfox3-3-SPL | /5Phos/ACGACCGCTCCATAAGAAAAAACTCAAGGATCAGGTC ATACACTA |
| Rbfox3-3-SPR | /5Phos/AAGATAAACATCGTAGACTAGATCACTCAAAATCCAT CCTGATAC |
| Dcx-1-SPL | /5Phos/AGTGGGCACTATGAGTAAAAAACTCAAGAGTCAGGTC ATACACTA |
| Dcx-1-SPR | /5Phos/AAGATAAACATCGTAGACTATCAGACTCAGTTCTGTA GAAGCTAC |
| Dcx-2-SPL | /5Phos/TTGGGGTTGACATTCTAAAAAACTCAAGAGTCAGGTC ATACACTA |
| Dcx-2-SPR | /5Phos/AAGATAAACATCGTAGACTATCAGACTCATACGTTGA CAGACCAG |
| Dcx-3-SPL | /5Phos/TGGGGAAGCCTTTGGGAAAAAACTCAAGAGTCAGGT CATACACTA |
| Dcx-3-SPR | /5Phos/AAGATAAACATCGTAGACTATCAGACTCAATGTCTTTT GAGGTGT |
| Tph2-1-SPL | /5Phos/TTTTCTTGTCCTCGCTAAAAAACTCAAGAGTCAGGTC ATACACTA |
| Tph2-1-SPR | /5Phos/AAGATAAACATCGTAGACTAAGAGACTCACCGGGCTC TTTGCCGC |
| Tph2-2-SPL | /5Phos/CGGATTCAGGGTCACAAAAAAACTCAAGAGTCAGGTC ATACACTA |
| Tph2-2-SPR | /5Phos/AAGATAAACATCGTAGACTAAGAGACTCATCCAAATG CTCTCAGG |
| Tph2-3-SPL | /5Phos/TAAGCCTATCTCTTGGAAAAAACTCAAGAGTCAGGTC ATACACTA |
| Tph2-3-SPR | /5Phos/AAGATAAACATCGTAGACTAAGAGACTCAAGGCTCCC AGAGACGC |
| Pecam1-1-SPL | /5Phos/ACTAACACGTGGTCCTAAAAAACTCACTAGTCAGGTC ATACACTA |
| Pecam1-1-SPR | /5Phos/AAGATAAACATCGTAGACTACTCTACTCAAGCTTGGC AGCGAAAC |
| Pecam1-2-SPL | /5Phos/ATTGGAGGTCACCTCGAAAAAACTCACTAGTCAGGTC ATACACTA |
| Pecam1-2-SPR | /5Phos/AAGATAAACATCGTAGACTACTCTACTCATGAATGTTG CTGGGTC |
| Pecam1-3-SPL | /5Phos/ACGGGTTTCTGTTTGGAAAAAACTCACTAGTCAGGTC |

| | |
|---|---|
| | ATACACTA |
| Pecam1-3-SPR | /5Phos/AAGATAAACATCGTAGACTACTCTACTCATGGCCTGG ACATCTCC |
| Pdgfra-1-SPL | /5Phos/TTCTTCAGACATGGGGAAAAAACTCAAGAGTCAGGTC ATACACTA |
| Pdgfra-1-SPR | /5Phos/AAGATAAACATCGTAGACTATCTCACTCACCACGTTG GGGTCGTC |
| Pdgfra-2-SPL | /5Phos/AGCTCAAACGTGTAACAAAAAACTCAAGAGTCAGGTC ATACACTA |
| Pdgfra-2-SPR | /5Phos/AAGATAAACATCGTAGACTATCTCACTCAAGGAACTA GGGTTGAC |
| Pdgfra-3-SPL | /5Phos/TCTGGCCAGGCCAAAAAAAAAACTCAAGAGTCAGGTC ATACACTA |
| Pdgfra-3-SPR | /5Phos/AAGATAAACATCGTAGACTATCTCACTCAAATCGTGC ATGATGTC |
| Amigo2-1-SPL | /5Phos/TGAGGCGCTGGGGCTCAAAAAACTCATCTCTCAGGT CATACACTA |
| Amigo2-1-SPR | /5Phos/AAGATAAACATCGTAGACTATCAGACTCACAGTGGGG CACATTCC |
| Amigo2-2-SPL | /5Phos/AGCCAGCTTGAACCTCAAAAAACTCATCTCTCAGGTC ATACACTA |
| Amigo2-2-SPR | /5Phos/AAGATAAACATCGTAGACTATCAGACTCACTAAAAAT GTCAGATC |
| Amigo2-3-SPL | /5Phos/TGGCCACGCAGGCAGCAAAAAACTCATCTCTCAGGT CATACACTA |
| Amigo2-3-SPR | /5Phos/AAGATAAACATCGTAGACTATCAGACTCAAGCACTAG AACTATAC |
| Wfs1-1-SPL | /5Phos/AGGTTGGTGGGAATGAAAAAAACTCATCAGTCAGGTC ATACACTA |
| Wfs1-1-SPR | /5Phos/AAGATAAACATCGTAGACTAAGCTACTCACTCATCCT GCAGGAAC |
| Wfs1-2-SPL | /5Phos/AGAAGAGCAGGATGGCAAAAAACTCATCAGTCAGGT CATACACTA |
| Wfs1-2-SPR | /5Phos/AAGATAAACATCGTAGACTAAGCTACTCATACACGTA GAACCAGC |
| Wfs1-3-SPL | /5Phos/TCGCTGCTGGCACGGAAAAAAACTCATCAGTCAGGT CATACACTA |
| Wfs1-3-SPR | /5Phos/AAGATAAACATCGTAGACTAAGCTACTCACAGCACGT CCTTGAAC |
| Mbp-1-SPL | /5Phos/TGTGGCCAGGTACTTGAAAAAACTCAAGCTTCAGGTC ATACACTA |
| Mbp-1-SPR | /5Phos/AAGATAAACATCGTAGACTATCGAACTCAGGTCCATG GTACTTGC |
| Mbp-2-SPL | /5Phos/TAAAGAAGCGCCCGATAAAAAACTCAAGCTTCAGGTC |

| | |
|---|---|
| | ATACACTA |
| Mbp-2-SPR | /5Phos/AAGATAAACATCGTAGACTATCGAACTCAGCACCCCT GTCACCGC |
| Mbp-3-SPL | /5Phos/TGTGGCCAGGTACTTGAAAAAACTCAAGCTTCAGGTC ATACACTA |
| Mbp-3-SPR | /5Phos/AAGATAAACATCGTAGACTATCGAACTCAGGTCCATG GTACTTGC |
| Mbp-4-SPL | /5Phos/TGTGGCCAGGTACTTGAAAAAACTCAAGCTTCAGGTC ATACACTA |
| Mbp-4-SPR | /5Phos/AAGATAAACATCGTAGACTATCGAACTCAGGTCCATG GTACTTGC |
| Gfap-1-SPL | /5Phos/ATGGTACCCAGTTGTCAAAAAACTCAAGCTTCAGGTC ATACACTA |
| Gfap-1-SPR | /5Phos/AAGATAAACATCGTAGACTAGATCACTCACAAGGAGA AGCGTGGC |
| Gfap-2-SPL | /5Phos/CACACGAGCCAGGGTGAAAAAACTCAAGCTTCAGGT CATACACTA |
| Gfap-2-SPR | /5Phos/AAGATAAACATCGTAGACTAGATCACTCACCTTTCTCT CCAAATC |
| Gfap-3-SPL | /5Phos/TCCTCTGTCTCTTGCAAAAAAACTCAAGCTTCAGGTC ATACACTA |
| Gfap-3-SPR | /5Phos/AAGATAAACATCGTAGACTAGATCACTCACTTAGACC GATACCAC |
| Gfap-4-SPL | /5Phos/CTTTTGAGAGGTCTTGAAAAAACTCAAGCTTCAGGTC ATACACTA |
| Gfap-4-SPR | /5Phos/AAGATAAACATCGTAGACTAGATCACTCAAACTTGTAT TGTGAGC |
| Npy-1-SPL | /5Phos/ATAGAGCGAGGGTCAGAAAAAACTCAAGTCTCAGGT CATACACTA |
| Npy-1-SPR | /5Phos/AAGATAAACATCGTAGACTACTAGACTCAAAACACAC GAGCAGAG |
| Npy-2-SPL | /5Phos/CATGTCCTCTGCTGGCAAAAAACTCAAGTCTCAGGTC ATACACTA |
| Npy-2-SPR | /5Phos/AAGATAAACATCGTAGACTACTAGACTCACGGAGTAG TATCTGGC |
| Npy-3-SPL | /5Phos/TCAGGGCTGGATCTCTAAAAAACTCAAGTCTCAGGTC ATACACTA |
| Npy-3-SPR | /5Phos/AAGATAAACATCGTAGACTACTAGACTCAGTCTGAAA TCAGTGTC |
| Npy-4-SPL | /5Phos/CTTGTTCTGGGGGCGTAAAAAACTCAAGTCTCAGGTC ATACACTA |
| Npy-4-SPR | /5Phos/AAGATAAACATCGTAGACTACTAGACTCAGGAAGGGT CTTCAAGC |
| Sst-1-SPL | /5Phos/TGGGGTCCGAGGGCGCAAAAAACTCAAGTCTCAGGT |

| | |
|---|---|
| | CATACACTA |
| Sst-1-SPR | /5Phos/AAGATAAACATCGTAGACTAAGCTACTCAAGAAACTG ACGGAGTC |
| Sst-2-SPL | /5Phos/TCGGACAGCAGCTCTGAAAAAACTCAAGTCTCAGGTC ATACACTA |
| Sst-2-SPR | /5Phos/AAGATAAACATCGTAGACTAAGCTACTCACTCTGTCT GGTTGGGC |
| Sst-3-SPL | /5Phos/AGACCTCTGCAGCTCCAAAAAACTCAAGTCTCAGGTC ATACACTA |
| Sst-3-SPR | /5Phos/AAGATAAACATCGTAGACTAAGCTACTCACTGGGTTC GAGTTGGC |
| Sst-4-SPL | /5Phos/TAACAGGATGTGAATGAAAAAACTCAAGTCTCAGGTC ATACACTA |
| Sst-4-SPR | /5Phos/AAGATAAACATCGTAGACTAAGCTACTCAACAACAAT ATTAAAGC |
| Gad1-1-SPL | /5Phos/TTGGAGGACTGCCTCTAAAAAACTCAAGGATCAGGTC ATACACTA |
| Gad1-1-SPR | /5Phos/AAGATAAACATCGTAGACTACTTCACTCAACAGGAAA GCAGGTTC |
| Gad1-2-SPL | /5Phos/TTGCGGACATAGTTGAAAAAAACTCAAGGATCAGGTC ATACACTA |
| Gad1-2-SPR | /5Phos/AAGATAAACATCGTAGACTACTTCACTCAGGAGCGAT CAAATGTC |
| Gad1-3-SPL | /5Phos/ACCAGCTAAACCAATGAAAAAACTCAAGGATCAGGTC ATACACTA |
| Gad1-3-SPR | /5Phos/AAGATAAACATCGTAGACTACTTCACTCATCGATGTC AGCCATTC |
| Gad1-4-SPL | /5Phos/TGATGGCATATTGGTAAAAAAACTCAAGGATCAGGTC ATACACTA |
| Gad1-4-SPR | /5Phos/AAGATAAACATCGTAGACTACTTCACTCAAACACTCC CTCATGTC |
| Pvalb-1-SPL | /5Phos/TTGGATGAGCAGAGGCAAAAAACTCAAGGATCAGGT CATACACTA |
| Pvalb-1-SPR | /5Phos/AAGATAAACATCGTAGACTATCCTACTCAATCGACAT CCTGCAAC |
| Pvalb-2-SPL | /5Phos/TTCTTCACCTCATCCGAAAAAACTCAAGGATCAGGTC ATACACTA |
| Pvalb-2-SPR | /5Phos/AAGATAAACATCGTAGACTATCCTACTCACAGAATAT GGAACACC |
| Pvalb-3-SPL | /5Phos/TGGCATCTGAGGAGAAAAAAAACTCAAGGATCAGGTC ATACACTA |
| Pvalb-3-SPR | /5Phos/AAGATAAACATCGTAGACTATCCTACTCATTAGCAGA CAAGTCTC |
| Pvalb-4-SPL | /5Phos/TCCAGCGGCCAGAAGCAAAAAACTCAAGGATCAGGT |

| | |
|---|---|
| | CATACACTA |
| Pvalb-4-SPR | /5Phos/AAGATAAACATCGTAGACTATCCTACTCACGTCCCCA TCCTTGTC |
| Slc17a6-1-SPL | /5Phos/CGCGATGCGATATATCAAAAAACTCAAGAGTCAGGTC ATACACTA |
| Slc17a6-1-SPR | /5Phos/AAGATAAACATCGTAGACTAGATCACTCACCGGTTAG CAGCCAGC |
| Slc17a6-2-SPL | /5Phos/CAGAAGCAGCGTGGCTAAAAAACTCAAGAGTCAGGT CATACACTA |
| Slc17a6-2-SPR | /5Phos/AAGATAAACATCGTAGACTAGATCACTCAGAGAGTAG CCAACAAC |
| Slc17a6-3-SPL | /5Phos/AATGAGGAAGACATACAAAAAACTCAAGAGTCAGGTC ATACACTA |
| Slc17a6-3-SPR | /5Phos/AAGATAAACATCGTAGACTAGATCACTCAAGTGGACG AGTGCAGC |
| Slc17a6-4-SPL | /5Phos/CGTAAGATTTGGTGGTAAAAAACTCAAGAGTCAGGTC ATACACTA |
| Slc17a6-4-SPR | /5Phos/AAGATAAACATCGTAGACTAGATCACTCATCCTGTGA GGTAGCAC |
| Arc-1-SPL | /5Phos/CAGCGCTGTGAGTCGCAAAAAACTCACTAGTCAGGT CATACACTA |
| Arc-1-SPR | /5Phos/AAGATAAACATCGTAGACTATCGAACTCACTTGATGG ACTTCTTC |
| Arc-2-SPL | /5Phos/TCCTTCTTGAACTCCAAAAAAACTCACTAGTCAGGTC ATACACTA |
| Arc-2-SPR | /5Phos/AAGATAAACATCGTAGACTATCGAACTCAACTGTATT GCAGAAAC |
| Arc-3-SPL | /5Phos/TGCTTCTGCGGCAGCTAAAAAACTCACTAGTCAGGTC ATACACTA |
| Arc-3-SPR | /5Phos/AAGATAAACATCGTAGACTATCGAACTCAGTCGAGTG GTTCACCC |
| Arc-4-SPL | /5Phos/TGGTAAGAGCAGGTGTAAAAAACTCACTAGTCAGGTC ATACACTA |
| Arc-4-SPR | /5Phos/AAGATAAACATCGTAGACTATCGAACTCACTGGCTAC TGACTCGC |
| Homer1-1-SPL | /5Phos/ATGAGCTCGAGTGCTGAAAAAACTCACTAGTCAGGTC ATACACTA |
| Homer1-1-SPR | /5Phos/AAGATAAACATCGTAGACTAGATCACTCAGGTCAATC TGGAAGAC |
| Homer1-2-SPL | /5Phos/GTGTTTGCCCGGCTATAAAAAACTCACTAGTCAGGTC ATACACTA |
| Homer1-2-SPR | /5Phos/AAGATAAACATCGTAGACTAGATCACTCAATCCCAGT CCATAAACA |
| Homer1-3-SPL | /5Phos/TTTTCTGCGAATTTTGAAAAAACTCACTAGTCAGGTCA |

| | |
|---|---|
| | TACACTA |
| Homer1-3-SPR | /5Phos/AAGATAAACATCGTAGACTACTTCACTCACTTAAATTC CTGAAAC |
| Homer1-4-SPL | /5Phos/ATCGTCTGTCCCATTGAAAAAACTCACTAGTCAGGTC ATACACTA |
| Homer1-4-SPR | /5Phos/AAGATAAACATCGTAGACTAGATCACTCACATCGGGT GTTCTCTC |
| Nptx2-1-SPL | /5Phos/TGTCCTGGGCTCGTCCAAAAAACTCACTGATCAGGTC ATACACTA |
| Nptx2-1-SPR | /5Phos/AAGATAAACATCGTAGACTAAGTCACTCACTGCCAGG TATCGGGC |
| Nptx2-2-SPL | /5Phos/CGGATGGCTTCTCGCTAAAAAACTCACTGATCAGGTC ATACACTA |
| Nptx2-2-SPR | /5Phos/AAGATAAACATCGTAGACTAAGTCACTCACTTGCCGG TGAGCTCT |
| Nptx2-3-SPL | /5Phos/GAATGGCGTACCGATGAAAAAACTCACTGATCAGGTC ATACACTA |
| Nptx2-3-SPR | /5Phos/AAGATAAACATCGTAGACTAAGTCACTCACGGGCACA GCGTAGGA |
| Nptx2-4-SPL | /5Phos/TGCCTCCCACCGTGTCAAAAAACTCACTGATCAGGTC ATACACTA |
| Nptx2-4-SPR | /5Phos/AAGATAAACATCGTAGACTAAGTCACTCATGCGTGGC ATCAAATC |
| Bdnf-1-SPL | /5Phos/CGTGGACGTTTACTTCAAAAAACTCACTGATCAGGTC ATACACTA |
| Bdnf-1-SPR | /5Phos/AAGATAAACATCGTAGACTATCAGACTCAGCCAAGTT GCCTTGTC |
| Bdnf-2-SPL | /5Phos/CTGCCCTGGGCCCATTAAAAAACTCACTGATCAGGTC ATACACTA |
| Bdnf-2-SPR | /5Phos/AAGATAAACATCGTAGACTATCAGACTCAGTCAGACC TCTCGAAC |
| Bdnf-3-SPL | /5Phos/CGTCCCGCCAGACATGAAAAAACTCACTGATCAGGTC ATACACTA |
| Bdnf-3-SPR | /5Phos/AAGATAAACATCGTAGACTATCAGACTCATCTCTAGG ACTGTGAC |
| Bdnf-4-SPL | /5Phos/CAGTGCCTTTTGTCTAAAAAAACTCACTGATCAGGTC ATACACTA |
| Bdnf-4-SPR | /5Phos/AAGATAAACATCGTAGACTATCAGACTCATCGGCATT GCGAGTTC |
| Npas4-1-SPL | /5Phos/CTCGAATAAGCACCAGAAAAAACTCACTTCTCAGGTC ATACACTA |
| Npas4-1-SPR | /5Phos/AAGATAAACATCGTAGACTAAGGAACTCATGAGCATG GAATCGAC |
| Npas4-2-SPL | /5Phos/AGGGAATGGCAGGTAAAAAAAACTCACTTCTCAGGTC |

| | |
|---|---|
| | ATACACTA |
| Npas4-2-SPR | /5Phos/AAGATAAACATCGTAGACTAAGGAACTCAAAGGCTCA GGCCTAGC |
| Npas4-3-SPL | /5Phos/TGGGTGTCAACTGTTCAAAAAACTCACTTCTCAGGTC ATACACTA |
| Npas4-3-SPR | /5Phos/AAGATAAACATCGTAGACTAAGGAACTCAGGGAAGGT AGCACTGC |
| Npas4-4-SPL | /5Phos/AGGGGGTCCAGCCCCTAAAAAACTCACTTCTCAGGT CATACACTA |
| Npas4-4-SPR | /5Phos/AAGATAAACATCGTAGACTAAGGAACTCAGATCCAGG CTAAGCAC |
| Fos-1-SPL | /5Phos/CGAGAACATCATGGTCAAAAAACTCACTTCTCAGGTC ATACACTA |
| Fos-1-SPR | /5Phos/AAGATAAACATCGTAGACTAGAAGACTCAAGTCGGCG TTGAAACC |
| Fos-2-SPL | /5Phos/AGATCTGCGCAAAAGTAAAAAACTCACTTCTCAGGTC ATACACTA |
| Fos-2-SPR | /5Phos/AAGATAAACATCGTAGACTAGAAGACTCAGGCACTAG AGACGGAC |
| Fos-3-SPL | /5Phos/TGGCGTAAGCCCCAGCAAAAAACTCACTTCTCAGGTC ATACACTA |
| Fos-3-SPR | /5Phos/AAGATAAACATCGTAGACTAGAAGACTCATTCACCAT TCCCGCTC |
| Fos-4-SPL | /5Phos/CGTTTCTCTTCCTCTTAAAAAACTCACTTCTCAGGTCA TACACTA |
| Fos-4-SPR | /5Phos/AAGATAAACATCGTAGACTAGAAGACTCATTCCCTTC GGATTCTC |
| Fosb-1-SPL | /5Phos/GTAGTCTCCGGGAAAAAAAAAACTCACTCTTCAGGTC ATACACTA |
| Fosb-1-SPR | /5Phos/AAGATAAACATCGTAGACTAAGGAACTCAACCGGGAG CCGGAGTC |
| Fosb-2-SPL | /5Phos/TGAGACTCGGCGGAGGAAAAAACTCACTCTTCAGGT CATACACTA |
| Fosb-2-SPR | /5Phos/AAGATAAACATCGTAGACTAAGGAACTCACACCGAAG ACAGGTAC |
| Fosb-3-SPL | /5Phos/TCGGGGTCTTCTAGGCAAAAAACTCACTCTTCAGGTC ATACACTA |
| Fosb-3-SPR | /5Phos/AAGATAAACATCGTAGACTAAGGAACTCAGGGTAAGT GTCTCTTC |
| Fosb-4-SPL | /5Phos/TGTCAGCTCCCTCCGAAAAAAACTCACTCTTCAGGTC ATACACTA |
| Fosb-4-SPR | /5Phos/AAGATAAACATCGTAGACTAAGGAACTCACCGCCTGA AGTCGATC |
| Egr1-1-SPL | /5Phos/TCCACCATCGCCTTCTAAAAAACTCAGACTTCAGGTC |

| | |
|---|---|
| | ATACACTA |
| Egr1-1-SPR | /5Phos/AAGATAAACATCGTAGACTAAGAGACTCAGCTGGGATAACTCGTC |
| Egr1-2-SPL | /5Phos/CGAGTAGATGGGACTGAAAAAACTCAGACTTCAGGTCATACACTA |
| Egr1-2-SPR | /5Phos/AAGATAAACATCGTAGACTAAGAGACTCAGAAAGGTGGGCGCAGC |
| Egr1-3-SPL | /5Phos/CACTGGGGATGGGTAAAAAAAACTCAGACTTCAGGTCATACACTA |
| Egr1-3-SPR | /5Phos/AAGATAAACATCGTAGACTAAGAGACTCAATGGGTAGGAGGTAGC |
| Aqp4-1-SPL | /5Phos/TGAGTCCAAAGCAAAGAAAAAACTCATCAGTCAGGTCATACACTA |
| Aqp4-1-SPR | /5Phos/AAGATAAACATCGTAGACTAGAGAACTCAACCATGGTAGCAATGC |
| Aqp4-2-SPL | /5Phos/CGTGGTGACTCCCAATAAAAAACTCATCAGTCAGGTCATACACTA |
| Aqp4-2-SPR | /5Phos/AAGATAAACATCGTAGACTAGAGAACTCATGAGGTTTCCATGAAC |
| Aqp4-3-SPL | /5Phos/AGTTTCCCATGATAACAAAAAACTCATCAGTCAGGTCATACACTA |
| Aqp4-3-SPR | /5Phos/AAGATAAACATCGTAGACTAGAGAACTCAATCCAGTGGTTTGCCC |
| Plppr1-1-SPL | /5Phos/CTGACAGAAGAATCCTAAAAAACTCATCCTTCAGGTCATACACTA |
| Plppr1-1-SPR | /5Phos/AAGATAAACATCGTAGACTAAGGAACTCATCATTAAGTCTCCGTC |
| Plppr1-2-SPL | /5Phos/AGTGACGACTTGCCCCAAAAAACTCATCCTTCAGGTCATACACTA |
| Plppr1-2-SPR | /5Phos/AAGATAAACATCGTAGACTAAGGAACTCAATGGTGTTAGGTGACC |
| Plppr1-3-SPL | /5Phos/TGTCAGGAAGGCGGTAAAAAAACTCATCCTTCAGGTCATACACTA |
| Plppr1-3-SPR | /5Phos/AAGATAAACATCGTAGACTAAGGAACTCAAGACGCGGTTGAGGCC |
| Plppr2-1-SPL | /5Phos/CGTGAACTCCAGGCGGAAAAAACTCACTGATCAGGTCATACACTA |
| Plppr2-1-SPR | /5Phos/AAGATAAACATCGTAGACTACTAGACTCAGCACAGGGAAGGTGTC |
| Plppr2-2-SPL | /5Phos/CACCACTTGTCCCGCAAAAAAACTCACTGATCAGGTCATACACTA |
| Plppr2-2-SPR | /5Phos/AAGATAAACATCGTAGACTACTAGACTCAGTGTGGGGTTACCGGT |
| Plppr2-3-SPL | /5Phos/CTGAGCCACACTTAACAAAAAACTCACTGATCAGGTC |

| | |
|---|---|
| | ATACACTA |
| Plppr2-3-SPR | /5Phos/AAGATAAACATCGTAGACTACTAGACTCATGCAGGTC TCGGGTTC |
| Plppr3-1-SPL | /5Phos/AGTCAGCTCCAGGAAGAAAAAACTCAAGCTTCAGGTC ATACACTA |
| Plppr3-1-SPR | /5Phos/AAGATAAACATCGTAGACTACTGAACTCACCGGCTTG AACAGGTC |
| Plppr3-2-SPL | /5Phos/TCACAGGAAGTGCCCAAAAAAACTCAAGCTTCAGGTC ATACACTA |
| Plppr3-2-SPR | /5Phos/AAGATAAACATCGTAGACTACTGAACTCAGATGTAAG GGTTCGAC |
| Plppr3-3-SPL | /5Phos/CACATCCACGCTGGCTAAAAAACTCAAGCTTCAGGTC ATACACTA |
| Plppr3-3-SPR | /5Phos/AAGATAAACATCGTAGACTACTGAACTCAGTGGGGCC AGCAGGTC |
| Plppr4-1-SPL | /5Phos/TGTTAGGCTCCAATCCAAAAAACTCACTTCTCAGGTC ATACACTA |
| Plppr4-1-SPR | /5Phos/AAGATAAACATCGTAGACTATCCTACTCACAGCCTCC GGCGTTGA |
| Plppr4-2-SPL | /5Phos/CTCCTCCTATTAAAAAAAAAAACTCACTTCTCAGGTCA TACACTA |
| Plppr4-2-SPR | /5Phos/AAGATAAACATCGTAGACTATCCTACTCAAAATATAGT GCGATTC |
| Plppr4-3-SPL | /5Phos/TCGGGATGAGCCTCCAAAAAAACTCACTTCTCAGGTC ATACACTA |
| Plppr4-3-SPR | /5Phos/AAGATAAACATCGTAGACTATCCTACTCAGATGGGCC TGTTGTTC |
| Plppr5-1-SPL | /5Phos/TGCACGTTCACAGTGAAAAAAACTCAAGAGTCAGGTC ATACACTA |
| Plppr5-1-SPR | /5Phos/AAGATAAACATCGTAGACTAGAGAACTCAGTGGCAGA AGAATCCC |
| Plppr5-2-SPL | /5Phos/TTGGATGGGAATGTTTAAAAAACTCAAGAGTCAGGTC ATACACTA |
| Plppr5-2-SPR | /5Phos/AAGATAAACATCGTAGACTAGAGAACTCAACTCAGGG CAGCTTCC |
| Plppr5-3-SPL | /5Phos/ATACAACCAGAAATACAAAAAACTCAAGAGTCAGGTC ATACACTA |
| Plppr5-3-SPR | /5Phos/AAGATAAACATCGTAGACTAGAGAACTCAAAATTGTTT ACCACGC |
| Cxcl1-1-SPL | /5Phos/GAGCGGGTGGCTGGGAAAAAAACTCAAGGATCAGGT CATACACTA |
| Cxcl1-1-SPR | /5Phos/AAGATAAACATCGTAGACTACTCTACTCACGCTGCAC AGAGAAGC |
| Cxcl1-2-SPL | /5Phos/GATAGGCGCCCCTGTGAAAAAACTCAAGGATCAGGT |

| | |
|---|---|
| | CATACACTA |
| Cxcl1-2-SPR | /5Phos/AAGATAAACATCGTAGACTACTCTACTCAAGCGCAGCTCATTGGC |
| Cxcl1-3-SPL | /5Phos/TTGAGTGTGGCTATGAAAAAAACTCAAGGATCAGGTCATACACTA |
| Cxcl1-3-SPR | /5Phos/AAGATAAACATCGTAGACTACTCTACTCAAGCCTCGCGACCATTC |
| Il6-1-SPL | /5Phos/TGGATGGAAGTCTCTTAAAAAACTCAAGCTTCAGGTCATACACTA |
| Il6-1-SPR | /5Phos/AAGATAAACATCGTAGACTAGAGAACTCAGTCCCAAGAAGGCAAC |
| Il6-2-SPL | /5Phos/ATCGTTGTTCATACAAAAAAAACTCAAGCTTCAGGTCATACACTA |
| Il6-2-SPR | /5Phos/AAGATAAACATCGTAGACTAGAGAACTCATTTCTGCAAGTGCATC |
| Il6-3-SPL | /5Phos/CAGAAGACCAGAGGAAAAAAAACTCAAGCTTCAGGTCATACACTA |
| Il6-3-SPR | /5Phos/AAGATAAACATCGTAGACTAGAGAACTCAGGTAGCTATGGTACTC |
| Atm-1-SPL | /5Phos/TGTGCTTTTCTTATCTAAAAAACTCAAGAGTCAGGTCATACACTA |
| Atm-1-SPR | /5Phos/AAGATAAACATCGTAGACTAGACTACTCATTTTCTCTTGCTCGTC |
| Atm-2-SPL | /5Phos/TGCATGACAGCATCTTAAAAAACTCAAGAGTCAGGTCATACACTA |
| Atm-2-SPR | /5Phos/AAGATAAACATCGTAGACTAGACTACTCACATCTGGAAGACCTGC |
| Atm-3-SPL | /5Phos/AGAGAAGCCAACACTAAAAAAACTCAAGAGTCAGGTCATACACTA |
| Atm-3-SPR | /5Phos/AAGATAAACATCGTAGACTAGACTACTCATGAGATTTTGCTGGAC |
| Nbn-1-SPL | /5Phos/TTGGTCCTGGAGTCGTAAAAAACTCACAAGTCAGGTCATACACTA |
| Nbn-1-SPR | /5Phos/AAGATAAACATCGTAGACTACTTCACTCAAGGACTTGGGAAAGGC |
| Nbn-2-SPL | /5Phos/TGCTGCTGCTGAGAAGAAAAAACTCACAAGTCAGGTCATACACTA |
| Nbn-2-SPR | /5Phos/AAGATAAACATCGTAGACTACTTCACTCAGTTTTTGATGGAGTTC |
| Nbn-3-SPL | /5Phos/TGTCCTGAAGACCATCAAAAAACTCACAAGTCAGGTCATACACTA |
| Nbn-3-SPR | /5Phos/AAGATAAACATCGTAGACTACTTCACTCAGGCAGCTCCTCACTGC |
| Nfkb1-1-SPL | /5Phos/TGTCCGAGAAGTTCGGAAAAAACTCAAGCTTCAGGTC |

| | |
|---|---|
| | ATACACTA |
| Nfkb1-1-SPR | /5Phos/AAGATAAACATCGTAGACTAGACTACTCACTGCCGCCACCGAAGC |
| Nfkb1-2-SPL | /5Phos/TCTGTCATCCGTGCTTAAAAAACTCAAGCTTCAGGTCATACACTA |
| Nfkb1-2-SPR | /5Phos/AAGATAAACATCGTAGACTAGACTACTCAGCCCCTAATACACGCC |
| Nfkb1-3-SPL | /5Phos/TGAGTTTGCGGAAGGAAAAAAACTCAAGCTTCAGGTCATACACTA |
| Nfkb1-3-SPR | /5Phos/AAGATAAACATCGTAGACTAGACTACTCAAGAGACTCTGTAAAGC |
| Nfkb2-1-SPL | /5Phos/TCCACGATGGAGGGGCAAAAAACTCAAGCTTCAGGTCATACACTA |
| Nfkb2-1-SPR | /5Phos/AAGATAAACATCGTAGACTATCCTACTCAGGCTGGATCCTTAGGC |
| Nfkb2-2-SPL | /5Phos/TTGTCCATTCGGGAGAAAAAAACTCAAGCTTCAGGTCATACACTA |
| Nfkb2-2-SPR | /5Phos/AAGATAAACATCGTAGACTATCCTACTCACACGGAACCCGCTGTC |
| Nfkb2-3-SPL | /5Phos/ATGGATGATGGCCAGGAAAAAACTCAAGCTTCAGGTCATACACTA |
| Nfkb2-3-SPR | /5Phos/AAGATAAACATCGTAGACTATCCTACTCAATGACACCAGTCTGCCC |
| Cep128-1-SPL | /5Phos/TCTCGTTCTATTTCCTAAAAAACTCAAGTCTCAGGTCATACACTA |
| Cep128-1-SPR | /5Phos/AAGATAAACATCGTAGACTACTGAACTCATCTTTCTAAGCGCATC |
| Cep128-2-SPL | /5Phos/GAGGGAATTCTTGACCAAAAAACTCAAGTCTCAGGTCATACACTA |
| Cep128-2-SPR | /5Phos/AAGATAAACATCGTAGACTACTGAACTCAATCTATGGATGGTGTC |
| Cep128-3-SPL | /5Phos/TCACACAGCCACTGCAAAAAAACTCAAGTCTCAGGTCATACACTA |
| Cep128-3-SPR | /5Phos/AAGATAAACATCGTAGACTACTGAACTCATCTCTCTTTCAGTTCC |
| Irf1-1-SPL | /5Phos/AGGCATCCTTGTTGATAAAAAACTCAAGTCTCAGGTCATACACTA |
| Irf1-1-SPR | /5Phos/AAGATAAACATCGTAGACTATCCTACTCACAGCTCCGGAACAGAC |
| Irf1-2-SPL | /5Phos/TTGTTCCTACTCTGATAAAAAACTCAAGTCTCAGGTCATACACTA |
| Irf1-2-SPR | /5Phos/AAGATAAACATCGTAGACTATCCTACTCACACAGCAGAGCTGCCC |
| Irf1-3-SPL | /5Phos/CATCGATGTGTGTCGGAAAAAACTCAAGTCTCAGGTC |

| | |
|---|---|
| | ATACACTA |
| Irf1-3-SPR | /5Phos/AAGATAAACATCGTAGACTATCCTACTCAAGCAAGTATCCCTTGC |
| Mapk1-1-SPL | /5Phos/TCAAAAGGACTGATTTAAAAAACTCAAGGATCAGGTCATACACTA |
| Mapk1-1-SPR | /5Phos/AAGATAAACATCGTAGACTAGACTACTCAACAGTAGGTCTGGTGC |
| Mapk1-2-SPL | /5Phos/AGCAGGAGGTTGGAAGAAAAAAACTCAAGGATCAGGTCATACACTA |
| Mapk1-2-SPR | /5Phos/AAGATAAACATCGTAGACTAGACTACTCAATCACAAGTGGTGTTC |
| Mapk1-3-SPL | /5Phos/GGGAGAGAAAGCAAATAAAAAACTCAAGGATCAGGTCATACACTA |
| Mapk1-3-SPR | /5Phos/AAGATAAACATCGTAGACTAGACTACTCACACCTTATTTTTGTGC |
| Brca1-1-SPL | /5Phos/AGTGGTTTTGCCACTGAAAAAACTCACTAGTCAGGTCATACACTA |
| Brca1-1-SPR | /5Phos/AAGATAAACATCGTAGACTATCCTACTCATCTCCTCAGATCGGTC |
| Brca1-2-SPL | /5Phos/AGCGTTTGGACCTACCAAAAAACTCACTAGTCAGGTCATACACTA |
| Brca1-2-SPR | /5Phos/AAGATAAACATCGTAGACTATCCTACTCATATCACTAAGGGAGTC |
| Brca1-3-SPL | /5Phos/TTGGACCTTGGTGATTAAAAAACTCACTAGTCAGGTCATACACTA |
| Brca1-3-SPR | /5Phos/AAGATAAACATCGTAGACTATCCTACTCAGATTCTCTGGATCGCC |
| Sod1-1-SPL | /5Phos/TCGAAGTGGATGGTTCAAAAAACTCACTGATCAGGTCATACACTA |
| Sod1-1-SPR | /5Phos/AAGATAAACATCGTAGACTAGATCACTCAACCGCTTGCCTTCTGC |
| Sod1-2-SPL | /5Phos/TGCACTGGTACAGCCTAAAAAACTCACTGATCAGGTCATACACTA |
| Sod1-2-SPR | /5Phos/AAGATAAACATCGTAGACTAGATCACTCAGATTAAAATGAGGTCC |
| Sod1-3-SPL | /5Phos/CCAGGTCTCCAACATGAAAAAACTCACTGATCAGGTCATACACTA |
| Sod1-3-SPR | /5Phos/AAGATAAACATCGTAGACTAGATCACTCACCAGCAGTCACATTGC |
| Sod2-1-SPL | /5Phos/AATCCCCAGCAGCGGAAAAAAACTCACTGATCAGGTCATACACTA |
| Sod2-1-SPR | /5Phos/AAGATAAACATCGTAGACTATCGAACTCACGTGCTCCCACACGTC |
| Sod2-2-SPL | /5Phos/AGGTCTGGGAGGCTGTAAAAAACTCACTGATCAGGT |

| | |
|---|---|
| | CATACACTA |
| Sod2-2-SPR | /5Phos/AAGATAAACATCGTAGACTATCGAACTCAGCCATAGTCGTAAGGC |
| Sod2-3-SPL | /5Phos/GACCTGAGTTGTAACAAAAAAACTCACTGATCAGGTCATACACTA |
| Sod2-3-SPR | /5Phos/AAGATAAACATCGTAGACTATCGAACTCAGTGCAGGCTGAAGAGC |
| Cat-1-SPL | /5Phos/TGCTTCATCTGGTCGCAAAAAACTCACTTCTCAGGTCATACACTA |
| Cat-1-SPR | /5Phos/AAGATAAACATCGTAGACTACTGAACTCACCGCTGCTCCTTCCAC |
| Cat-2-SPL | /5Phos/TGAAGCATTTTGTCAGAAAAAACTCACTTCTCAGGTCATACACTA |
| Cat-2-SPR | /5Phos/AAGATAAACATCGTAGACTACTGAACTCAGGCAAAAAGGCGGCCC |
| Cat-3-SPL | /5Phos/TGAACAAGAAAGAAACAAAAAACTCACTTCTCAGGTCATACACTA |
| Cat-3-SPR | /5Phos/AAGATAAACATCGTAGACTACTGAACTCAGGAATCCCTCGGTCAC |
| Clock-1-SPL | /5Phos/AGAAGGAGTTGGGCTGAAAAAACTCACTAGTCAGGTCATACACTA |
| Clock-1-SPR | /5Phos/AAGATAAACATCGTAGACTAGAAGACTCAAGCTTCTAGAGGAGGC |
| Clock-2-SPL | /5Phos/TGCCGATGAATATTTGAAAAAACTCACTAGTCAGGTCATACACTA |
| Clock-2-SPR | /5Phos/AAGATAAACATCGTAGACTAGAAGACTCACCTTAGTTCTTCTTGC |
| Clock-3-SPL | /5Phos/TGGGAAATCACCATCGAAAAAACTCACTAGTCAGGTCATACACTA |
| Clock-3-SPR | /5Phos/AAGATAAACATCGTAGACTAGAAGACTCAGCTCCCAGCTGCAGGC |
| Clock-4-SPL | /5Phos/TGTTGCTGTGGTGGCTAAAAAACTCACTAGTCAGGTCATACACTA |
| Clock-4-SPR | /5Phos/AAGATAAACATCGTAGACTAGAAGACTCACTGCTGTTGTTGTTGC |
| Bmal1-1-SPL | /5Phos/ACGGTGAGTTTATCTAAAAAAACTCATCCTTCAGGTCATACACTA |
| Bmal1-1-SPR | /5Phos/AAGATAAACATCGTAGACTAAGTCACTCAAACAGCCATCCTTAGC |
| Bmal1-2-SPL | /5Phos/AGCGGCGGGCTCCAGAAAAAAACTCATCCTTCAGGTCATACACTA |
| Bmal1-2-SPR | /5Phos/AAGATAAACATCGTAGACTAAGTCACTCAATTCTACAGAAGAAAG |
| Bmal1-3-SPL | /5Phos/ATGTGCGAGTGCAGGCAAAAAACTCATCCTTCAGGTC |

| | |
|---|---|
| | ATACACTA |
| Bmal1-3-SPR | /5Phos/AAGATAAACATCGTAGACTAAGTCACTCACGCTGGTT GTGGAACC |
| Bmal1-4-SPL | /5Phos/AGCGGGCTGGAGCCAGAAAAAACTCATCCTTCAGGT CATACACTA |
| Bmal1-4-SPR | /5Phos/AAGATAAACATCGTAGACTAAGTCACTCACGTACTTG TGATGTTC |
| Cry1-1-SPL | /5Phos/GCAGCGGATGGTGTCGAAAAAACTCATCCTTCAGGT CATACACTA |
| Cry1-1-SPR | /5Phos/AAGATAAACATCGTAGACTAAGCTACTCAGGTCGAGG ATATAGAC |
| Cry1-2-SPL | /5Phos/AGAGGACAGGCCATCTAAAAAACTCATCCTTCAGGTC ATACACTA |
| Cry1-2-SPR | /5Phos/AAGATAAACATCGTAGACTAAGCTACTCACTCCTGGC CACACTGC |
| Cry1-3-SPL | /5Phos/TGCTGTATAAAAAAATAAAAAACTCATCCTTCAGGTCA TACACTA |
| Cry1-3-SPR | /5Phos/AAGATAAACATCGTAGACTAAGCTACTCAGTGGGTTG TTTGTGGC |
| Cry1-4-SPL | /5Phos/TCGGGACATTCTCTCCAAAAAACTCATCCTTCAGGTC ATACACTA |
| Cry1-4-SPR | /5Phos/AAGATAAACATCGTAGACTAAGCTACTCACCGCTGCT GCTACAAC |
| Cry2-1-SPL | /5Phos/AGCTTTCTTAAGCTTGAAAAAACTCATCCTTCAGGTCA TACACTA |
| Cry2-1-SPR | /5Phos/AAGATAAACATCGTAGACTAAGCTACTCAAAACAGAC GCGAATTC |
| Cry2-2-SPL | /5Phos/AGTTCGATGATTCTGTAAAAAACTCATCCTTCAGGTCA TACACTA |
| Cry2-2-SPR | /5Phos/AAGATAAACATCGTAGACTAAGCTACTCATGGTTTCT GCCCATTC |
| Cry2-3-SPL | /5Phos/CAAGGGAAGCCTGTCTAAAAAACTCATCCTTCAGGTC ATACACTA |
| Cry2-3-SPR | /5Phos/AAGATAAACATCGTAGACTAAGCTACTCACATGATGG CGTCAATC |
| Cry2-4-SPL | /5Phos/GACAGCTGTTGGTAGAAAAAAACTCATCCTTCAGGTC ATACACTA |
| Cry2-4-SPR | /5Phos/AAGATAAACATCGTAGACTAAGCTACTCAGAGTCCCC GGTATCTC |
| Dbp-1-SPL | /5Phos/CTGCCGCAACAGGGCGAAAAAACTCATCCTTCAGGT CATACACTA |
| Dbp-1-SPR | /5Phos/AAGATAAACATCGTAGACTACTTCACTCAGCACAGCC ACCACCTC |
| Dbp-2-SPL | /5Phos/TTGCTGTTTCCCTGCAAAAAAACTCATCCTTCAGGTC |

| | |
|---|---|
| | ATACACTA |
| Dbp-2-SPR | /5Phos/AAGATAAACATCGTAGACTACTTCACTCAGGCCGGTT CTTTGGGC |
| Dbp-3-SPL | /5Phos/AGGGCGAGATCAGCGGAAAAAACTCATCCTTCAGGT CATACACTA |
| Dbp-3-SPR | /5Phos/AAGATAAACATCGTAGACTACTTCACTCAGCCTGGAA TGCTTGAC |
| Dbp-4-SPL | /5Phos/TTCATTGTTCTTGTACAAAAAACTCATCCTTCAGGTCA TACACTA |
| Dbp-4-SPR | /5Phos/AAGATAAACATCGTAGACTACTTCACTCATCGACCTC TTGGCTGC |
| Gria4-1-SPL | /5Phos/TGGTGTTATGAAGAAAAAAAAACTCATCTCTCAGGTC ATACACTA |
| Gria4-1-SPR | /5Phos/AAGATAAACATCGTAGACTACTTCACTCATCAGATGC ATTGGGGC |
| Gria4-2-SPL | /5Phos/ATTTCCTCTCCTTGAAAAAAAACTCATCTCTCAGGTCA TACACTA |
| Gria4-2-SPR | /5Phos/AAGATAAACATCGTAGACTACTTCACTCACCAGACAA TCCCCGGC |
| Gria4-3-SPL | /5Phos/AGGGAAAACCAGAGGCAAAAAACTCATCTCTCAGGTC ATACACTA |
| Gria4-3-SPR | /5Phos/AAGATAAACATCGTAGACTACTTCACTCATTGCATAAA GGCACCC |
| Gria4-4-SPL | /5Phos/AGCATTTCTTAATGAGAAAAAACTCATCTCTCAGGTCA TACACTA |
| Gria4-4-SPR | /5Phos/AAGATAAACATCGTAGACTACTTCACTCAAAACTGCG AGGTTAAC |
| F2rl1-1-SPL | /5Phos/TGGGTTTCTAATCTGCAAAAAACTCAGATCTCAGGTC ATACACTA |
| F2rl1-1-SPR | /5Phos/AAGATAAACATCGTAGACTACTAGACTCACCCAGTGA TTGGAGGC |
| F2rl1-2-SPL | /5Phos/ACGGCGGGGTGTTTCTAAAAAACTCAGATCTCAGGTC ATACACTA |
| F2rl1-2-SPR | /5Phos/AAGATAAACATCGTAGACTACTAGACTCAGTTGGCCA TGTAAATC |
| F2rl1-3-SPL | /5Phos/ACCCAGTACCTCTGCAAAAAAACTCAGATCTCAGGTC ATACACTA |
| F2rl1-3-SPR | /5Phos/AAGATAAACATCGTAGACTACTAGACTCACATGGGGT TCACGATC |
| F2rl1-4-SPL | /5Phos/TGTTGAGGGTCGACAGAAAAAACTCAGATCTCAGGTC ATACACTA |
| F2rl1-4-SPR | /5Phos/AAGATAAACATCGTAGACTACTAGACTCAAAGGGGTC TATGCAGC |
| C5ar1-1-SPL | /5Phos/AGGTATGTTAGGATCCAAAAAACTCAGAGATCAGGTC |

| | |
|---|---|
| | ATACACTA |
| C5ar1-1-SPR | /5Phos/AAGATAAACATCGTAGACTACTCTACTCAGGTGAATGCCATCCGC |
| C5ar1-2-SPL | /5Phos/GAGAGGAGGTCGGCCAAAAAAACTCAGAGATCAGGTCATACACTA |
| C5ar1-2-SPR | /5Phos/AAGATAAACATCGTAGACTACTCTACTCAAGGCAGTGCCAAGCAC |
| C5ar1-3-SPL | /5Phos/TTTGAGCGTCTTGGTGAAAAAACTCAGAGATCAGGTCATACACTA |
| C5ar1-3-SPR | /5Phos/AAGATAAACATCGTAGACTACTCTACTCACCACAGCCATCACCAC |
| C5ar1-4-SPL | /5Phos/AGGGACACGCACAGGGAAAAAACTCAGAGATCAGGTCATACACTA |
| C5ar1-4-SPR | /5Phos/AAGATAAACATCGTAGACTACTCTACTCAGCAGTTGATGTAGGCC |
| Tnf-1-SPL | /5Phos/TGCCAGTTCCACGTCGAAAAAACTCAAGGATCAGGTCATACACTA |
| Tnf-1-SPR | /5Phos/AAGATAAACATCGTAGACTACTAGACTCAGGGGGAGTGCCTCTTC |
| Tnf-2-SPL | /5Phos/CATTTGGGAACTTCTCAAAAAACTCAAGGATCAGGTCATACACTA |
| Tnf-2-SPR | /5Phos/AAGATAAACATCGTAGACTACTAGACTCACTGATGAGAGGGAGGC |
| Tnf-3-SPL | /5Phos/TAGTTGGTTGTCTTTGAAAAAACTCAAGGATCAGGTCATACACTA |
| Tnf-3-SPR | /5Phos/AAGATAAACATCGTAGACTACTAGACTCACATCGGCTGGCACCAC |
| C5ar2-1-SPL | /5Phos/AGCCAGCAGCCTGCCGAAAAAACTCAGAGATCAGGTCATACACTA |
| C5ar2-1-SPR | /5Phos/AAGATAAACATCGTAGACTACTAGACTCAGGTGATGGAGGACAAC |
| C5ar2-2-SPL | /5Phos/ACAAGTCGGGGAGGATAAAAAACTCAGAGATCAGGTCATACACTA |
| C5ar2-2-SPR | /5Phos/AAGATAAACATCGTAGACTACTAGACTCAATAGTCTATGCCACAC |
| C5ar2-3-SPL | /5Phos/CAGCAAATGAAAAACCAAAAAACTCAGAGATCAGGTCATACACTA |
| C5ar2-3-SPR | /5Phos/AAGATAAACATCGTAGACTACTAGACTCAGACGTGATAAGGGGTC |
| C5ar2-4-SPL | /5Phos/ACTGTGAGCCAGGGCCAAAAAACTCAGAGATCAGGTCATACACTA |
| C5ar2-4-SPR | /5Phos/AAGATAAACATCGTAGACTACTAGACTCATTATGGGATTGAGAGC |
| Csf3r-1-SPL | /5Phos/AGAAGAGGAAGGCCTGAAAAAACTCAGAAGTCAGGT |

| | |
|---|---|
| | CATACACTA |
| Csf3r-1-SPR | /5Phos/AAGATAAACATCGTAGACTATCGAACTCATCCCATGGCACTAAGC |
| Csf3r-2-SPL | /5Phos/CTTGATGATCTGGGGAAAAAAACTCAGAAGTCAGGTCATACACTA |
| Csf3r-2-SPR | /5Phos/AAGATAAACATCGTAGACTATCGAACTCAAGGTGTATGTCCTGCC |
| Csf3r-3-SPL | /5Phos/TTCAGGACAAAGTCATAAAAAACTCAGAAGTCAGGTCATACACTA |
| Csf3r-3-SPR | /5Phos/AAGATAAACATCGTAGACTATCGAACTCAGGCGGGCTCCAGGTGC |
| Csf3r-4-SPL | /5Phos/ACCTGACCATAGAGGAAAAAAACTCAGAAGTCAGGTCATACACTA |
| Csf3r-4-SPR | /5Phos/AAGATAAACATCGTAGACTATCGAACTCAGGTGGGGCTCTCAAGC |
| Ccr7-1-SPL | /5Phos/CAGCCCGTTGCCGAGCAAAAAACTCAGAAGTCAGGTCATACACTA |
| Ccr7-1-SPR | /5Phos/AAGATAAACATCGTAGACTATCTCACTCATGTACGTCAGTATCAC |
| Ccr7-2-SPL | /5Phos/GCCAAAGATCCAGGACAAAAAACTCAGAAGTCAGGTCATACACTA |
| Ccr7-2-SPR | /5Phos/AAGATAAACATCGTAGACTATCTCACTCACCTTACACAGGTAGAC |
| Ccr7-3-SPL | /5Phos/AGCGTGTCCTCGCCGCAAAAAACTCAGAAGTCAGGTCATACACTA |
| Ccr7-3-SPR | /5Phos/AAGATAAACATCGTAGACTATCTCACTCAGACCAGTGAGCATCTC |
| Ccr7-4-SPL | /5Phos/AATGATCACCTTGATGAAAAAACTCAGAAGTCAGGTCATACACTA |
| Ccr7-4-SPR | /5Phos/AAGATAAACATCGTAGACTATCTCACTCAAGACTACCACCACGGC |
| Xcl1-1-SPL | /5Phos/TCCCAGGAAAGTCAGGAAAAAACTCAGAAGTCAGGTCATACACTA |
| Xcl1-1-SPR | /5Phos/AAGATAAACATCGTAGACTACTGAACTCAGGGTGAGGCAGCAGAC |
| Xcl1-2-SPL | /5Phos/TGGGTTTGTAAGTTCAAAAAAACTCAGAAGTCAGGTCATACACTA |
| Xcl1-2-SPR | /5Phos/AAGATAAACATCGTAGACTACTGAACTCATTGAACTGGCAGCCGC |
| Xcl1-3-SPL | /5Phos/CTCGTTTGGTGACAAAAAAAAACTCAGAAGTCAGGTCATACACTA |
| Xcl1-3-SPR | /5Phos/AAGATAAACATCGTAGACTACTGAACTCAGCACAAATTTTTAGTC |
| Xcl1-4-SPL | /5Phos/TCTGGGCTCCTGTGGGAAAAAACTCAGAAGTCAGGT |

| | |
|---|---|
| | CATACACTA |
| Xcl1-4-SPR | /5Phos/AAGATAAACATCGTAGACTACTGAACTCAGCTGTGCTGGTGGACC |
| Cd74-1-SPL | /5Phos/AGAGACACCGGTGTACAAAAAACTCAGAAGTCAGGTCATACACTA |
| Cd74-1-SPR | /5Phos/AAGATAAACATCGTAGACTAGAAGACTCAGCAGAGCCACCAGGAC |
| Cd74-2-SPL | /5Phos/CTCGTGAGCAGATGCAAAAAAACTCAGAAGTCAGGTCATACACTA |
| Cd74-2-SPR | /5Phos/AAGATAAACATCGTAGACTAGAAGACTCACTCCAGGGGTCCAGAC |
| Cd74-3-SPL | /5Phos/TTGCTCATCTCAAACAAAAAAACTCAGAAGTCAGGTCATACACTA |
| Cd74-3-SPR | /5Phos/AAGATAAACATCGTAGACTAGAAGACTCACTCCTCCAGGGAGTTC |
| Cd74-4-SPL | /5Phos/TCGTCGCACTTGGGACAAAAAACTCAGAAGTCAGGTCATACACTA |
| Cd74-4-SPR | /5Phos/AAGATAAACATCGTAGACTAGAAGACTCACAAATAGTTACCGTTC |
| Pik3cb-1-SPL | /5Phos/TTTCCTCCATAAAGTTAAAAAACTCACTCTTCAGGTCATACACTA |
| Pik3cb-1-SPR | /5Phos/AAGATAAACATCGTAGACTACTAGACTCACACAGCCACAACCAGC |
| Pik3cb-2-SPL | /5Phos/TCAGCTGTCCTTTGAAAAAAAACTCACTCTTCAGGTCATACACTA |
| Pik3cb-2-SPR | /5Phos/AAGATAAACATCGTAGACTACTAGACTCAATGACGTCTCCAGACC |
| Pik3cb-3-SPL | /5Phos/AGGAGTGTGCACCTCTAAAAAACTCACTCTTCAGGTCATACACTA |
| Pik3cb-3-SPR | /5Phos/AAGATAAACATCGTAGACTACTAGACTCAACTGTACGGACACAGC |
| Pik3cb-4-SPL | /5Phos/AGGAGTGCGTCTTTGTAAAAAACTCACTCTTCAGGTCATACACTA |
| Pik3cb-4-SPR | /5Phos/AAGATAAACATCGTAGACTACTAGACTCACTCCTTGAGCCAGTTC |
| Pik3cd-1-SPL | /5Phos/TGAGCATGTGGAAGAGAAAAAACTCACTCTTCAGGTCATACACTA |
| Pik3cd-1-SPR | /5Phos/AAGATAAACATCGTAGACTAGATCACTCATAGGCCTCGGGGTCAC |
| Pik3cd-2-SPL | /5Phos/TTCTTGGCAGGGATCGAAAAAACTCACTCTTCAGGTCATACACTA |
| Pik3cd-2-SPR | /5Phos/AAGATAAACATCGTAGACTAGATCACTCAGGACACAGAGGAGGGC |
| Pik3cd-3-SPL | /5Phos/CCACGTTGCCAGCACTAAAAAACTCACTCTTCAGGTC |

| | |
|---|---|
| | ATACACTA |
| Pik3cd-3-SPR | /5Phos/AAGATAAACATCGTAGACTAGATCACTCATTCTTAAAG ATGATGC |
| Pik3cd-4-SPL | /5Phos/AAAGTCGTAGGTGAGAAAAAAACTCACTCTTCAGGTC ATACACTA |
| Pik3cd-4-SPR | /5Phos/AAGATAAACATCGTAGACTAGATCACTCAGCTGGATC ACGTGGAC |
| Prkca-1-SPL | /5Phos/TCATGCACGTTCTTCTAAAAAACTCACTTCTCAGGTCA TACACTA |
| Prkca-1-SPR | /5Phos/AAGATAAACATCGTAGACTACTAGACTCATTTGTGGT CTTTCACC |
| Prkca-2-SPL | /5Phos/AGGTGTCACATTTCATAAAAAACTCACTTCTCAGGTCA TACACTA |
| Prkca-2-SPR | /5Phos/AAGATAAACATCGTAGACTACTAGACTCATGAACATT CATGTCGC |
| Prkca-3-SPL | /5Phos/TTCCCAAAACTCCCCTAAAAAACTCACTTCTCAGGTC ATACACTA |
| Prkca-3-SPR | /5Phos/AAGATAAACATCGTAGACTACTAGACTCAGTCAGCAA GCATCACC |
| Prkca-4-SPL | /5Phos/TCTCTGACATCCCTCTAAAAAACTCACTTCTCAGGTCA TACACTA |
| Prkca-4-SPR | /5Phos/AAGATAAACATCGTAGACTACTAGACTCACCTGAAGA AGGCATGC |
| Vav1-1-SPL | /5Phos/TGTTGGGAAGGGCATGAAAAAACTCACTTCTCAGGTC ATACACTA |
| Vav1-1-SPR | /5Phos/AAGATAAACATCGTAGACTATCGAACTCATCAGAGCG CTGTCCTC |
| Vav1-2-SPL | /5Phos/ATAGGTACCATCAGCAAAAAAACTCACTTCTCAGGTC ATACACTA |
| Vav1-2-SPR | /5Phos/AAGATAAACATCGTAGACTATCGAACTCACTTCAGCA CCCGCTGC |
| Vav1-3-SPL | /5Phos/CTGCGAAATCTTGCCCAAAAAACTCACTTCTCAGGTC ATACACTA |
| Vav1-3-SPR | /5Phos/AAGATAAACATCGTAGACTATCGAACTCATCCTTCTTC ATGGTTC |
| Vav1-4-SPL | /5Phos/AAGGTGGTGTCCAACGAAAAAACTCACTTCTCAGGTC ATACACTA |
| Vav1-4-SPR | /5Phos/AAGATAAACATCGTAGACTATCGAACTCACTTATAAG GAAACTGC |
| Rac1-1-SPL | /5Phos/AGGTTTTACCAACAGCAAAAAACTCACTTCTCAGGTC ATACACTA |
| Rac1-1-SPR | /5Phos/AAGATAAACATCGTAGACTAAGCTACTCATAACTGAT GAGCAGGC |
| Rac1-2-SPL | /5Phos/AGATTCACTGGTTTTCAAAAAACTCACTTCTCAGGTCA |

| | |
|---|---|
| | TACACTA |
| Rac1-2-SPR | /5Phos/AAGATAAACATCGTAGACTAAGCTACTCATGTGTCCCATAGGCCC |
| Rac1-3-SPL | /5Phos/ACAAGGGAAAAGCAAAAAAAAACTCACTTCTCAGGTCATACACTA |
| Rac1-3-SPR | /5Phos/AAGATAAACATCGTAGACTAAGCTACTCAAAATGATGCAGGACTC |
| Rac1-4-SPL | /5Phos/AGCTTCTTCTCCTTCAAAAAAACTCACTTCTCAGGTCATACACTA |
| Rac1-4-SPR | /5Phos/AAGATAAACATCGTAGACTAAGCTACTCAGTAGGTGATGGGAGTC |
| Rac2-1-SPL | /5Phos/TGTCCACCATCACATTAAAAAACTCACTGATCAGGTCATACACTA |
| Rac2-1-SPR | /5Phos/AAGATAAACATCGTAGACTAGACTACTCAAGGTTCACCGGCTTAC |
| Rac2-2-SPL | /5Phos/TGGGCTGACTAGCGAGAAAAAACTCACTGATCAGGTCATACACTA |
| Rac2-2-SPR | /5Phos/AAGATAAACATCGTAGACTAGACTACTCACATTCTCATAGGAGGC |
| Rac2-3-SPL | /5Phos/TCGATGGTGTCCTTGTAAAAAACTCACTGATCAGGTCATACACTA |
| Rac2-3-SPR | /5Phos/AAGATAAACATCGTAGACTAGACTACTCACTTCTCCTTCAGCTTC |
| Rac2-4-SPL | /5Phos/TTCAGGCCTCGCTGGGAAAAAACTCACTGATCAGGTCATACACTA |
| Rac2-4-SPR | /5Phos/AAGATAAACATCGTAGACTAGACTACTCACTCATCGAAGACGGTC |
| Fyn-1-SPL | /5Phos/AGTGTGAGAGGAGGAGAAAAAACTCACTGATCAGGTCATACACTA |
| Fyn-1-SPR | /5Phos/AAGATAAACATCGTAGACTAGAAGACTCATCGTGCGTAGGGTCCC |
| Fyn-2-SPL | /5Phos/TGGGCCCGCGTTGTGAAAAAAACTCACTGATCAGGTCATACACTA |
| Fyn-2-SPR | /5Phos/AAGATAAACATCGTAGACTAGAAGACTCACTGAAGTGTTTCAAAC |
| Fyn-3-SPL | /5Phos/ATGGTGCCTGGCTTAAAAAAAACTCACTGATCAGGTCATACACTA |
| Fyn-3-SPR | /5Phos/AAGATAAACATCGTAGACTAGAAGACTCAGAAGGACTCCGGAGAC |
| Fyn-4-SPL | /5Phos/ACCTGCTCCAGCACCTAAAAAACTCACTGATCAGGTCATACACTA |
| Fyn-4-SPR | /5Phos/AAGATAAACATCGTAGACTAGAAGACTCACCTATAGCCTCTCTCC |
| Mpl-1-SPL | /5Phos/AGCAAGAAGACATCTTAAAAAACTCACTGATCAGGTC |

| | |
|---|---|
| | ATACACTA |
| Mpl-1-SPR | /5Phos/AAGATAAACATCGTAGACTACTGAACTCACTCTGTGCCCAAGGCC |
| Mpl-2-SPL | /5Phos/TGAATGACGGAGGGGGAAAAAACTCACTGATCAGGTCATACACTA |
| Mpl-2-SPR | /5Phos/AAGATAAACATCGTAGACTACTGAACTCATTCTGTGGAGAGCAGC |
| Mpl-3-SPL | /5Phos/TCTTGAGCTGCCCAAGAAAAAACTCACTGATCAGGTCATACACTA |
| Mpl-3-SPR | /5Phos/AAGATAAACATCGTAGACTACTGAACTCAGAGCTGGTAGCAGGTC |
| Mpl-4-SPL | /5Phos/GCAGGAAATTGCCACTAAAAAACTCACTGATCAGGTCATACACTA |
| Mpl-4-SPR | /5Phos/AAGATAAACATCGTAGACTACTGAACTCACAGTCTCCTGTAGTGC |
| Itgb2-1-SPL | /5Phos/CTCAGCTTCTCAGGATAAAAAACTCACTGATCAGGTCATACACTA |
| Itgb2-1-SPR | /5Phos/AAGATAAACATCGTAGACTACTTCACTCAGTTGGGACATGGGTTC |
| Itgb2-2-SPL | /5Phos/TGTGGCAAACACCAGCAAAAAACTCACTGATCAGGTCATACACTA |
| Itgb2-2-SPR | /5Phos/AAGATAAACATCGTAGACTACTTCACTCAAGTGGAAGCCATCGTC |
| Itgb2-3-SPL | /5Phos/AAAGGACTGCTCCTGGAAAAAACTCACTGATCAGGTCATACACTA |
| Itgb2-3-SPR | /5Phos/AAGATAAACATCGTAGACTACTTCACTCACCAGTGCCCGGATGAC |
| Itgb2-4-SPL | /5Phos/ATTCAGACAGCCCGTGAAAAAACTCACTGATCAGGTCATACACTA |
| Itgb2-4-SPR | /5Phos/AAGATAAACATCGTAGACTACTTCACTCAACTCTACCAGCCGTGC |
| P2ry12-1-SPL | /5Phos/AAAGAACAGGACGGTGAAAAAACTCACTGATCAGGTCATACACTA |
| P2ry12-1-SPR | /5Phos/AAGATAAACATCGTAGACTAAGCTACTCATCGTGATGAGCCCAGC |
| P2ry12-2-SPL | /5Phos/CCAGATGACAACAGAAAAAAAACTCACTGATCAGGTCATACACTA |
| P2ry12-2-SPR | /5Phos/AAGATAAACATCGTAGACTAAGCTACTCATTAAGAACATGAAGGC |
| P2ry12-3-SPL | /5Phos/AAAGACGGCCCGAGTTAAAAAACTCACTGATCAGGTCATACACTA |
| P2ry12-3-SPR | /5Phos/AAGATAAACATCGTAGACTAAGCTACTCATCTCAGCACTGCAGTC |
| Ptafr-1-SPL | /5Phos/TGCAGTAGGTATTGATAAAAAACTCACTAGTCAGGTC |

| | |
|---|---|
| | ATACACTA |
| Ptafr-1-SPR | /5Phos/AAGATAAACATCGTAGACTAAGCTACTCACCCAAAAA<br>GGCCACAC |
| Ptafr-2-SPL | /5Phos/GATGAAGACATGAACAAAAAAACTCACTAGTCAGGTC<br>ATACACTA |
| Ptafr-2-SPR | /5Phos/AAGATAAACATCGTAGACTAAGCTACTCAGGAAGAAG<br>CAGAAGGC |
| Ptafr-3-SPL | /5Phos/CATCCACAGCGCCCTCAAAAAACTCACTAGTCAGGTC<br>ATACACTA |
| Ptafr-3-SPR | /5Phos/AAGATAAACATCGTAGACTAAGCTACTCACCAAGACC<br>GTGCAGAC |
| Ptafr-4-SPL | /5Phos/TGAGGAGGCAGAGGGTAAAAAACTCACTAGTCAGGT<br>CATACACTA |
| Ptafr-4-SPR | /5Phos/AAGATAAACATCGTAGACTAAGCTACTCAAAGACACA<br>GTTGGTGC |
| Ppp1r3a-1-SPL | /5Phos/CAGGCTGTCAGCAAAGAAAAAACTCACTAGTCAGGTC<br>ATACACTA |
| Ppp1r3a-1-SPR | /5Phos/AAGATAAACATCGTAGACTAAGAGACTCAACACAAGG<br>CTGAATCC |
| Ppp1r3a-2-SPL | /5Phos/AGTGCTTATTTTTGGGAAAAAACTCACTAGTCAGGTC<br>ATACACTA |
| Ppp1r3a-2-SPR | /5Phos/AAGATAAACATCGTAGACTAAGAGACTCATAGCTTTAT<br>CCCCTTC |
| Ppp1r3a-3-SPL | /5Phos/TCTCGGCAGGTAAACAAAAAAACTCACTAGTCAGGTC<br>ATACACTA |
| Ppp1r3a-3-SPR | /5Phos/AAGATAAACATCGTAGACTAAGAGACTCATCCGTAGC<br>ATTCTGTC |
| Ppp1r3a-4-SPL | /5Phos/TCCTCCACTACCTGATAAAAAACTCACTAGTCAGGTC<br>ATACACTA |
| Ppp1r3a-4-SPR | /5Phos/AAGATAAACATCGTAGACTAAGAGACTCAGACCCCCG<br>CCTCGTTC |
| Tyro3-1-SPL | /5Phos/TTCACAAGAAAACTCTAAAAAACTCACTAGTCAGGTC<br>ATACACTA |
| Tyro3-1-SPR | /5Phos/AAGATAAACATCGTAGACTACTGAACTCACTTTTATGT<br>TGCGGGC |
| Tyro3-2-SPL | /5Phos/CCTTTGTCTGAAAGGGAAAAAACTCACTAGTCAGGTC<br>ATACACTA |
| Tyro3-2-SPR | /5Phos/AAGATAAACATCGTAGACTACTGAACTCACTGGCTGG<br>CGCTAGGC |
| Tyro3-3-SPL | /5Phos/AACAAGCTTGGCCACGAAAAAACTCACTAGTCAGGTC<br>ATACACTA |
| Tyro3-3-SPR | /5Phos/AAGATAAACATCGTAGACTACTGAACTCATCCGGAGG<br>CTCACCCC |
| Pcp4-1-SPL | /5Phos/TTTGTCTCTCACTCATAAAAAACTCAGAAGTCAGGTCA |

| | |
|---|---|
| | TACACTA |
| Pcp4-1-SPR | /5Phos/AAGATAAACATCGTAGACTAAGCTACTCATTGGTCGC TCCGGCAC |
| Pcp4-2-SPL | /5Phos/CATGTCGATATCAAATAAAAAACTCAGAAGTCAGGTC ATACACTA |
| Pcp4-2-SPR | /5Phos/AAGATAAACATCGTAGACTAAGCTACTCACTGTCTCT GGTGCATC |
| Pcp4-3-SPL | /5Phos/TTCTGGAATTTTCTGAAAAAAACTCAGAAGTCAGGTC ATACACTA |
| Pcp4-3-SPR | /5Phos/AAGATAAACATCGTAGACTAAGCTACTCATGATCCTG CCTTTTTC |
| Grik4-1-SPL | /5Phos/TGATGTCGGTGTTGCTAAAAAACTCAGAGATCAGGTC ATACACTA |
| Grik4-1-SPR | /5Phos/AAGATAAACATCGTAGACTAGACTACTCACCAGCCAC AGCCACGC |
| Grik4-2-SPL | /5Phos/AATAGTGGAGCCTTGCAAAAAACTCAGAGATCAGGTC ATACACTA |
| Grik4-2-SPR | /5Phos/AAGATAAACATCGTAGACTAGACTACTCAACAAAGCT CGAGGGGC |
| Grik4-3-SPL | /5Phos/CAGTTCTGTCACCATCAAAAAACTCAGAGATCAGGTC ATACACTA |
| Grik4-3-SPR | /5Phos/AAGATAAACATCGTAGACTAGACTACTCAACAGGATG ATGTTGCG |
| Zbtb20-1-SPL | /5Phos/AGGCAGGGAAGGGCCGAAAAAACTCAGAGATCAGGT CATACACTA |
| Zbtb20-1-SPR | /5Phos/AAGATAAACATCGTAGACTAGAGAACTCAAACAGCTT CAAAGTTC |
| Zbtb20-2-SPL | /5Phos/CCTCTGCTTGGTCAGTAAAAAACTCAGAGATCAGGTC ATACACTA |
| Zbtb20-2-SPR | /5Phos/AAGATAAACATCGTAGACTAGAGAACTCAGGCTCGCT CTCAGTGC |
| Zbtb20-3-SPL | /5Phos/AAGGAGAAGGAGCGCCAAAAAACTCAGAGATCAGGT CATACACTA |
| Zbtb20-3-SPR | /5Phos/AAGATAAACATCGTAGACTAGAGAACTCAGATAAGGT AATCCTTC |
| Gpx1-1-SPL | /5Phos/ATACACGGTGGACTGTAAAAAACTCACTTCTCAGGTC ATACACTA |
| Gpx1-1-SPR | /5Phos/AAGATAAACATCGTAGACTAGACTACTCAGGCGCGC GGAGAAGGC |
| Gpx1-2-SPL | /5Phos/TGATTGCACGGGAAACAAAAAACTCACTTCTCAGGTC ATACACTA |
| Gpx1-2-SPR | /5Phos/AAGATAAACATCGTAGACTAGACTACTCACTCCTGGT GTCCGAAC |
| Gpx1-3-SPL | /5Phos/CGGAGACCAAATGATGAAAAAACTCACTTCTCAGGTC |

| | |
|---|---|
| | ATACACTA |
| Gpx1-3-SPR | /5Phos/AAGATAAACATCGTAGACTAGACTACTCATGTCGTTG CGGCACAC |
| Gclc-1-SPL | /5Phos/TTGAGCACGTCCTTGTAAAAAACTCACTCTTCAGGTC ATACACTA |
| Gclc-1-SPR | /5Phos/AAGATAAACATCGTAGACTATCAGACTCACACCTCGT CACCCCAC |
| Gclc-2-SPL | /5Phos/TCCTTCCTCTGGGTTGAAAAAACTCACTCTTCAGGTC ATACACTA |
| Gclc-2-SPR | /5Phos/AAGATAAACATCGTAGACTATCAGACTCAGGGACTTT GATGCGCC |
| Gclc-3-SPL | /5Phos/TCCCCAGCGACAATCAAAAAAACTCACTCTTCAGGTC ATACACTA |
| Gclc-3-SPR | /5Phos/AAGATAAACATCGTAGACTATCAGACTCACAGAGGCA GAAATCAC |
| Gclm-1-SPL | /5Phos/TGATTTGGGAACTCCAAAAAAACTCACTCTTCAGGTC ATACACTA |
| Gclm-1-SPR | /5Phos/AAGATAAACATCGTAGACTAAGCTACTCACTGACTAA ATCGGGGC |
| Gclm-2-SPL | /5Phos/TGCACTTCTAGTTGATAAAAAACTCACTCTTCAGGTCA TACACTA |
| Gclm-2-SPR | /5Phos/AAGATAAACATCGTAGACTAAGCTACTCAAGCATGCC ATGTCAAC |
| Gclm-3-SPL | /5Phos/TCAGAGAGCAGTTCTTAAAAAACTCACTCTTCAGGTC ATACACTA |
| Gclm-3-SPR | /5Phos/AAGATAAACATCGTAGACTAAGCTACTCATTCCTGGA AACTTGCC |
| Vegfa-1-SPL | /5Phos/TCCCAGCTCCGATCGGAAAAAACTCATCCTTCAGGTC ATACACTA |
| Vegfa-1-SPR | /5Phos/AAGATAAACATCGTAGACTATCCTACTCACCCGAGCT AGCACTTC |
| Vegfa-2-SPL | /5Phos/TTGGCGATTTAGCAGCAAAAAACTCATCCTTCAGGTC ATACACTA |
| Vegfa-2-SPR | /5Phos/AAGATAAACATCGTAGACTATCCTACTCACCCTAATCT TCCGGGC |
| Vegfa-3-SPL | /5Phos/TGTGCTGTAGGAAGCTAAAAAACTCATCCTTCAGGTC ATACACTA |
| Vegfa-3-SPR | /5Phos/AAGATAAACATCGTAGACTATCCTACTCACTGCATTCA CATCTGC |
| Nos3-1-SPL | /5Phos/TGCGTTTGGGGCTGAAAAAAAACTCATCCTTCAGGTC ATACACTA |
| Nos3-1-SPR | /5Phos/AAGATAAACATCGTAGACTACTGAACTCACTCTGGCG CTTCCAGC |
| Nos3-2-SPL | /5Phos/TGAACTGACAGAGTAGAAAAAACTCATCCTTCAGGTC |

| | |
|---|---|
| | ATACACTA |
| Nos3-2-SPR | /5Phos/AAGATAAACATCGTAGACTACTGAACTCAGGTGGGCG CTGGGTGC |
| Nos3-3-SPL | /5Phos/ACATCCTCAAGTATGTAAAAAACTCATCCTTCAGGTCA TACACTA |
| Nos3-3-SPR | /5Phos/AAGATAAACATCGTAGACTACTGAACTCAATCCATGC ACACAGCC |
| Mmp9-1-SPL | /5Phos/TGCCGGACTCAAAGACAAAAAACTCATCGATCAGGTC ATACACTA |
| Mmp9-1-SPR | /5Phos/AAGATAAACATCGTAGACTAGAAGACTCACATTGCAA GGATTGTC |
| Mmp9-2-SPL | /5Phos/TTCCCGAGACGACGCGAAAAAACTCATCGATCAGGT CATACACTA |
| Mmp9-2-SPR | /5Phos/AAGATAAACATCGTAGACTAGAAGACTCAGCTGAACA GCAGAGCC |
| Mmp9-3-SPL | /5Phos/TCCGTTGCCGTGCTCCAAAAAACTCATCGATCAGGTC ATACACTA |
| Mmp9-3-SPR | /5Phos/AAGATAAACATCGTAGACTAGAAGACTCACACAGGGT TTGCCTTC |
| Hif1a-1-SPL | /5Phos/TTGCTGGCTGATCTTGAAAAAACTCATCAGTCAGGTC ATACACTA |
| Hif1a-1-SPR | /5Phos/AAGATAAACATCGTAGACTATCAGACTCACTTCCATC AGAAGGAC |
| Hif1a-2-SPL | /5Phos/TGGTGAGGCTGTCCGAAAAAAACTCATCAGTCAGGTC ATACACTA |
| Hif1a-2-SPR | /5Phos/AAGATAAACATCGTAGACTATCAGACTCATCTTTCCTG CTCTGTC |
| Hif1a-3-SPL | /5Phos/TGAGCATTCTGCGAAGAAAAAACTCATCAGTCAGGTC ATACACTA |
| Hif1a-3-SPR | /5Phos/AAGATAAACATCGTAGACTATCAGACTCACATTTTTCG CTTCCTC |
| Casp1-1-SPL | /5Phos/CACGGCATGCCTGAATAAAAAACTCATCAGTCAGGTC ATACACTA |
| Casp1-1-SPR | /5Phos/AGATAAACATCGTAGACTAAGGAACTCAACTCCTTGT TTCTCTC |
| Casp1-2-SPL | /5Phos/TTTGCCCTCAGGATCTAAAAAACTCATCAGTCAGGTC ATACACTA |
| Casp1-2-SPR | /5Phos/AAGATAAACATCGTAGACTAAGGAACTCAGATAAATT GCTTCCTC |
| Casp1-3-SPL | /5Phos/CCAGTCAGTCCTGGAAAAAAAACTCATCAGTCAGGTC ATACACTA |
| Casp1-3-SPR | /5Phos/AAGATAAACATCGTAGACTAAGGAACTCAGCAAAACT TGAGGGTC |
| Casp3-1-SPL | /5Phos/TTTTGCTATGATCTTCAAAAAACTCATCTCTCAGGTCA |

| | |
|---|---|
| | TACACTA |
| Casp3-1-SPR | /5Phos/AAGATAAACATCGTAGACTAGAAGACTCACACACAAA GCTGCTCC |
| Casp3-2-SPL | /5Phos/TCCTCATCAGTCCCACAAAAAACTCATCTCTCAGGTC ATACACTA |
| Casp3-2-SPR | /5Phos/AAGATAAACATCGTAGACTAGAAGACTCACTTCTGGC AAGCCATC |
| Casp3-3-SPL | /5Phos/GAGTGAGAATGTGCATAAAAAACTCATCTCTCAGGTC ATACACTA |
| Casp3-3-SPR | /5Phos/AAGATAAACATCGTAGACTAGAAGACTCAACCTTCCT GTTAACGC |
| Casp9-1-SPL | /5Phos/TGGATCCTGCTTGGCTAAAAAACTCATCTCTCAGGTC ATACACTA |
| Casp9-1-SPR | /5Phos/AAGATAAACATCGTAGACTATCGAACTCAGGGGTTTA ACAGCCTC |
| Casp9-2-SPL | /5Phos/TGGGAAGGTGGAGTAGAAAAAACTCATCTCTCAGGTC ATACACTA |
| Casp9-2-SPR | /5Phos/AAGATAAACATCGTAGACTATCGAACTCAAACAGCAT TGGCAACC |
| Casp9-3-SPL | /5Phos/CAGAACCAATGTCCACAAAAAACTCATCTCTCAGGTC ATACACTA |
| Casp9-3-SPR | /5Phos/AAGATAAACATCGTAGACTATCGAACTCAACATCATG AGCTCCGC |
| Gpx4-1-SPL | /5Phos/CAATCATCGCGGGATGAAAAAACTCAGATCTCAGGTC ATACACTA |
| Gpx4-1-SPR | /5Phos/AAGATAAACATCGTAGACTAGACTACTCAGGAGCGCG CACAGCGC |
| Gpx4-2-SPL | /5Phos/ATATCGGGCATGCAGAAAAAAACTCAGATCTCAGGTC ATACACTA |
| Gpx4-2-SPR | /5Phos/AAGATAAACATCGTAGACTAGACTACTCAGTAAACCA CACTCAGC |
| Gpx4-3-SPL | /5Phos/TGGACTTTCATCCATTAAAAAACTCAGATCTCAGGTCA TACACTA |
| Gpx4-3-SPR | /5Phos/AAGATAAACATCGTAGACTAGACTACTCAGCCCCTGC CCTTGGGC |
| Slc7a11-1-SPL | /5Phos/TCCAGGGCGTATTACGAAAAAACTCAGATCTCAGGTC ATACACTA |
| Slc7a11-1-SPR | /5Phos/AAGATAAACATCGTAGACTATCCTACTCAATATCACAG CAGTAGC |
| Slc7a11-2-SPL | /5Phos/TTGTGGACATGAATCAAAAAAACTCAGATCTCAGGTC ATACACTA |
| Slc7a11-2-SPR | /5Phos/AAGATAAACATCGTAGACTATCCTACTCATGGCAGAG GAGTGTGC |
| Slc7a11-3-SPL | /5Phos/CGATGACGGTGCCGATAAAAAACTCAGATCTCAGGTC |

| | |
|---|---|
| | ATACACTA |
| Slc7a11-3-SPR | /5Phos/AAGATAAACATCGTAGACTATCCTACTCAATGAAGATGCCTGATC |
| Slc7a11-4-SPL | /5Phos/ATGAAGATGCCTGATCAAAAAACTCAGATCTCAGGTCATACACTA |
| Slc7a11-4-SPR | /5Phos/AAGATAAACATCGTAGACTATCCTACTCAAAGTTGAGGTAAAACC |
| Tfrc-1-SPL | /5Phos/CCAAAGCGTCTCTCTGAAAAAACTCAGAAGTCAGGTCATACACTA |
| Tfrc-1-SPR | /5Phos/AAGATAAACATCGTAGACTAGACTACTCAGCCGCAACACCAGCAC |
| Tfrc-2-SPL | /5Phos/CAGAGAGGGCATTTGCAAAAAACTCAGAAGTCAGGTCATACACTA |
| Tfrc-2-SPR | /5Phos/AAGATAAACATCGTAGACTAGACTACTCAATATTCCAAATGTCAC |
| Tfrc-3-SPL | /5Phos/TGCTTGATGGTGTCAGAAAAAACTCAGAAGTCAGGTCATACACTA |
| Tfrc-3-SPR | /5Phos/AAGATAAACATCGTAGACTAGACTACTCATGTATTCTGGCTCAGC |
| Pkm-1-SPL | /5Phos/CATGGCTGCATGGAGCAAAAAACTCAGAAGTCAGGTCATACACTA |
| Pkm-1-SPR | /5Phos/AAGATAAACATCGTAGACTAAGGAACTCACCAGGAAGGTGTCAGC |
| Pkm-2-SPL | /5Phos/CAGGTCACCACGAGCCAAAAAACTCAGAAGTCAGGTCATACACTA |
| Pkm-2-SPR | /5Phos/AAGATAAACATCGTAGACTAAGGAACTCACAGGAATCTCAATGCC |
| Pkm-3-SPL | /5Phos/TTGAAGGAGGCCTCCAAAAAAACTCAGAAGTCAGGTCATACACTA |
| Pkm-3-SPR | /5Phos/AAGATAAACATCGTAGACTAAGGAACTCAGGCCCCACTGCAGCAC |
| Nfe2l2-1-SPL | /5Phos/TTGTTTTCGGTATTAAAAAAAACTCAGACTTCAGGTCATACACTA |
| Nfe2l2-1-SPR | /5Phos/AAGATAAACATCGTAGACTAAGGAACTCAAGTATCAGCCAGCTGC |
| Nfe2l2-2-SPL | /5Phos/CCTAAGCTCATCTCGTAAAAAACTCAGACTTCAGGTCATACACTA |
| Nfe2l2-2-SPR | /5Phos/AAGATAAACATCGTAGACTAAGGAACTCATATGGAGAGCTTTTGC |
| Nfe2l2-3-SPL | /5Phos/TGGATGTGCTGGGCCGAAAAAACTCAGACTTCAGGTCATACACTA |
| Nfe2l2-3-SPR | /5Phos/AAGATAAACATCGTAGACTAAGGAACTCATCCACTGGTGTCTGTC |
| Pink1-1-SPL | /5Phos/CGGTAGCGGTCCGGGAAAAAAACTCAGACTTCAGGT |

| | |
|---|---|
| | CATACACTA |
| Pink1-1-SPR | /5Phos/AAGATAAACATCGTAGACTACTGAACTCACGACTGGC GGAAGAAG |
| Pink1-2-SPL | /5Phos/TGTGGACACCTCAGGGAAAAAACTCAGACTTCAGGTC ATACACTA |
| Pink1-2-SPR | /5Phos/AAGATAAACATCGTAGACTACTGAACTCATGGGGCCA GAATGGGC |
| Pink1-3-SPL | /5Phos/TCTTCTCTCTCAGCCTAAAAAACTCAGACTTCAGGTC ATACACTA |
| Pink1-3-SPR | /5Phos/AAGATAAACATCGTAGACTACTGAACTCATTTGTCTCC ACGCAGC |
| Prkn-1-SPL | /5Phos/AACCACTTCCTTGAGCAAAAAACTCAGAGATCAGGTC ATACACTA |
| Prkn-1-SPR | /5Phos/AAGATAAACATCGTAGACTATCGAACTCACCCCCTGT CGCTTAGC |
| Prkn-2-SPL | /5Phos/TGGCACTCACCACTCAAAAAAACTCAGAGATCAGGTC ATACACTA |
| Prkn-2-SPR | /5Phos/AAGATAAACATCGTAGACTATCGAACTCAAGGGCAGT CTGGAGAC |
| Prkn-3-SPL | /5Phos/CATTTGCAGCACGCATAAAAAACTCAGAGATCAGGTC ATACACTA |
| Prkn-3-SPR | /5Phos/AAGATAAACATCGTAGACTATCGAACTCAGGCACAGC ACACCTCC |
| Map1lc3a-1-SPL | /5Phos/TCCTTACAGCGGTCGGAAAAAACTCAGAGATCAGGTC ATACACTA |
| Map1lc3a-1-SPR | /5Phos/AAGATAAACATCGTAGACTAAGTCACTCAGCGGATCT GCTGCACC |
| Map1lc3a-2-SPL | /5Phos/TCGATGATCACCGGGAAAAAAACTCAGAGATCAGGTC ATACACTA |
| Map1lc3a-2-SPR | /5Phos/AAGATAAACATCGTAGACTAAGTCACTCACTCACCCT TGTAGCGC |
| Map1lc3a-3-SPL | /5Phos/GCGTAGACCATGTAGAAAAAAACTCAGAGATCAGGTC ATACACTA |
| Map1lc3a-3-SPR | /5Phos/AAGATAAACATCGTAGACTAAGTCACTCAGAAGGTTT CTTGGGAG |
| Sqstm1-1-SPL | /5Phos/CCATCCTCATCGCGGTAAAAAACTCAAGAGTCAGGTC ATACACTA |
| Sqstm1-1-SPR | /5Phos/AAGATAAACATCGTAGACTACTTCACTCAAAAGGCAA CCAAGTCC |
| Sqstm1-2-SPL | /5Phos/ATGTTTCGGGGTGCCTAAAAAACTCAAGAGTCAGGTC ATACACTA |
| Sqstm1-2-SPR | /5Phos/AAGATAAACATCGTAGACTACTTCACTCACACATTGG GGTGCACC |
| Sqstm1-3-SPL | /5Phos/AGCCCTGTGGGTCCTTAAAAAAACTCAAGAGTCAGGT |

| | |
|---|---|
| | CATACACTA |
| Sqstm1-3-SPR | /5Phos/AAGATAAACATCGTAGACTACTTCACTCATAGGGCAGCTTCCTTC |
| Tomm20-1-SPL | /5Phos/GAAGCCTGTTCTTGAAAAAAAACTCACTGATCAGGTCATACACTA |
| Tomm20-1-SPR | /5Phos/AAGATAAACATCGTAGACTATCCTACTCATTCTTTCTTCGTTCTC |
| Tomm20-2-SPL | /5Phos/CCTTCTCGTAGTCACCAAAAAACTCACTGATCAGGTCATACACTA |
| Tomm20-2-SPR | /5Phos/AAGATAAACATCGTAGACTATCCTACTCAGTCAGGTGGTCCACAC |
| Tomm20-3-SPL | /5Phos/TGACTAATGGTCGGAAAAAAAACTCACTGATCAGGTCATACACTA |
| Tomm20-3-SPR | /5Phos/AAGATAAACATCGTAGACTATCCTACTCAAGCACTTACAATTCTC |
| Gsdmd-1-SPL | /5Phos/TGACAACATCACTCTGAAAAAACTCATCTCTCAGGTCATACACTA |
| Gsdmd-1-SPR | /5Phos/AAGATAAACATCGTAGACTAAGCTACTCATCCCTTCTCCCATGCC |
| Gsdmd-2-SPL | /5Phos/ACAAACAGGTCATCCCAAAAAACTCATCTCTCAGGTCATACACTA |
| Gsdmd-2-SPR | /5Phos/AAGATAAACATCGTAGACTAAGCTACTCACAGCACCTCGGTCACC |
| Gsdmd-3-SPL | /5Phos/TTCACCTCAGCATACAAAAAAACTCATCTCTCAGGTCATACACTA |
| Gsdmd-3-SPR | /5Phos/AAGATAAACATCGTAGACTAAGCTACTCATTCTGAGGAGCAAGCC |
| Nlrp3-1-SPL | /5Phos/TGTTGATCGCAGCAAAAAAAAACTCACTCTTCAGGTCATACACTA |
| Nlrp3-1-SPR | /5Phos/AAGATAAACATCGTAGACTAGAGAACTCATCCCAGAGGTCTCGCC |
| Nlrp3-2-SPL | /5Phos/TTGCTTGGATGCTCCTAAAAAACTCACTCTTCAGGTCATACACTA |
| Nlrp3-2-SPR | /5Phos/AAGATAAACATCGTAGACTAGAGAACTCAATGCTCCCGCTCCTGC |
| Nlrp3-3-SPL | /5Phos/TCGTACAGGCAGTAGAAAAAAACTCACTCTTCAGGTCATACACTA |
| Nlrp3-3-SPR | /5Phos/AAGATAAACATCGTAGACTAGAGAACTCAGTCTTCCTCCTGCATC |
| Tgfb1-1-SPL | /5Phos/AGCTGCCGCACACAGCAAAAAACTCATCGATCAGGTCATACACTA |
| Tgfb1-1-SPR | /5Phos/AAGATAAACATCGTAGACTATCTCACTCACCTAAAGTCAATGTAC |
| Tgfb1-2-SPL | /5Phos/TGACGTATTGAAGAACAAAAAACTCATCGATCAGGTC |

| | |
|---|---|
| | ATACACTA |
| Tgfb1-2-SPR | /5Phos/AAGATAAACATCGTAGACTATCTCACTCACTGCTTCC CGAATGTC |
| Tgfb1-3-SPL | /5Phos/TGCCGTACAACTCCAGAAAAAACTCATCGATCAGGTC ATACACTA |
| Tgfb1-3-SPR | /5Phos/AAGATAAACATCGTAGACTATCTCACTCATCCTTGGTT CAGCCAC |
| Tef-1-SPL | /5Phos/TTGAGGACCTCAGGCAAAAAAACTCATCCTTCAGGTC ATACACTA |
| Tef-1-SPR | /5Phos/AAGATAAACATCGTAGACTAAGAGACTCAATGCTCCA GCAGGGAC |
| Tef-2-SPL | /5Phos/TCTGCCACAGGCAGCAAAAAAACTCATCCTTCAGGTC ATACACTA |
| Tef-2-SPR | /5Phos/AAGATAAACATCGTAGACTAAGAGACTCACTCCTTTC CTTCAAGC |
| Tef-3-SPL | /5Phos/TGCAAACCTGTGCTTCAAAAAACTCATCCTTCAGGTC ATACACTA |
| Tef-3-SPR | /5Phos/AAGATAAACATCGTAGACTAAGAGACTCAGCTTCAGG TCCTCCTC |
| Tef-4-SPL | /5Phos/AGGAACGCTGCCCGGAAAAAAACTCATCCTTCAGGT CATACACTA |
| Tef-4-SPR | /5Phos/AAGATAAACATCGTAGACTAAGAGACTCATGTGTTCT CCTTCTCC |
| Syt1-1-SPL | /5Phos/CTTCCCCAGGACTGGCAAAAAACTCATCTCTCAGGTC ATACACTA |
| Syt1-1-SPR | /5Phos/AAGATAAACATCGTAGACTACTGAACTCAAAGGCATC TTCCTTCC |
| Syt1-2-SPL | /5Phos/ACCAGCAGCTGGTTATAAAAAACTCATCTCTCAGGTC ATACACTA |
| Syt1-2-SPR | /5Phos/AAGATAAACATCGTAGACTACTGAACTCAAGCCTGGA TGATTCCC |
| Syt1-3-SPL | /5Phos/GTGCTTGGAGAAGCGGAAAAAACTCATCTCTCAGGTC ATACACTA |
| Syt1-3-SPR | /5Phos/AAGATAAACATCGTAGACTACTGAACTCAACTCTCCA ATGATGTC |
| Syt1-4-SPL | /5Phos/TTGCACTTTCTGGATTAAAAAACTCATCTCTCAGGTCA TACACTA |
| Syt1-4-SPR | /5Phos/AAGATAAACATCGTAGACTACTGAACTCAAAACAGTT ACCACCAC |
| Comt-1-SPL | /5Phos/AGGAGGAAGGCCAGCAAAAAAACTCATCGATCAGGT CATACACTA |
| Comt-1-SPR | /5Phos/AAGATAAACATCGTAGACTAAGGAACTCATAGGTGTC GCAGGAGC |
| Comt-2-SPL | /5Phos/CAGGCTTTGCGTGTTGAAAAAACTCATCGATCAGGTC |

| | |
|---|---|
| | ATACACTA |
| Comt-2-SPR | /5Phos/AAGATAAACATCGTAGACTAAGGAACTCAACGCTCTG GGGGTCTC |
| Comt-3-SPL | /5Phos/AGGGTTAATCTCCATGAAAAAACTCATCGATCAGGTC ATACACTA |
| Comt-3-SPR | /5Phos/AAGATAAACATCGTAGACTAAGGAACTCATGATGGCA GCGTAGTC |
| Comt-4-SPL | /5Phos/GGTCTTTCCAGTGGTCAAAAAACTCATCGATCAGGTC ATACACTA |
| Comt-4-SPR | /5Phos/AAGATAAACATCGTAGACTAAGGAACTCAGTGTCTGG AAGGTAGC |
| Dbh-1-SPL | /5Phos/CACAGCAGTGCCGTACAAAAAACTCATCGATCAGGTC ATACACTA |
| Dbh-1-SPR | /5Phos/AAGATAAACATCGTAGACTAAGTCACTCATGACCAGG AAGATGGC |
| Dbh-2-SPL | /5Phos/ATGGATGGAGTGGGGAAAAAAACTCATCGATCAGGT CATACACTA |
| Dbh-2-SPR | /5Phos/AAGATAAACATCGTAGACTAAGTCACTCATTGTACATC CTCAGGC |
| Dbh-3-SPL | /5Phos/CGGTCAACACAAAGGCAAAAAACTCATCGATCAGGTC ATACACTA |
| Dbh-3-SPR | /5Phos/AAGATAAACATCGTAGACTAAGTCACTCATTGTCTGT GCAGTAGC |
| Dbh-4-SPL | /5Phos/CAGGGCACGGAGGAGAAAAAAACTCATCGATCAGGT CATACACTA |
| Dbh-4-SPR | /5Phos/AAGATAAACATCGTAGACTAAGTCACTCACCGATTGA AAGAGTTC |
| Cdkn1a-1-SPL | /5Phos/ACACTTTGCTCCTGTGAAAAAACTCATCGATCAGGTC ATACACTA |
| Cdkn1a-1-SPR | /5Phos/AAGATAAACATCGTAGACTACTGAACTCACCGAAGAG ACAACGGC |
| Cdkn1a-2-SPL | /5Phos/CCTCCAGCGGCGTCTCAAAAAACTCATCGATCAGGTC ATACACTA |
| Cdkn1a-2-SPR | /5Phos/AAGATAAACATCGTAGACTACTGAACTCATCCCAGAC GAAGTTGC |
| Cdkn1a-3-SPL | /5Phos/TGGGCCTCTTGTCCCCAAAAAACTCATCGATCAGGTC ATACACTA |
| Cdkn1a-3-SPR | /5Phos/AAGATAAACATCGTAGACTACTGAACTCAAGGGCAGA GGAAGTAC |
| Cdkn1a-4-SPL | /5Phos/AGGCCGCTCAGACACCAAAAAACTCATCGATCAGGT CATACACTA |
| Cdkn1a-4-SPR | /5Phos/AAGATAAACATCGTAGACTACTGAACTCACACCCGGG GAATCTTC |
| Cks1b-1-SPL | /5Phos/CTCGTCGTCGTATTTGAAAAAACTCATCGATCAGGTC |

| | |
|---|---|
| | ATACACTA |
| Cks1b-1-SPR | /5Phos/AAGATAAACATCGTAGACTATCCTACTCAGCCGGTATTCGAACTC |
| Cks1b-2-SPL | /5Phos/AGATGGGTTTTCGGGAAAAAAACTCATCGATCAGGTCATACACTA |
| Cks1b-2-SPR | /5Phos/AAGATAAACATCGTAGACTATCCTACTCATTCAGATTCAGACATC |
| Cks1b-3-SPL | /5Phos/CGTCGGAACAGTAAGAAAAAAACTCATCGATCAGGTCATACACTA |
| Cks1b-3-SPR | /5Phos/AAGATAAACATCGTAGACTATCCTACTCACTTCTTGGGCAGTGGC |
| Nrgn-1-SPL | /5Phos/ATATCGTCGTCTGGCTAAAAAACTCACTCTTCGATCGATAGCTAAACACTA |
| Nrgn-1-SPR | /5Phos/AAGATACTTGGTATAATCGCTTCTCACTCAAAAAACAGCGGGATGTCAAGA |
| Nrgn-2-SPL | /5Phos/TATCTTCTTCCTCGCCAAAAAACTCACTCTTCGATCGATAGCTAAACACTA |
| Nrgn-2-SPR | /5Phos/AAGATACTTGGTATAATCGCTTCTCACTCAAAAAACACACTCTCCGCTCTT |
| Cx3cr1-1-SPL | /5Phos/AGACGGACAGGAAGATAAAAAACTCAAGCTTCGATCGATAGCTAAACACTA |
| Cx3cr1-1-SPR | /5Phos/AAGATACTTGGTATAATCGCTAGAGACTCAAAAAAAAGACGAGGGCGTAGA |
| Cx3cr1-2-SPL | /5Phos/AGAATATGCCCCCAAAAAAAAACTCAAGCTTCGATCGATAGCTAAACACTA |
| Cx3cr1-2-SPR | /5Phos/AAGATACTTGGTATAATCGCTAGAGACTCAAAAAACTGATGACGGTGATGA |
| Cx3cr1-3-SPL | /5Phos/AACAGCGTCTGGATGAAAAAAACTCAAGCTTCGATCGATAGCTAAACACTA |
| Cx3cr1-3-SPR | /5Phos/AAGATACTTGGTATAATCGCTAGAGACTCAAAAAAGCGATTCTTGCAGGAA |
| Cx3cr1-4-SPL | /5Phos/TCGTTGTCCTTTCTCTAAAAAACTCAAGCTTCGATCGATAGCTAAACACTA |
| Cx3cr1-4-SPR | /5Phos/AAGATACTTGGTATAATCGCTAGAGACTCAAAAAAGTAGTCACCCAGACAC |
| Cx3cr1-5-SPL | /5Phos/AGGTGGCCACAAAGAGAAAAAACTCAAGCTTCGATCGATAGCTAAACACTA |
| Cx3cr1-5-SPR | /5Phos/AAGATACTTGGTATAATCGCTAGAGACTCAAAAAATGAGTCCAGAAGGGCA |
| Cx3cr1-6-SPL | /5Phos/TCATGTCACAACTGGGAAAAAACTCAAGCTTCGATCGATAGCTAAACACTA |
| Cx3cr1-6-SPR | /5Phos/AAGATACTTGGTATAATCGCTAGAGACTCAAAAAAAACCTCAGGTCCCTCT |
| Tmem119-1-SPL | /5Phos/AGCAGCAGAGACAGGAAAAAAACTCAAGTCTCGATC |

| | |
|---|---|
| | GATAGCTAAACACTA |
| Tmem119-1-SPR | /5Phos/AAGATACTTGGTATAATCGCTAGTCACTCAAAAAAACA<br>GGCCTCGCAAGT |
| Tmem119-2-SPL | /5Phos/ATCCATGATCCCTTCCAAAAAACTCAAGTCTCGATCG<br>ATAGCTAAACACTA |
| Tmem119-2-SPR | /5Phos/AAGATACTTGGTATAATCGCTAGTCACTCAAAAAACG<br>TACTGCCGGAAGAA |
| Tmem119-3-SPL | /5Phos/TGCTCCCTGGGATTCAAAAAACTCAAGTCTCGATCGA<br>TAGCTAAACACTA |
| Tmem119-3-SPR | /5Phos/AAGATACTTGGTATAATCGCTAGTCACTCAAAAAATTT<br>CAGGAGGACCAGT |
| Tmem119-4-SPL | /5Phos/TGAAGGCTGGAGTCCCAAAAAACTCAAGTCTCGATCG<br>ATAGCTAAACACTA |
| Tmem119-4-SPR | /5Phos/AAGATACTTGGTATAATCGCTAGTCACTCAAAAAACTC<br>TCCGGTGTGGGAC |
| Tmem119-5-SPL | /5Phos/AGGCCTTCTTCATGCCAAAAAACTCAAGTCTCGATCG<br>ATAGCTAAACACTA |
| Tmem119-5-SPR | /5Phos/AAGATACTTGGTATAATCGCTAGTCACTCAAAAAACT<br>GATGGGAGGTGTCC |
| Tmem119-6-SPL | /5Phos/TCCTCTGGGACCCCCAAAAAAACTCAAGTCTCGATCG<br>ATAGCTAAACACTA |
| Tmem119-6-SPR | /5Phos/AAGATACTTGGTATAATCGCTAGTCACTCAAAAAACTC<br>TGCTGAGACCGAC |
| Hexb-1-SPL | /5Phos/TGCTGAAGTCCTCCGAAAAAACTCAAGTCTCGATCGA<br>TAGCTAAACACTA |
| Hexb-1-SPR | /5Phos/AAGATACTTGGTATAATCGCTGACTACTCAAAAAATTG<br>GGACTGTGGTCGA |
| Hexb-2-SPL | /5Phos/TGAGGACGGCTACTGAAAAAACTCAAGTCTCGATCGA<br>TAGCTAAACACTA |
| Hexb-2-SPR | /5Phos/AAGATACTTGGTATAATCGCTGACTACTCAAAAAACAA<br>ACGCTGTTGGCCT |
| Hexb-3-SPL | /5Phos/ACAGGCCCAAACACTTAAAAAACTCAAGTCTCGATCG<br>ATAGCTAAACACTA |
| Hexb-3-SPR | /5Phos/AAGATACTTGGTATAATCGCTGACTACTCAAAAAATGT<br>TTACAGTTGGGTCT |
| Hexb-4-SPL | /5Phos/AGGGCCATGATGTCTCAAAAAACTCAAGTCTCGATCG<br>ATAGCTAAACACTA |
| Hexb-4-SPR | /5Phos/AAGATACTTGGTATAATCGCTGACTACTCAAAAAACA<br>GCTCGAAATCTAGC |
| Hexb-5-SPL | /5Phos/CGAGGGTAATGGAGACAAAAAACTCAAGTCTCGATC<br>GATAGCTAAACACTA |
| Hexb-5-SPR | /5Phos/AAGATACTTGGTATAATCGCTGACTACTCAAAAAAGA<br>CTCGCACTCTGACT |
| Hexb-6-SPL | /5Phos/AAAGAGTAGCTTCCCTAAAAAACTCAAGTCTCGATCG |

| | |
|---|---|
| | ATAGCTAAACACTA |
| Hexb-6-SPR | /5Phos/AAGATACTTGGTATAATCGCTGACTACTCAAAAAATGT ATAGACATGAGAC |
| Slc17a7-1-SPL | /5Phos/ACTGGGCTTTCTGCACAAAAAACTCACTTCTCGATCG ATAGCTAAACACTA |
| Slc17a7-1-SPR | /5Phos/AAGATACTTGGTATAATCGCTCTTCACTCAAAAAATCT GGATCCCAGTTGA |
| Slc17a7-2-SPL | /5Phos/TGTATTTGCGCTCCTCAAAAAACTCACTTCTCGATCG ATAGCTAAACACTA |
| Slc17a7-2-SPR | /5Phos/AAGATACTTGGTATAATCGCTCTTCACTCAAAAAACC GATGGCATCCTCAA |
| Slc17a7-3-SPL | /5Phos/TCACTTTCGTCACTGCAAAAAACTCACTTCTCGATCG ATAGCTAAACACTA |
| Slc17a7-3-SPR | /5Phos/AAGATACTTGGTATAATCGCTCTTCACTCAAAAAAAGC CTCGTCCTCCATT |
| Foxj1-1-SPL | /5Phos/ACCCCAGGGGGCAGCGAAAAAACTCAGACTTCGATC GATAGCTAAACACTA |
| Foxj1-1-SPR | /5Phos/AAGATACTTGGTATAATCGCTGACTACTCAAAAAATCA CGGTCAGAGGCCC |
| Foxj1-2-SPL | /5Phos/CATGTGAGCTGGGGCTAAAAAACTCAGACTTCGATCG ATAGCTAAACACTA |
| Foxj1-2-SPR | /5Phos/AAGATACTTGGTATAATCGCTGACTACTCAAAAAAGTA AGATCCACATCTC |
| Foxj1-3-SPL | /5Phos/CCCCAGGTAGCAGGGCAAAAAACTCAGACTTCGATC GATAGCTAAACACTA |
| Foxj1-3-SPR | /5Phos/AAGATACTTGGTATAATCGCTGACTACTCAAAAAACT GCTCCGCTGGAGGT |
| Lefty1-1-SPL | /5Phos/AGTCCTCACGTGCGAGAAAAAACTCATCCTTCGATCG ATAGCTAAACACTA |
| Lefty1-1-SPR | /5Phos/AAGATACTTGGTATAATCGCTCTTCACTCAAAAAAGCA GGGCCACATACTG |
| Lefty1-2-SPL | /5Phos/TGTCAGGAACCCTGGCAAAAAACTCATCCTTCGATCG ATAGCTAAACACTA |
| Lefty1-2-SPR | /5Phos/AAGATACTTGGTATAATCGCTCTTCACTCAAAAAATGC CCACACATTCATA |
| Lefty1-3-SPL | /5Phos/TCCTTGGGGAAGCCACAAAAAACTCATCCTTCGATCG ATAGCTAAACACTA |
| Lefty1-3-SPR | /5Phos/AAGATACTTGGTATAATCGCTCTTCACTCAAAAAATTG CATGAAAGGCACA |
| Opalin-1-SPL | /5Phos/AACAGCAAAGCCACCAAAAAAACTCATCTCTCGATCG ATAGCTAAACACTA |
| Opalin-1-SPR | /5Phos/AAGATACTTGGTATAATCGCTTCGAACTCAAAAAATC GCTGGATCAAGGTA |
| Opalin-2-SPL | /5Phos/TCATGTGTGGGTGACCAAAAAACTCATCTCTCGATCG |

| | |
|---|---|
| | ATAGCTAAACACTA |
| Opalin-2-SPR | /5Phos/AAGATACTTGGTATAATCGCTTCGAACTCAAAAAACC CCGTGGGTTCATT |
| Opalin-3-SPL | /5Phos/TTCTAGGCTCAGGCTGAAAAAACTCATCTCTCGATCG ATAGCTAAACACTA |
| Opalin-3-SPR | /5Phos/AAGATACTTGGTATAATCGCTTCGAACTCAAAAAATGA TGCGGTGACATCA |
| Cldn11-1-SPL | /5Phos/AACCCACCACCTGAAGAAAAAACTCATCGATCGATCG ATAGCTAAACACTA |
| Cldn11-1-SPR | /5Phos/AAGATACTTGGTATAATCGCTTCTCACTCAAAAAAACG AAGCTCGTGACGA |
| Cldn11-2-SPL | /5Phos/AGCCTGGAAGGATGAGAAAAAACTCATCGATCGATC GATAGCTAAACACTA |
| Cldn11-2-SPR | /5Phos/AAGATACTTGGTATAATCGCTTCTCACTCAAAAAACTA CAAGCCTGCACGT |
| Cldn11-3-SPL | /5Phos/AGCTCACGATGGTGATAAAAAACTCATCGATCGATCG ATAGCTAAACACTA |
| Cldn11-3-SPR | /5Phos/AAGATACTTGGTATAATCGCTTCTCACTCAAAAAATAC AGCGAGTAGCCAA |
| Lct-1-SPL | /5Phos/ACTGTCTCATGCTGCTAAAAAACTCATCGATCGATCG ATAGCTAAACACTA |
| Lct-1-SPR | /5Phos/AAGATACTTGGTATAATCGCTGAGAACTCAAAAAAAG GGAATGAGGACACA |
| Lct-2-SPL | /5Phos/ATTCGAGGCCAGGAGAAAAAAACTCATCGATCGATCG ATAGCTAAACACTA |
| Lct-2-SPR | /5Phos/AAGATACTTGGTATAATCGCTGAGAACTCAAAAAATCT TCCCGTTGGGAAA |
| Lct-3-SPL | /5Phos/TCATCACCTCAGGGTAAAAAAACTCATCGATCGATCG ATAGCTAAACACTA |
| Lct-3-SPR | /5Phos/AAGATACTTGGTATAATCGCTGAGAACTCAAAAAATC ACGAATCCGTGTCT |
| Adora2a-1-SPL | /5Phos/ATCGCAATGATGCCCTAAAAAACTCATCCTTCGATCG ATAGCTAAACACTA |
| Adora2a-1-SPR | /5Phos/AAGATACTTGGTATAATCGCTAGGAACTCAAAAAATG ACAGCACCCAGCAA |
| Adora2a-2-SPL | /5Phos/TTGCTGTGGGAGAGGAAAAAAACTCATCCTTCGATCG ATAGCTAAACACTA |
| Adora2a-2-SPR | /5Phos/AAGATACTTGGTATAATCGCTAGGAACTCAAAAAAGG GGTTGACAACGGAA |
| Adora2a-3-SPL | /5Phos/AGACCATGAGGCTGTTAAAAAACTCATCCTTCGATCG ATAGCTAAACACTA |
| Adora2a-3-SPR | /5Phos/AAGATACTTGGTATAATCGCTAGGAACTCAAAAAAAA GGGGCAAACTCTGA |
| Drd2-1-SPL | /5Phos/AGTTGTAGTGGGGCCTAAAAAACTCATCAGTCGATCG |

| | |
|---|---|
| | ATAGCTAAACACTA |
| Drd2-1-SPR | /5Phos/AAGATACTTGGTATAATCGCTTCAGACTCAAAAAAAG CAGCATGGCATAGT |
| Drd2-2-SPL | /5Phos/TGGTGAAGGACAGGACAAAAAACTCATCAGTCGATC GATAGCTAAACACTA |
| Drd2-2-SPR | /5Phos/AAGATACTTGGTATAATCGCTTCAGACTCAAAAAAAG CAGTGGGCAAGAGA |
| Drd2-3-SPL | /5Phos/TGAAGAAGGGCAGCCAAAAAAACTCATCAGTCGATC GATAGCTAAACACTA |
| Drd2-3-SPR | /5Phos/AAGATACTTGGTATAATCGCTTCAGACTCAAAAAATTC AGGATGTGCGTGA |
| Shox2-1-SPL | /5Phos/ACGCCGTAAGTTCTTCAAAAAACTCAGATCTCGATCG ATAGCTAAACACTA |
| Shox2-1-SPR | /5Phos/AAGATACTTGGTATAATCGCTTCTCACTCAAAAAAAAA GACTTGGAGACGA |
| Shox2-2-SPL | /5Phos/TCCTTCAGTTCAGGGGAAAAAACTCAGATCTCGATCG ATAGCTAAACACTA |
| Shox2-2-SPR | /5Phos/AAGATACTTGGTATAATCGCTTCTCACTCAAAAAACG CATCGTCTTTGCGA |
| Shox2-3-SPL | /5Phos/TGACATAGGGTGCAACAAAAAACTCAGATCTCGATCG ATAGCTAAACACTA |
| Shox2-3-SPR | /5Phos/AAGATACTTGGTATAATCGCTTCTCACTCAAAAAACCT TAAAGCACCTACGT |
| Gbx2-1-SPL | /5Phos/ATGCTGAAGGCGGTACAAAAAACTCAGAGATCGATC GATAGCTAAACACTA |
| Gbx2-1-SPR | /5Phos/AAGATACTTGGTATAATCGCTGATCACTCAAAAAACC GATCAGCGAGTCT |
| Gbx2-2-SPL | /5Phos/AGGCTTTGCCGTCTTCAAAAAACTCAGAGATCGATCG ATAGCTAAACACTA |
| Gbx2-2-SPR | /5Phos/AAGATACTTGGTATAATCGCTGATCACTCAAAAAACC CTCCTTGGCCAAGA |
| Gbx2-3-SPL | /5Phos/TCGCTCTCCAGAGAGAAAAAAACTCAGAGATCGATCG ATAGCTAAACACTA |
| Gbx2-3-SPR | /5Phos/AAGATACTTGGTATAATCGCTGATCACTCAAAAAACT GAGCTGTAATCCACA |
| Gpx3-1-SPL | /5Phos/TGGTACCACTCATACCAAAAAACTCAGACTTCGATCG ATAGCTAAACACTA |
| Gpx3-1-SPR | /5Phos/AAGATACTTGGTATAATCGCTCTGAACTCAAAAAAGC TCCATACTCGTAGA |
| Gpx3-2-SPL | /5Phos/ACTTGAGACTGGGGAGAAAAAACTCAGACTTCGATCG ATAGCTAAACACTA |
| Gpx3-2-SPR | /5Phos/AAGATACTTGGTATAATCGCTCTGAACTCAAAAAACC ACCTGGTCGAACAT |
| Lmo3-1-SPL | /5Phos/AAGGTTAGCCTTCGTGAAAAAACTCAGACTTCGATCG |

| | |
|---|---|
| | ATAGCTAAACACTA |
| Lmo3-1-SPR | /5Phos/AAGATACTTGGTATAATCGCTTCAGACTCAAAAAACTC TGCGACAAAGGAT |
| Lmo3-2-SPL | /5Phos/TTATCCTTGGCACGCAAAAAAACTCAGACTTCGATCG ATAGCTAAACACTA |
| Lmo3-2-SPR | /5Phos/AAGATACTTGGTATAATCGCTTCAGACTCAAAAAAGT CCAGGTGGTACACA |
| Slc30a3-1-SPL | /5Phos/ATCCAGAGGGAGACCAAAAAAACTCAGAGATCGATC GATAGCTAAACACTA |
| Slc30a3-1-SPR | /5Phos/AAGATACTTGGTATAATCGCTCTCTACTCAAAAAAGA GGATGCCAGTGACT |
| Slc30a3-2-SPL | /5Phos/TTGGGTATCCATGCCCAAAAAACTCAGAGATCGATCG ATAGCTAAACACTA |
| Slc30a3-2-SPR | /5Phos/AAGATACTTGGTATAATCGCTCTCTACTCAAAAAAGTG TTTCCCAGGGACA |
| Slc30a3-3-SPL | /5Phos/TAGCCAGGTGTGCAGAAAAAAACTCAGAGATCGATC GATAGCTAAACACTA |
| Slc30a3-3-SPR | /5Phos/AAGATACTTGGTATAATCGCTCTCTACTCAAAAAATCA GCCGTGGAGTCAA |
| Epop-1-SPL | /5Phos/TGTTCCTGGCGGAGATAAAAAACTCAGAGATCGATCG ATAGCTAAACACTA |
| Epop-1-SPR | /5Phos/AAGATACTTGGTATAATCGCTTCGAACTCAAAAAAGAT CAGGCGGTTGAAA |
| Epop-2-SPL | /5Phos/TGAAGGTGCTCGATGTAAAAAACTCAGAGATCGATCG ATAGCTAAACACTA |
| Epop-2-SPR | /5Phos/AAGATACTTGGTATAATCGCTTCGAACTCAAAAAAAA GCAGTCAAGGAGGC |
| Epop-3-SPL | /5Phos/AGGATTTGAGACCCGGAAAAAACTCAGAGATCGATC GATAGCTAAACACTA |
| Epop-3-SPR | /5Phos/AAGATACTTGGTATAATCGCTTCGAACTCAAAAAATCC AAGTTGACCCACC |
| Cobl-1-SPL | /5Phos/TGGTGAGTTGGGTGTTAAAAAACTCAGATCTCGATCG ATAGCTAAACACTA |
| Cobl-1-SPR | /5Phos/AAGATACTTGGTATAATCGCTAGCTACTCAAAAAAAC CGAGAATGCAAGGA |
| Cobl-2-SPL | /5Phos/ATGGTCCTGCTTCCGTAAAAAACTCAGATCTCGATCG ATAGCTAAACACTA |
| Cobl-2-SPR | /5Phos/AAGATACTTGGTATAATCGCTAGCTACTCAAAAAAAG CTGATGGTGGGGAT |
| Cobl-3-SPL | /5Phos/TAGCTTCCATCAGGGCAAAAAACTCAGATCTCGATCG ATAGCTAAACACTA |
| Cobl-3-SPR | /5Phos/AAGATACTTGGTATAATCGCTAGCTACTCAAAAAACCT CCTGATGAGTGGA |
| Scn5a-1-SPL | /5Phos/AAAGTTACGCACGCACAAAAAACTCAGATCTCGATCG |

| | |
|---|---|
| | ATAGCTAAACACTA |
| Scn5a-1-SPR | /5Phos/AAGATACTTGGTATAATCGCTCTAGACTCAAAAAATG CCATTGAGCTCAGT |
| Scn5a-2-SPL | /5Phos/TGGTGAGGTCAGCAAAAAAAAACTCAGATCTCGATCG ATAGCTAAACACTA |
| Scn5a-2-SPR | /5Phos/AAGATACTTGGTATAATCGCTCTAGACTCAAAAAAAC GATGCACATGGTGA |
| Scn5a-3-SPL | /5Phos/TAAGGTGTTGGCCACGAAAAAACTCAGATCTCGATCG ATAGCTAAACACTA |
| Scn5a-3-SPR | /5Phos/AAGATACTTGGTATAATCGCTCTAGACTCAAAAAACC ATCTCGGCAAAGCC |
| Aldh3b2-1-SPL | /5Phos/TCTTGCAGCAGCTCTTAAAAAACTCATCTCTCGATCG ATAGCTAAACACTA |
| Aldh3b2-1-SPR | /5Phos/AAGATACTTGGTATAATCGCTAGAGACTCAAAAAAGT CCTTAGCCAGTGCA |
| Aldh3b2-2-SPL | /5Phos/TTCACCACCTGGTTGTAAAAAACTCATCTCTCGATCG ATAGCTAAACACTA |
| Aldh3b2-2-SPR | /5Phos/AAGATACTTGGTATAATCGCTAGAGACTCAAAAAAGC GTTCCAACATCTGA |
| Aldh3b2-3-SPL | /5Phos/TAAGGCGGGTAACGGAAAAAAACTCATCTCTCGATCG ATAGCTAAACACTA |
| Aldh3b2-3-SPR | /5Phos/AAGATACTTGGTATAATCGCTAGAGACTCAAAAAATT GGTTCCAGGGACCA |
| Resp18-1-SPL | /5Phos/TGCCTGGACGACTGTAAAAAAACTCAAGGATCGATCG ATAGCTAAACACTA |
| Resp18-1-SPR | /5Phos/AAGATACTTGGTATAATCGCTAGGAACTCAAAAAATTT ATGTCGCTGCAGC |
| Resp18-2-SPL | /5Phos/TGCCTTCGGGTACAATAAAAAACTCAAGGATCGATCG ATAGCTAAACACTA |
| Resp18-2-SPR | /5Phos/AAGATACTTGGTATAATCGCTAGGAACTCAAAAAATC ATCTGCCCAGAACA |
| Resp18-3-SPL | /5Phos/ATCCTTCCTGCATGGAAAAAAACTCAAGGATCGATCG ATAGCTAAACACTA |
| Resp18-3-SPR | /5Phos/AAGATACTTGGTATAATCGCTAGGAACTCAAAAAAGG GAAACTGCCTTCAT |
| Scg2-1-SPL | /5Phos/AGGGCTTGTTCTCTTTAAAAAACTCAAGAGTCGATCG ATAGCTAAACACTA |
| Scg2-1-SPR | /5Phos/AAGATACTTGGTATAATCGCTGAGAACTCAAAAAATC CAGATTCAAGGCAT |
| Scg2-2-SPL | /5Phos/AGGCAATGTTGTTGGTAAAAAACTCAAGAGTCGATCG ATAGCTAAACACTA |
| Scg2-2-SPR | /5Phos/AAGATACTTGGTATAATCGCTGAGAACTCAAAAAACC CACGACATCTTCAT |
| Scg2-3-SPL | /5Phos/TCTGGTTGGCTCTAGAAAAAAACTCAAGAGTCGATCG |

| | |
|---|---|
| | ATAGCTAAACACTA |
| Scg2-3-SPR | /5Phos/AAGATACTTGGTATAATCGCTGAGAACTCAAAAAACA GGCTACTTTGGGAA |
| Hap1-1-SPL | /5Phos/AGGACACAACGCTTCCAAAAAACTCAAGCTTCGATCG ATAGCTAAACACTA |
| Hap1-1-SPR | /5Phos/AAGATACTTGGTATAATCGCTCTCTACTCAAAAAATCG TGGGTGGTGGATT |
| Hap1-2-SPL | /5Phos/AGCTTGGTGATCTCGGAAAAAACTCAAGCTTCGATCG ATAGCTAAACACTA |
| Hap1-2-SPR | /5Phos/AAGATACTTGGTATAATCGCTCTCTACTCAAAAAACTG ACAACGCTGTTGT |
| Hap1-3-SPL | /5Phos/TAGTGGGTGACAGAGCAAAAAACTCAAGCTTCGATCG ATAGCTAAACACTA |
| Hap1-3-SPR | /5Phos/AAGATACTTGGTATAATCGCTCTCTACTCAAAAAAAG GCACACTGTATCCA |
| Pnmal2-1-SPL | /5Phos/TCTTGCAGAAGGTTCAAAAAAACTCAAGCTTCGATCG ATAGCTAAACACTA |
| Pnmal2-1-SPR | /5Phos/AAGATACTTGGTATAATCGCTGATCACTCAAAAAACA GCTCCTTACACCAA |
| Pnmal2-2-SPL | /5Phos/AGCGCGACTAAATCAGAAAAAACTCAAGCTTCGATCG ATAGCTAAACACTA |
| Pnmal2-2-SPR | /5Phos/AAGATACTTGGTATAATCGCTGATCACTCAAAAAAATC TCTCACAGCCAGT |
| Pnmal2-3-SPL | /5Phos/TATCTTGCGCAGCCAAAAAAAACTCAAGCTTCGATCG ATAGCTAAACACTA |
| Pnmal2-3-SPR | /5Phos/AAGATACTTGGTATAATCGCTGATCACTCAAAAAACTC ACAGCTTCCACCA |
| Ly6h-1-SPL | /5Phos/GGAGGAAGCACACATCAAAAAACTCAAGTCTCGATCG ATAGCTAAACACTA |
| Ly6h-1-SPR | /5Phos/AAGATACTTGGTATAATCGCTCTGAACTCAAAAAAGC TTAACGAAGTCGCA |
| Ly6h-2-SPL | /5Phos/TGGCTGGAATTGGCCAAAAAAACTCAAGTCTCGATCG ATAGCTAAACACTA |
| Ly6h-2-SPR | /5Phos/AAGATACTTGGTATAATCGCTCTGAACTCAAAAAATG CTTCGGAGCGCAA |
| Prkcd-1-SPL | /5Phos/TTGAAGGAGATGCGCAAAAAAACTCAAGAGTCGATCG ATAGCTAAACACTA |
| Prkcd-1-SPR | /5Phos/AAGATACTTGGTATAATCGCTAGCTACTCAAAAAAGC CCAGCTCATAGGAA |
| Prkcd-2-SPL | /5Phos/ACTGCACACACATCAGAAAAAACTCAAGAGTCGATCG ATAGCTAAACACTA |
| Prkcd-2-SPR | /5Phos/AAGATACTTGGTATAATCGCTAGCTACTCAAAAAACC ATCCTCCAGGAAAT |
| Prkcd-3-SPL | /5Phos/ACAGATGAGGTGGGTGAAAAAACTCAAGAGTCGATC |

| | |
|---|---|
| | GATAGCTAAACACTA |
| Prkcd-3-SPR | /5Phos/AAGATACTTGGTATAATCGCTAGCTACTCAAAAAACCT TGGTCTGGAAGGT |
| Synpo2-1-SPL | /5Phos/AGCTCAACCACATGGCAAAAAACTCAAGAGTCGATCG ATAGCTAAACACTA |
| Synpo2-1-SPR | /5Phos/AAGATACTTGGTATAATCGCTCTAGACTCAAAAAATGA GAGGGACAGCTGT |
| Synpo2-2-SPL | /5Phos/TTCTGTTGTGGCACACAAAAAACTCAAGAGTCGATCG ATAGCTAAACACTA |
| Synpo2-2-SPR | /5Phos/AAGATACTTGGTATAATCGCTCTAGACTCAAAAAACA CTCCACTGAAGCCA |
| Synpo2-3-SPL | /5Phos/ACACAGAAGAAGCCTGAAAAAACTCAAGAGTCGATCG ATAGCTAAACACTA |
| Synpo2-3-SPR | /5Phos/AAGATACTTGGTATAATCGCTCTAGACTCAAAAAATAA GCTGGTACCGAGT |
| Plekhg1-1-SPL | /5Phos/TGGCATCCAGCACAACAAAAAACTCAAGTCTCGATCG ATAGCTAAACACTA |
| Plekhg1-1-SPR | /5Phos/AAGATACTTGGTATAATCGCTTCAGACTCAAAAAACG CTGCATCGTGTCTA |
| Plekhg1-2-SPL | /5Phos/TTTGGCCTGAACTGTGAAAAAACTCAAGTCTCGATCG ATAGCTAAACACTA |
| Plekhg1-2-SPR | /5Phos/AAGATACTTGGTATAATCGCTTCAGACTCAAAAAAGTT TGTCTTGCTGGGA |
| Plekhg1-3-SPL | /5Phos/ACTGTAGGGCGAATGCAAAAAACTCAAGTCTCGATCG ATAGCTAAACACTA |
| Plekhg1-3-SPR | /5Phos/AAGATACTTGGTATAATCGCTTCAGACTCAAAAAACT GCCAACTCTCCGTT |
| Ntng1-1-SPL | /5Phos/TGTTAACCTGGAGAGGAAAAAACTCACTAGTCGATCG ATAGCTAAACACTA |
| Ntng1-1-SPR | /5Phos/AAGATACTTGGTATAATCGCTAGAGACTCAAAAAACT CCAAGACAGAGTGA |
| Ntng1-2-SPL | /5Phos/ACTCTTCCGTGCAAATAAAAAACTCACTAGTCGATCG ATAGCTAAACACTA |
| Ntng1-2-SPR | /5Phos/AAGATACTTGGTATAATCGCTAGAGACTCAAAAAAGA GTACCCAGTGGAGT |
| Ntng1-3-SPL | /5Phos/TTTCCTGGCAGACACTAAAAAACTCACTAGTCGATCG ATAGCTAAACACTA |
| Ntng1-3-SPR | /5Phos/AAGATACTTGGTATAATCGCTAGAGACTCAAAAAAGC CGTAGTACCAGGAA |
| Mal-1-SPL | /5Phos/TCCAGGAAGTCTCACCAAAAAACTCAAGCTTCGATCG ATAGCTAAACACTA |
| Mal-1-SPR | /5Phos/AAGATACTTGGTATAATCGCTTCGAACTCAAAAAAGC TGCATCCAGTGTGA |
| Mal-2-SPL | /5Phos/GGGCTTCCAGAACTGAAAAAAACTCAAGCTTCGATCG |

| | |
|---|---|
| | ATAGCTAAACACTA |
| Mal-2-SPR | /5Phos/AAGATACTTGGTATAATCGCTTCGAACTCAAAAAAATT GAGATGGTGGCCA |
| Mal-3-SPL | /5Phos/AGACCGAGAAGCCACTAAAAAACTCAAGCTTCGATCG ATAGCTAAACACTA |
| Mal-3-SPR | /5Phos/AAGATACTTGGTATAATCGCTTCGAACTCAAAAAATCA GGGAAGGTGGTGA |
| Cacng4-1-SPL | /5Phos/TTGTTCTTGCGGCTGTAAAAAACTCAAGAGTCGATCG ATAGCTAAACACTA |
| Cacng4-1-SPR | /5Phos/AAGATACTTGGTATAATCGCTTCTCACTCAAAAAACG CGCTGAGGACAATA |
| Cacng4-2-SPL | /5Phos/AAGGAACTCCCGCTTGAAAAAACTCAAGAGTCGATCG ATAGCTAAACACTA |
| Cacng4-2-SPR | /5Phos/AAGATACTTGGTATAATCGCTTCTCACTCAAAAAAAG GAGGAAGAGGCCTT |
| Cacng4-3-SPL | /5Phos/TGGCTCCGGTGATCTTAAAAAACTCAAGAGTCGATCG ATAGCTAAACACTA |
| Cacng4-3-SPR | /5Phos/AAGATACTTGGTATAATCGCTTCTCACTCAAAAAAAGC TCACCCATGGGAA |
| Cspg5-1-SPL | /5Phos/TCGCACTGCCTGTTTCAAAAAACTCACTAGTCGATCG ATAGCTAAACACTA |
| Cspg5-1-SPR | /5Phos/AAGATACTTGGTATAATCGCTGATCACTCAAAAAAAG CTCTTCAGCCTCGA |
| Cspg5-2-SPL | /5Phos/TGATGGACTCACAGCGAAAAAACTCACTAGTCGATCG ATAGCTAAACACTA |
| Cspg5-2-SPR | /5Phos/AAGATACTTGGTATAATCGCTGATCACTCAAAAAAAC CTGGAAGTCCGTGA |
| Cspg5-3-SPL | /5Phos/AAAGGACTCTTCCTCCAAAAAACTCACTAGTCGATCG ATAGCTAAACACTA |
| Cspg5-3-SPR | /5Phos/AAGATACTTGGTATAATCGCTGATCACTCAAAAAATG GAGTTCTGGATGTT |
| Kcnip3-1-SPL | /5Phos/TGGACATGGGTTCCAAAAAAAACTCACTAGTCGATCG ATAGCTAAACACTA |
| Kcnip3-1-SPR | /5Phos/AAGATACTTGGTATAATCGCTCTCTACTCAAAAAATCA CTGCTGTCTTCCA |
| Kcnip3-2-SPL | /5Phos/TCTCCCTGAGGGAAGAAAAAAACTCACTAGTCGATCG ATAGCTAAACACTA |
| Kcnip3-2-SPR | /5Phos/AAGATACTTGGTATAATCGCTCTCTACTCAAAAAATGC ATAGGTGGTGGCA |
| Csf1r-1-SPL | /5Phos/TTGCTCACACATCGCAAAAAAACTCACTGATCGATCG ATAGCTAAACACTA |
| Csf1r-1-SPR | /5Phos/AAGATACTTGGTATAATCGCTAGTCACTCAAAAAACC ATTCCACACTGCCA |
| Csf1r-2-SPL | /5Phos/ATATGCCAGCGTCTTGAAAAAACTCACTGATCGATCG |

| | |
|---|---|
| | ATAGCTAAACACTA |
| Csf1r-2-SPR | /5Phos/AAGATACTTGGTATAATCGCTAGTCACTCAAAAAACT GGCCACACAAGAAT |
| Csf1r-3-SPL | /5Phos/AACACTGACCTCTGGGAAAAAACTCACTGATCGATCG ATAGCTAAACACTA |
| Csf1r-3-SPR | /5Phos/AAGATACTTGGTATAATCGCTAGTCACTCAAAAAATCA CAGGCATCCATGT |
| Ctss-1-SPL | /5Phos/TATCGTTCATGCCCACAAAAAACTCACTGATCGATCG ATAGCTAAACACTA |
| Ctss-1-SPR | /5Phos/AAGATACTTGGTATAATCGCTCTGAACTCAAAAAATTG GTCATGTCTCCCA |
| Ctss-2-SPL | /5Phos/TCCTCGTCACCAAACGAAAAAACTCACTGATCGATCG ATAGCTAAACACTA |
| Ctss-2-SPR | /5Phos/AAGATACTTGGTATAATCGCTCTGAACTCAAAAAATG CTTCTTTCAGGGCA |
| Bsg-1-SPL | /5Phos/ACCTTGCCACCTCTCAAAAAAACTCAGAAGTCGATCG ATAGCTAAACACTA |
| Bsg-1-SPR | /5Phos/AAGATACTTGGTATAATCGCTTCCTACTCAAAAAAAGT GTCCTCCTGCAGT |
| Bsg-2-SPL | /5Phos/TTGGTGATTGCCTCTTAAAAAACTCAGAAGTCGATCG ATAGCTAAACACTA |
| Bsg-2-SPR | /5Phos/AAGATACTTGGTATAATCGCTTCCTACTCAAAAAAATT GGCTTCAGTGCTA |
| Bsg-3-SPL | /5Phos/TGGTCAGCTGTGACTTAAAAAACTCAGAAGTCGATCG ATAGCTAAACACTA |
| Bsg-3-SPR | /5Phos/AAGATACTTGGTATAATCGCTTCCTACTCAAAAAAACG TCAAGGTTGCTGA |
| Myh9-1-SPL | /5Phos/TTCTTGCTGGAAGGCAAAAAAACTCAGACTTCGATCG ATAGCTAAACACTA |
| Myh9-1-SPR | /5Phos/AAGATACTTGGTATAATCGCTAGTCACTCAAAAAAAG CTGGTTCAAAGCCA |
| Myh9-2-SPL | /5Phos/TTCTTCACCGTCTTGGAAAAAACTCAGACTTCGATCG ATAGCTAAACACTA |
| Myh9-2-SPR | /5Phos/AAGATACTTGGTATAATCGCTAGTCACTCAAAAAATC GAGAGGAGTTGTCA |
| Myh9-3-SPL | /5Phos/TCCACATCCTTCCACAAAAAAACTCAGACTTCGATCG ATAGCTAAACACTA |
| Myh9-3-SPR | /5Phos/AAGATACTTGGTATAATCGCTAGTCACTCAAAAAACAA GCCAATGATCCGA |
| Cyp2f2-1-SPL | /5Phos/TTGGGAGAGGTTTGGGAAAAAACTCACTAGTCGATCG ATAGCTAAACACTA |
| Cyp2f2-1-SPR | /5Phos/AAGATACTTGGTATAATCGCTTCGAACTCAAAAAAAG CAGGTTTCCCAGGA |
| Cyp2f2-2-SPL | /5Phos/TGTTGTACATCTCGCCAAAAAACTCACTAGTCGATCG |

| | |
|---|---|
| | ATAGCTAAACACTA |
| Cyp2f2-2-SPR | /5Phos/AAGATACTTGGTATAATCGCTTCGAACTCAAAAAAAG GACACTTGGGAAGA |
| Cyp2f2-3-SPL | /5Phos/AAGGCATGGATGTACGAAAAAACTCACTAGTCGATCG ATAGCTAAACACTA |
| Cyp2f2-3-SPR | /5Phos/AAGATACTTGGTATAATCGCTTCGAACTCAAAAAAATC ACTGCATCTGTGT |
| Galm-1-SPL | /5Phos/TCTGCTGCAATGGTGAAAAAAACTCACTCTTCGATCG ATAGCTAAACACTA |
| Galm-1-SPR | /5Phos/AAGATACTTGGTATAATCGCTAGCTACTCAAAAAACA CGGGCAAATATGCA |
| Galm-2-SPL | /5Phos/TCCAATTCCGCAAAGCAAAAAACTCACTCTTCGATCG ATAGCTAAACACTA |
| Galm-2-SPR | /5Phos/AAGATACTTGGTATAATCGCTAGCTACTCAAAAAAGC TTCTGGAGGTAACCT |
| Galm-3-SPL | /5Phos/TACTTCCAGGATCCTCCAAAAAACTCACTCTTCGATC GATAGCTAAACACTA |
| Galm-3-SPR | /5Phos/AAGATACTTGGTATAATCGCTAGCTACTCAAAAAACCT GGTTGTGTGGTATA |
| H2-Aa-1-SPL | /5Phos/TTGGCCAAACTCAGGAAAAAAACTCACTCTTCGATCG ATAGCTAAACACTA |
| H2-Aa-1-SPR | /5Phos/AAGATACTTGGTATAATCGCTCTAGACTCAAAAAAGG TCAAAGCTTGCCAA |
| H2-Aa-2-SPL | /5Phos/TGATGAAGATGGTGCCAAAAAACTCACTCTTCGATCG ATAGCTAAACACTA |
| H2-Aa-2-SPR | /5Phos/AAGATACTTGGTATAATCGCTCTAGACTCAAAAAAGAT CGCAGGCCTTGAA |
| H2-Aa-3-SPL | /5Phos/AAGACAGCTTGTGGAAAAAAAACTCACTCTTCGATCG ATAGCTAAACACTA |
| H2-Aa-3-SPR | /5Phos/AAGATACTTGGTATAATCGCTCTAGACTCAAAAAAGG GATGAAGGTGAGAT |
| Rarres2-1-SPL | /5Phos/TCAGCAAGCACTTCATAAAAAACTCAGAGATCGATCG ATAGCTAAACACTA |
| Rarres2-1-SPR | /5Phos/AAGATACTTGGTATAATCGCTAGAGACTCAAAAAACA TAGGGCTAGGGAGA |
| Rarres2-2-SPL | /5Phos/CTTGGAAGGCCAACTGAAAAAACTCAGAGATCGATCG ATAGCTAAACACTA |
| Rarres2-2-SPR | /5Phos/AAGATACTTGGTATAATCGCTAGAGACTCAAAAAACT GTCCACACCGATCT |
| Rarres2-3-SPL | /5Phos/TGGTCTGCTGGAGCTTAAAAAACTCAGAGATCGATCG ATAGCTAAACACTA |
| Rarres2-3-SPR | /5Phos/AAGATACTTGGTATAATCGCTAGAGACTCAAAAAATC CTTCTTGGGGCAGT |
| Gja1-1-SPL | /5Phos/TGATCTGAAGGACCCAAAAAAACTCAGATCTCGATCG |

| | |
|---|---|
| | ATAGCTAAACACTA |
| Gja1-1-SPR | /5Phos/AAGATACTTGGTATAATCGCTGAGAACTCAAAAAAGG CACAGACACGAATA |
| Gja1-2-SPL | /5Phos/ACCCAGGAGGAGACATAAAAAACTCAGATCTCGATCG ATAGCTAAACACTA |
| Gja1-2-SPR | /5Phos/AAGATACTTGGTATAATCGCTGAGAACTCAAAAAACC AGTGACCAGCTTGT |
| Gja1-3-SPL | /5Phos/AGCTAAGGGCTGGAGTAAAAAACTCAGATCTCGATCG ATAGCTAAACACTA |
| Gja1-3-SPR | /5Phos/AAGATACTTGGTATAATCGCTGAGAACTCAAAAAAGT CGCTGATCCACGAT |
| Camk2a-1-SPL | /5Phos/AGAGGGTTAGCAACACAAAAAACTCAAGGATCGATCG ATAGCTAAACACTA |
| Camk2a-1-SPR | /5Phos/AAGATACTTGGTATAATCGCTCTTCACTCAAAAAAGTG GAGGAGAGAAAGT |
| Camk2a-2-SPL | /5Phos/TTCCTTCACACCATCGAAAAAACTCAAGGATCGATCG ATAGCTAAACACTA |
| Camk2a-2-SPR | /5Phos/AAGATACTTGGTATAATCGCTCTTCACTCAAAAAATGG TGCTCTCAGAAGA |
| Camk2a-3-SPL | /5Phos/TTGCTTATGGCTTCGAAAAAAACTCAAGGATCGATCG ATAGCTAAACACTA |
| Camk2a-3-SPR | /5Phos/AAGATACTTGGTATAATCGCTCTTCACTCAAAAAAGG ACTCAAAGTCTCCA |
| Dlg4-1-SPL | /5Phos/TGGAACCCGCCTCTTTAAAAAACTCAAGGATCGATCG ATAGCTAAACACTA |
| Dlg4-1-SPR | /5Phos/AAGATACTTGGTATAATCGCTTCCTACTCAAAAAAACG TAGAGGCGAACGA |
| Dlg4-2-SPL | /5Phos/ATGTGTTCTTCAGGGCAAAAAACTCAAGGATCGATCG ATAGCTAAACACTA |
| Dlg4-2-SPR | /5Phos/AAGATACTTGGTATAATCGCTTCCTACTCAAAAAAAG GTACACAACGTCAT |
| Syn1-1-SPL | /5Phos/TTGGCCATGAAGTTGCAAAAAACTCACTCTTCGATCG ATAGCTAAACACTA |
| Syn1-1-SPR | /5Phos/AAGATACTTGGTATAATCGCTGAGAACTCAAAAAAGT ACCCATTCGGCAGA |
| Syn1-2-SPL | /5Phos/AGTCTCCATTACGTGCAAAAAACTCACTCTTCGATCG ATAGCTAAACACTA |
| Syn1-2-SPR | /5Phos/AAGATACTTGGTATAATCGCTGAGAACTCAAAAAAAT GACCAAACTTCGGT |
| Syn1-3-SPL | /5Phos/TCTGCTCAAGCATAGCAAAAAACTCACTCTTCGATCG ATAGCTAAACACTA |
| Syn1-3-SPR | /5Phos/AAGATACTTGGTATAATCGCTGAGAACTCAAAAAACT GTCAGACATGGCAA |
| Syn2-1-SPL | /5Phos/TTCCGAAGTACCTGCAAAAAAACTCACTGATCGATCG |

| | |
|---|---|
| | ATAGCTAAACACTA |
| Syn2-1-SPR | /5Phos/AAGATACTTGGTATAATCGCTGACTACTCAAAAAAGA CAACCTTTGTGCCA |
| Syn2-2-SPL | /5Phos/TCTGAGCAAACACCCAAAAAAACTCACTGATCGATCG ATAGCTAAACACTA |
| Syn2-2-SPR | /5Phos/AAGATACTTGGTATAATCGCTGACTACTCAAAAAATTG AAGATGGCCACCA |
| Snap25-1-SPL | /5Phos/ACTCTCTTCAACCAGCAAAAAACTCACTGATCGATCG ATAGCTAAACACTA |
| Snap25-1-SPR | /5Phos/AAGATACTTGGTATAATCGCTTCAGACTCAAAAAATGA TGCCAGCATCTTT |
| Snap25-2-SPL | /5Phos/TGAAGCCACCACTGATAAAAAACTCACTGATCGATCG ATAGCTAAACACTA |
| Snap25-2-SPR | /5Phos/AAGATACTTGGTATAATCGCTTCAGACTCAAAAAATTT GTTACCCTGCGGA |
| Snap25-3-SPL | /5Phos/TTGCCCATGTCTAGAGAAAAAACTCACTGATCGATCG ATAGCTAAACACTA |
| Snap25-3-SPR | /5Phos/AAGATACTTGGTATAATCGCTTCAGACTCAAAAAACT GGGTGTCAATCTCA |
| Creb1-1-SPL | /5Phos/TGATGTTGCATGAGCTAAAAAACTCACTTCTCGATCG ATAGCTAAACACTA |
| Creb1-1-SPR | /5Phos/AAGATACTTGGTATAATCGCTAGGAACTCAAAAAATTA CAGTGGGAGCAGA |
| Creb1-2-SPL | /5Phos/TGTACTGCCCACTGCTAAAAAACTCACTTCTCGATCG ATAGCTAAACACTA |
| Creb1-2-SPR | /5Phos/AAGATACTTGGTATAATCGCTAGGAACTCAAAAAACC CTGGGTAATGGCAA |
| Creb1-3-SPL | /5Phos/TCTGCTGTCCATCAGTAAAAAACTCACTTCTCGATCG ATAGCTAAACACTA |
| Creb1-3-SPR | /5Phos/AAGATACTTGGTATAATCGCTAGGAACTCAAAAAATT GCTGGGCACTAGAA |
| Ccl2-1-SPL | /5Phos/TCATCTTGCTGGTGAAAAAAAACTCACTTCTCGATCG ATAGCTAAACACTA |
| Ccl2-1-SPR | /5Phos/AAGATACTTGGTATAATCGCTGAAGACTCAAAAAAAG CCTACTCATTGGGA |
| Ccl2-2-SPL | /5Phos/AGGGCAGATGCAGTTTAAAAAACTCACTTCTCGATCG ATAGCTAAACACTA |
| Ccl2-2-SPR | /5Phos/AAGATACTTGGTATAATCGCTGAAGACTCAAAAAAAG GTGCTGAAGACCTT |
| Ccl2-3-SPL | /5Phos/ACAGAAGTGCTTGAGGAAAAAACTCACTTCTCGATCG ATAGCTAAACACTA |
| Ccl2-3-SPR | /5Phos/AAGATACTTGGTATAATCGCTGAAGACTCAAAAAACA CACTGGTCACTCCT |
| Pycard-1-SPL | /5Phos/TTCAAGAGCGTCCAGGAAAAAACTCACTTCTCGATCG |

| | |
|---|---|
| | ATAGCTAAACACTA |
| Pycard-1-SPR | /5Phos/AAGATACTTGGTATAATCGCTTCCTACTCAAAAAACATCCCCTGACAAGTT |
| Pycard-2-SPL | /5Phos/ACTCTGAGCAGGGACAAAAAAACTCACTTCTCGATCGATAGCTAAACACTA |
| Pycard-2-SPR | /5Phos/AAGATACTTGGTATAATCGCTTCCTACTCAAAAAAGTCCTGTTCTGGCTGT |
| Pycard-3-SPL | /5Phos/ATTTCCTTCAAGGCCTAAAAAACTCACTTCTCGATCGATAGCTAAACACTA |
| Pycard-3-SPR | /5Phos/AAGATACTTGGTATAATCGCTTCCTACTCAAAAAACACCAAGTAGGGATGT |
| Nox4-1-SPL | /5Phos/TCCTCGAAGATAAGCCAAAAAACTCAGAAGTCGATCGATAGCTAAACACTA |
| Nox4-1-SPR | /5Phos/AAGATACTTGGTATAATCGCTAGGAACTCAAAAAATAGGGACCTTCTGTGA |
| Nox4-2-SPL | /5Phos/AAGACTAATGCAGCCAAAAAAACTCAGAAGTCGATCGATAGCTAAACACTA |
| Nox4-2-SPR | /5Phos/AAGATACTTGGTATAATCGCTAGGAACTCAAAAAAGGGATGATGTCTGGTT |
| Nox4-3-SPL | /5Phos/TCCAAATGGACCATCAAAAAAACTCAGAAGTCGATCGATAGCTAAACACTA |
| Nox4-3-SPR | /5Phos/AAGATACTTGGTATAATCGCTAGGAACTCAAAAAAACTCCTCAAATGGGCT |
| Nox2-1-SPL | /5Phos/TTCGTTCAGCTCCATGAAAAAACTCAGAAGTCGATCGATAGCTAAACACTA |
| Nox2-1-SPR | /5Phos/AAGATACTTGGTATAATCGCTCTTCACTCAAAAAAGTCTGTCCACGTACAA |
| Nox2-2-SPL | /5Phos/TTGCTGAGATCGCCAAAAAAAACTCAGAAGTCGATCGATAGCTAAACACTA |
| Nox2-2-SPR | /5Phos/AAGATACTTGGTATAATCGCTCTTCACTCAAAAAATGGTGATGACCACCTT |
| Nox2-3-SPL | /5Phos/TGGGTGCCATTCTAACAAAAAACTCAGAAGTCGATCGATAGCTAAACACTA |
| Nox2-3-SPR | /5Phos/AAGATACTTGGTATAATCGCTCTTCACTCAAAAAACAGAGGTCAGTGTGAA |
| Hmox1-1-SPL | /5Phos/TCACCAGCTTAAAGCCAAAAAACTCAGAAGTCGATCGATAGCTAAACACTA |
| Hmox1-1-SPR | /5Phos/AAGATACTTGGTATAATCGCTGAAGACTCAAAAAATGGTACAAGGAAGCCA |
| Hmox1-2-SPL | /5Phos/TTGTTGCGCTCTATCTAAAAAACTCAGAAGTCGATCGATAGCTAAACACTA |
| Hmox1-2-SPR | /5Phos/AAGATACTTGGTATAATCGCTGAAGACTCAAAAAAATAGACTGGGTTCTGC |
| Hmox1-3-SPL | /5Phos/CCAGAGTGTTCATTCGAAAAAACTCAGAAGTCGATCG |

| | |
|---|---|
| | ATAGCTAAACACTA |
| Hmox1-3-SPR | /5Phos/AAGATACTTGGTATAATCGCTGAAGACTCAAAAAAAC<br>CTCAGGTGTCATCT |
| Sirt3-1-SPL | /5Phos/ACTGCTGAAGGTTGCTAAAAAACTCATCAGTCGATCG<br>ATAGCTAAACACTA |
| Sirt3-1-SPR | /5Phos/AAGATACTTGGTATAATCGCTAGTCACTCAAAAAAGG<br>GTACGGGATGTCAT |
| Sirt3-2-SPL | /5Phos/ACTGCTTCAGACAAGCAAAAAACTCATCAGTCGATCG<br>ATAGCTAAACACTA |
| Sirt3-2-SPR | /5Phos/AAGATACTTGGTATAATCGCTAGTCACTCAAAAAAGG<br>GCACTGATTTCTGT |
| Sirt3-3-SPL | /5Phos/TCCAGCAGTTCTTGTGAAAAAACTCATCAGTCGATCG<br>ATAGCTAAACACTA |
| Sirt3-3-SPR | /5Phos/AAGATACTTGGTATAATCGCTAGTCACTCAAAAAATTC<br>CCGCTGCATAAGA |
| Per1-1-SPL | /5Phos/ATGAGAACTCCGCTGAAAAAAACTCATCAGTCGATCG<br>ATAGCTAAACACTA |
| Per1-1-SPR | /5Phos/AAGATACTTGGTATAATCGCTCTGAACTCAAAAAATG<br>CCAGAAGAGGAACT |
| Per1-2-SPL | /5Phos/AGACATGTCCATGGCAAAAAAACTCATCAGTCGATCG<br>ATAGCTAAACACTA |
| Per1-2-SPR | /5Phos/AAGATACTTGGTATAATCGCTCTGAACTCAAAAAACCT<br>CCAGGGTGTAAGT |
| Per1-3-SPL | /5Phos/TAGACAATCCGGCCTGAAAAAACTCATCAGTCGATCG<br>ATAGCTAAACACTA |
| Per1-3-SPR | /5Phos/AAGATACTTGGTATAATCGCTCTGAACTCAAAAAATG<br>CCTGCTCCGAAATA |
| Per2-1-SPL | /5Phos/TCAGTTCTTTGTGTGCGAAAAAACTCATCCTTCGATC<br>GATAGCTAAACACTA |
| Per2-1-SPR | /5Phos/AAGATACTTGGTATAATCGCTTCCTACTCAAAAAATCC<br>TTCAGGGTCCTTA |
| Per2-2-SPL | /5Phos/AGCAGCTGGTAGTACTAAAAAACTCATCCTTCGATCG<br>ATAGCTAAACACTA |
| Per2-2-SPR | /5Phos/AAGATACTTGGTATAATCGCTTCCTACTCAAAAAAGCT<br>CTCACTGGACATT |
| Per2-3-SPL | /5Phos/AGGGTGTCATGCGGAAAAAAAACTCATCCTTCGATCG<br>ATAGCTAAACACTA |
| Per2-3-SPR | /5Phos/AAGATACTTGGTATAATCGCTTCCTACTCAAAAAATGC<br>ACCTTGACCAGGT |
| Per3-1-SPL | /5Phos/TGGGTCCAGTTGTTCCAAAAAACTCATCGATCGATCG<br>ATAGCTAAACACTA |
| Per3-1-SPR | /5Phos/AAGATACTTGGTATAATCGCTAGCTACTCAAAAAAATA<br>CTGCGAGGCTCTT |
| Per3-2-SPL | /5Phos/TCAGAGGCCGATCTTCAAAAAACTCATCGATCGATCG |

| | |
|---|---|
| | ATAGCTAAACACTA |
| Pe3-2-SPR | /5Phos/AAGATACTTGGTATAATCGCTAGCTACTCAAAAAATG GTGTATGGCAACCA |
| Per3-3-SPL | /5Phos/TGATCTGGCCATGGACAAAAAACTCATCGATCGATCG ATAGCTAAACACTA |
| Per3-3-SPR | /5Phos/AAGATACTTGGTATAATCGCTAGCTACTCAAAAAACC ACTGGCTTCACTGA |
| Sirt1-1-SPL | /5Phos/ACCGTCTCTTGATCTGAAAAAACTCATCGATCGATCG ATAGCTAAACACTA |
| Sirt1-1-SPR | /5Phos/AAGATACTTGGTATAATCGCTCTAGACTCAAAAAACAA GGCGAGCATAGAT |
| Sirt1-2-SPL | /5Phos/TCCTTTGGATTCCTGCAAAAAACTCATCGATCGATCG ATAGCTAAACACTA |
| Sirt1-2-SPR | /5Phos/AAGATACTTGGTATAATCGCTCTAGACTCAAAAAACC ATGACACTGAAGGA |
| Sirt1-3-SPL | /5Phos/AGAGGACAAGACGTCAAAAAAACTCATCGATCGATCG ATAGCTAAACACTA |
| Sirt1-3-SPR | /5Phos/AAGATACTTGGTATAATCGCTCTAGACTCAAAAAATAC TGCCACAGGAACT |
| Npas2-1-SPL | /5Phos/TGGAGCTGAGCTCTTTAAAAAACTCATCTCTCGATCG ATAGCTAAACACTA |
| Npas2-1-SPR | /5Phos/AAGATACTTGGTATAATCGCTGATCACTCAAAAAAGT GTTACCAGGGAGCA |
| Npas2-2-SPL | /5Phos/AATGCCTCCAACATCAAAAAAACTCATCTCTCGATCG ATAGCTAAACACTA |
| Npas2-2-SPR | /5Phos/AAGATACTTGGTATAATCGCTGATCACTCAAAAAAGAT GACGAAGCCATCT |
| Npas2-3-SPL | /5Phos/ACTGTCGGACACATAGAAAAAACTCATCTCTCGATCG ATAGCTAAACACTA |
| Npas2-3-SPR | /5Phos/AAGATACTTGGTATAATCGCTGATCACTCAAAAAACAA GGAGAGGTGTGAT |
| anchor primer-L1 | /5Phos/TCAGGTCATACACTA |
| anchor primer-R1 | AAACATCGTAGACTA |
| anchor primer-L2 | /5Phos/TCGATCGATAGCTAA |
| anchor primer-R2 | CTTGGTATAATCGCT |
| sequencing interrogation probe-R1-GA | /5Phos/GANNACTCA/3Cy3/ |
| sequencing interrogation probe-R1-CT | /5Phos/CTNNACTCA/3TXR/ |
| sequencing interrogation probe-R1-AG | /5Phos/AGNNACTCA/3Cy5/ |
| sequencing interrogation probe-R1-TC | /5Phos/TCNNACTCA/3AF750/ |
| sequencing interrogation | /5Phos/NNGAACTCA/3Cy3/ |

| | |
|---|---|
| probe-R2-GA | |
| sequencing interrogation probe-R2-CT | /5Phos/NNCTACTCA/3TXR/ |
| sequencing interrogation probe-R2-AG | /5Phos/NNAGACTCA/3Cy5/ |
| sequencing interrogation probe-R2-TC | /5Phos/NNTCACTCA/3AF750/ |
| sequencing interrogation probe-L1-GA | /5Cy3/ACTCANNGA |
| sequencing interrogation probe-L1-CT | /5TXR/ACTCANNCT |
| sequencing interrogation probe-L1-AG | /5Cy5/ACTCANNAG |
| sequencing interrogation probe-L1-TC | /5AF750/ACTCANNTC |
| sequencing interrogation probe-L2-GA | /5Cy3/ACTCAGANN |
| sequencing interrogation probe-L2-CT | /5TXR/ACTCACTNN |
| sequencing interrogation probe-L2-AG | /5Cy5/ACTCAAGNN |
| sequencing interrogation probe-L2-TC | /5AF750/ACTCATCNN |

**Table S2.** Oligonucleotide sequences used in RIBOmap.

| Oligonucleotide name | Sequence (5'-3') |
|---|---|
| ACTB_RIBOmap_padlock_1 | /5Phos/AAGATAAACATCGTAGACTAGCAAAGGCGAG GCTCTGTGCTCAGGTCATACACTA |
| ACTB_RIBOmap_padlock_2 | /5Phos/AAGATAAACATCGTAGACTACACCATCACGCC CTGGTGCCTCAGGTCATACACTA |
| ACTB_RIBOmap_padlock_3 | /5Phos/AAGATAAACATCGTAGACTAGGGATAGCACAG CCTGGATAGCAATCAGGTCATACACTA |
| ACTB_RIBOmap_padlock_4 | /5Phos/AAGATAAACATCGTAGACTAGCGCTCGGTGAG GATCTTCATGTCAGGTCATACACTA |
| ACTB_RIBOmap_padlock_5 | /5Phos/AAGATAAACATCGTAGACTAGGCGTACAGGTC TTTGCGGTCAGGTCATACACTA |
| ACTB_RIBOmap_padlock_6 | /5Phos/AAGATAAACATCGTAGACTACTCATCTTGTTTT CTGCGCAAGTCAGGTCATACACTA |
| ACTB_RIBOmap_primer_1 | GGTGTGGACGGGCGGCGGATGTGTCTACGATG |
| ACTB_RIBOmap_primer_2 | TAGGAATCCTTCTGACCCATGCGTGTCTACGATG |
| ACTB_RIBOmap_primer_3 | TGGTACGGCCAGAGGCGTAGTGTCTACGATG |
| ACTB_RIBOmap_primer_4 | CCGTGGTGGTGAAGCTGTAGGTGTCTACGATG |
| ACTB_RIBOmap_primer_5 | GCCGCCAGACAGCACTGTGTGTGTCTACGATG |
| ACTB_RIBOmap_primer_6 | AACAAATAAAGCCATGCCAAGTGTCTACGATG |
| MALAT1_RIBOmap_padlock_1 | /5Phos/AAGATAAACATCGTAGACTACTTCAGACCTTC TGAACCGGTCAGGTCATACACTA |
| MALAT1_RIBOmap_padlock_2 | /5Phos/AAGATAAACATCGTAGACTACGATTATGGATC ATGCCCATCAGGTCATACACTA |
| MALAT1_RIBOmap_padlock_3 | AAGATAAACATCGTAGACTAAATCGGCCTACGTCCCC ATTCAGGTCATACACTA |
| MALAT1_RIBOmap_padlock_4 | AAGATAAACATCGTAGACTAACTTCCTTTGCTCTGCA GTTCAGGTCATACACTA |
| MALAT1_RIBOmap_padlock_5 | AAGATAAACATCGTAGACTAAATTCGATCACCTTCCG CCGCTCAGGTCATACACTA |
| MALAT1_RIBOmap_padlock_6 | AAGATAAACATCGTAGACTACGTAACAGGCCACTGCC AACTTCAGGTCATACACTA |
| MALAT1_RIBOmap_primer_1 | CTGTGTTATGCCTGGTTAGGTATGGTGTCTACGATG |
| MALAT1_RIBOmap_primer_2 | ACACCATCGTTACCTTGAAAGTGTCTACGATG |
| MALAT1_RIBOmap_primer_3 | AGAGAAACCTACAACACCCGGGTGTCTACGATG |
| MALAT1_RIBOmap_primer_4 | AGAAATCCCTTCAGGATCATTAAGGTGTCTACGATG |
| MALAT1_RIBOmap_primer_5 | ACGGAGAACAACTCGCATCACCGTGTCTACGATG |
| MALAT1_RIBOmap_primer_6 | ACCCACCCCACCAATCCCAGTGTCTACGATG |
| Splint_18SRNA_50A_InvdT_1 | ACAAAATAGAACCGCGGTCCTATTCAA AAA AAA AAA AAA AAA AAA AAA AAA AAA AAA AAA AAA AAA AAA AAA AAA TAT CTT TAG T*G*T* /3InvdT/ |
| Splint_18SRNA_50A_InvdT_2 | CATCGTTTATGGTCGGAACTACGACAA AAA AAA AAA AAA AAA AAA AAA AAA AAA AAA AAA AAA AAA |

| | |
|---|---|
| | AAA AAA TAT CTT TAG T*G*T* /3InvdT/ |
| Splint_18SRNA_50A_InvdT_3 | AGGTTTCCCGTGTTGAGTCAAATTAA AAA AAA AAA AAA AAA AAA AAA AAA AAA AAA AAA AAA AAA AAA AAA AAA TAT CTT TAG T*G*T* /3InvdT/ |
| Splint_18SRNA_50A_InvdT_4 | TGTTATTGCTCAATCTCGGGTGGCTAA AAA AAA AAA AAA AAA AAA AAA AAA AAA AAA AAA AAA AAA AAA AAA AAA TAT CTT TAG T*G*T* /3InvdT/ |
| Splint_18SRNA_50A_InvdT_5 | AGATAGTCAAGTTCGACCGTCTTCTAA AAA AAA AAA AAA AAA AAA AAA AAA AAA AAA AAA AAA AAA AAA AAA AAA TAT CTT TAG T*G*T* /3InvdT/ |
| detection probe | /5Cy3/TCATACACTAAAGATAAACA |
